# N-terminal toxin signal peptides efficiently load therapeutics into a natural nano-injection system

**DOI:** 10.1101/2023.11.13.566157

**Authors:** Eva M. Steiner-Rebrova, Rooshanie N. Ejaz, Claudia S. Kielkopf, Mar Pérez Ruiz, Leyre Marín-Arraiza, Ivo A. Hendriks, Jakob Nybo Nissen, Irina Pozdnyakova, Tillmann Pape, Alice Regaiolo, Kira Götz, Ralf Heermann, Simon Rasmussen, Michael Lund Nielsen, Nicholas M. I. Taylor

## Abstract

Targeted delivery of therapeutics to specific cells is a major bottleneck towards personalized medicine. The extracellular injection system (eCIS) of *Serratia entomophila*, the antifeeding prophage (Afp), promises potential for drug delivery purposes. However, the precise mechanism of action, toxin location, and Afp loading remain unclear. Here, we reveal a minimal N-terminal signal peptide (NtSP) of the toxin Afp18, that plays a key role in toxin packing. By engineering fusion proteins, we demonstrate that Afp18’s NtSP can shuttle effectors for Afp loading. We packed non-eCIS effectors, including CRISPR-Cas protein CasΦ-2 from Biggiephage, and a human antimicrobial peptide, LL37, into Afp. Additionally, NtSPs from eCIS effectors of other species facilitate loading of CasΦ-2 into Afp. We observed cargo being packed inside the Afp tail tube through cryo-EM single particle analysis. The presented results enhance our understanding of eCIS toxin packing and contribute to their development as targeted delivery systems.

**Teaser:** A novel use of the Afp nano injection system’s N-terminal signal peptide in targeted therapeutics delivery

## Introduction

Nature has evolved proteinaceous nanoscale injection structures, known as extracellular injection systems (eCIS), to inject effectors into eukaryotic or prokaryotic cells (1). The eCIS family is closely related to the contractile tails of bacteriophages and other related secretion systems e.g., Type VI Secretion Systems (T6SS) (2,3).

Membrane-bound contractile injection systems (CIS), such as the T6SS and Type III Secretion Systems (T3SS), have been engineered to deliver toxins or proteins through host membranes (4–6). Potential drawbacks of these systems are their cell envelope-bound state, therefore a limiting factor for large scale production, the need to generate toxic bacterial strains, and an upper size limit of non-eCIS related proteins for translocation (7). In contrast, eCIS are cell-free protein complexes that transport translated heterologous proteins specifically into eukaryotic or bacterial cells. Unlike viruses, eCIS do not inject genomic DNA or RNA material (1). Thus, the modification of eCIS could lead to non-viral, cell-free nano delivery systems that can deliver active effectors of varying length into target cells in a controlled manner.

One of the best studied eCIS is the antifeeding prophage (Afp). The Afp is encoded on the pADAP plasmid (153kB), harbored by the gram-negative bacterium *Serratia entomophila* (Fig. 1). The purified Afp particle, including a 264 kDa injected toxin, Afp18, causes a rapid anti-feeding effect against larvae (Coleoptera order) of the New Zealand grass grub, *Costelytra giveni*, resulting in larval starvation and mortality (8). *S. entomophila* has been used as a biopesticide for decades (9–11). The overall structure of the antifeeding prophage (Afp) has been determined (12). In the closely related *Photorhabdus* Virulence Cassettes (PVCs), and structurally related T6SS, cargo location, manipulation and/or use of leader or signal sequences, have been investigated (6, 13–15). N-terminal sequences have been proposed to be sufficient for packing of substrates into the PVC particle (13–16). The structures of related CIS have been published recently (12, 17–19), however their mechanism of action, and in particular, cargo packing, and translocation mechanisms remain elusive.

**Figure 1.**
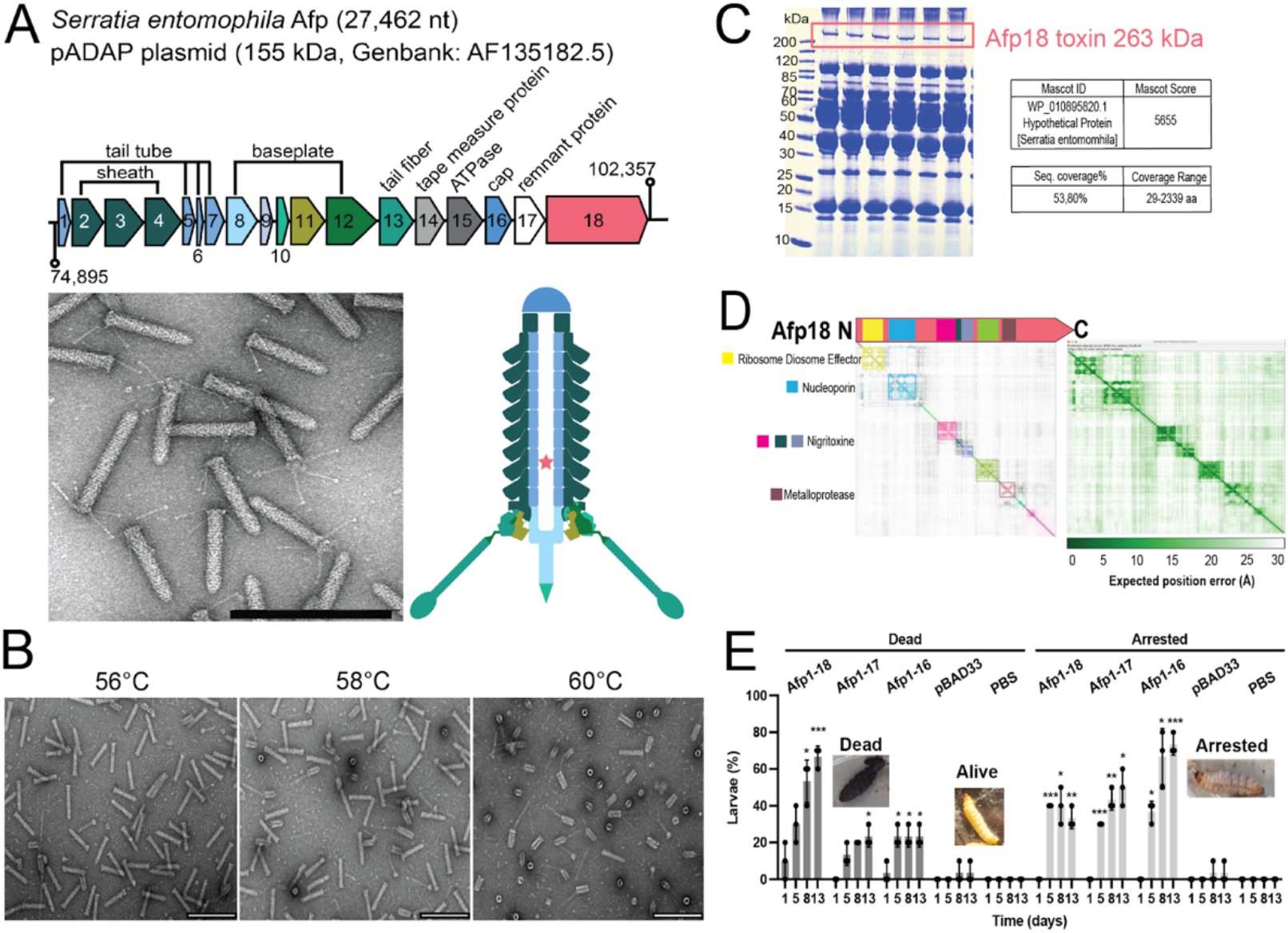
Components and properties of the antifeeding prophage (Afp) (A) Gene cluster organization of the *afp*-encoding region from the *S. entomophila* pADAP plasmid (GenBank: AF135182.5/CP082788.1), ranging from *afp1* (AAT48338/KHA73_24215) to *afp18* (AAT48355/KHA73_24130), gene sizes are scaled. Electron micrograph of negative stained fully assembled, loaded and intact Afp particles at 135,000x magnification showing a range of particle sizes and features e.g., tail fibers, contractile sheath, central spike (scale bar 200 nm). Cartoon representation of Afp particle features and location of putative effector inside the tail tube. (B) Representative electron micrographs as in (A) of Afp revealing high temperature stability (T_stabil_) with observed temperature-induced contraction between 58 - 60°C (particles without Afp18 toxin, Afp1-17, show similar T_stabil_, see Fig. S1.). (C) Coomassie gel of Afp proteins and Afp18 toxin (red box) and result of in-gel digestion and confirmation of toxin presence using LC-MS. (D) Afp18 toxin structure (diameter 110 - 140 Å) prediction using AlphaFold2 (Fig. S5.). Afp18 appears to have a pearl-chain like structure with large number of disordered regions, interspersed by rigid domain cores highlighted with a community clustering approach (https://github.com/tristanic/pae_to_domains) that extracts protein domains from a predicted aligned error (PAE) matrix in ChimeraX (26). For some domains, structural similarity is found using HHPRED and respective functional protein names are indicated (Fig. S6.). (E) Toxicity of Afp particles on *G. mellonella* larvae. Afp particle lysates (Afp1-18, Afp1-17, Afp1-16) were injected into the posterior proleg of *G. mellonella* larvae and the effect was tested over 13 days. The empty pBAD33 vector *E. coli* lysate (pBAD) and PBS were injected as a control (inset image of alive larvae from PBS control). The mortality of larvae can be observed over time and a developmental arrested larvae state was observed (inset images). All particle lysates cause significant mortality and arrested larvae over time, as tested by two-way ANOVA with Dunnett multiple testing compared to pBAD33 control. Shown are the individual numbers from each experiment, mean and standard deviation of three independent experiments (n = 3 experiments, 10 larvae in each treatment).

The eCIS operon encodes a contractile sheath, tail tube, baseplate complex with tail fibers, central spike, a putative tape measure protein, an AAA+-ATPase proposed to be involved in particle assembly or packing of effectors into the particle, a pseudotoxin or toxin remnant (*afp17*) as well as a toxin (*afp18*) at the 3’ end of the cassette (12, 20) (Fig. 1A).

Here, we present a minimal N-terminal signal peptide (NtSP, 20 amino acids) of the toxin Afp18, and reveal conserved physico-chemical properties that can be used to load toxins and effectors into the Afp particle. The NtSP is crucial for stable cargo packing of proteins varying in size and origin, including eCIS and secretion system-related toxins and effectors. Novel, cargo proteins not related to eCIS were loaded into the Afp particle, including a hypercompact CRISPR-Cas system, CasΦ-2, a human antimicrobial peptide, LL37, as well as T3SS effector ExoU and T6SS effector Tse1 (21, 22). Structural data from cryo-EM studies confirms that the cargo is packed inside the Afp tube. We confirmed the presence of the NtSP in the novel toxin chimeras using immuno-detection and mass spectrometry analysis on mature modified Afp particles. We tested in vivo efficacy of cleared lysates of bacterial cells expressing Afp particles with native toxins (Afp18 and Afp17) and without, as well as Afps containing toxin chimeras with anti-eukaryotic effectors ExoU and LL37 on *G. mellonella* larvae, and observed high larval mortality but both in presence and absence of Afp particles. Additionally, we show that NtSPs from other species with similar physico-chemical properties can load CasΦ-2 into Afp, and mutational analysis of Afp18 NtSP’s reveals that substantial mutation of hydrophilicity (down to 45%) does not abolish cargo packing capacity.

The Afp particle demonstrates high long-term stability, and an efficient method of loading a variety of eCIS and non-eCIS related cargo, making it a prime candidate for development as a biotechnological tool for targeted drug delivery. Understanding the Afp mechanism of action offers significant potential for agricultural pest control, and for use as a protein or therapy delivery tool. Modification of eCIS could lead to non-viral, cell-free nano delivery systems, capable of delivering cargos of varying length and effect, into assorted target cells in a controlled manner.

## Results

### Co-production of thermostable Afp syringe–cargo particles

We cloned the *afp*-encoding region from *S. entomophila* and overexpressed Afp in *Escherichia coli* cells (Fig. 1A). The thermostability of Afp was analyzed, as it could be an important factor in potential applications. We exposed Afp particles Afp1-18 and without Afp18 (Afp1-17) for 10 min at increasing temperatures and screened for intact particle morphology using negative staining electron microscopy. Afp appears to be very temperature stable (Fig. 1B) with a temperature stability (T_stabil_) of around 58°C for a fully loaded Afp particle (Afp1-18), as well as when Afp18 is not present (Afp1-17) (Fig. S1.). The Afp particle can be produced in *E. coli* and appears stable at 4°C for extended time frames (>2 years) (Fig. S2&S13.). All proteins encoded in the *afp* operon were detected by in solution liquid chromatography and mass spectrometry (LC/MS) experiments, except for Afp17, proposed to be an inactive toxin remnant (15) (Fig. S3.). The presence of the Afp18 toxin was confirmed by in-gel digest and LC/MS of a final particle preparation used for cryo-EM with full sequence coverage (Afp18 peptide coverage starting at amino acid 30) (Fig. 1C, Fig. S3&4). Afp18 is a large 264 kDa toxin and we predicted its structure using AlphaFold2 (23, 24), however, prediction accuracy is low. Together with the presence of several unstructured regions and its rather large diameter (100 - 140 Å), the structure prediction argues against a single, folded structure (Fig. 1D, Fig. S5.). Using structural similarity search tools (HHPRED), several Afp18 domains show structural homology to published structures, including nigritoxin (100% identity), a bacterial toxin against crustaceans and insects (Fig. 1D, Fig. S6).

*S. entomophila* and purified Afp particles cause the highly host-specific amber disease, cessation of feeding and larval mortality in New Zealand grass grub, *Costelytra giveni* (10, 11). Due to the lack of *C. giveni* larvae to test Afp particle activity, we tested killing potency and the effect of Afp particles on *Galleria mellonella* larvae, prompted by the successful and rapid killing efficacy of a closely related PVC from *Photorhabdus asymbiotica* (PaATCC43949 PVCpnf, which induces rapid melanisation and death of larvae within 30 min) (16). Heterologously produced Afp particles (Afp1-18), Afp lacking its large Afp18 toxin (Afp1-17) and Afp lacking both toxins (Afp1-16) were overexpressed in *E. coli,* and the cleared non-purified cell lysates were injected into *G. mellonella* and larval development observed over 13 days. Afp1-18 lysates cause the highest killing of larvae (Fig. 1E), resulting in dark larval color change, no response upon pinch stress and no butterfly development, suggesting a killing effect of the Afp18 toxin, although we cannot exclude that killing is established in an Afp1-17-independent fashion. The remaining larvae that were not killed showed a novel phenotype. All tested Afp particles (Afp1-18, Afp1-17 and Afp1-16) caused larvae to stop developing into mature butterflies, while being responsive to pinch stress, here termed arrested larvae. The highest number of arrested larvae could be observed when treated with Afp1-16 lysates (Fig. 1E right). Overall, all lysates lead to significantly more dead and arrested larvae compared to the pBAD33 control. Afp17 is a predicted remnant toxin and bioinformatic analysis (HHPRED, BlastP) suggests structural homology to a two component-system histidine-kinase (KdpD, signaling protein) or ADP-ribosylation capabilities, which are used as bacterial adaptation strategies or counteract host defenses (25). The results indicate the Afp could be potentially targeting a broader range of arthropods than only Coleoptera (*Costelytra giveni*, beetles).

### Stable C-terminal Afp18 toxin co-produced with Afp particle

We did not observe any differences in assembly or morphology of Afp particles produced with or without toxin cargo or with truncated Afp18 toxin variants, and toxin levels can be detected by Coomassie staining and in immuno-detection (Fig. 2A,B and C, Fig. S1).

**Figure 2.**
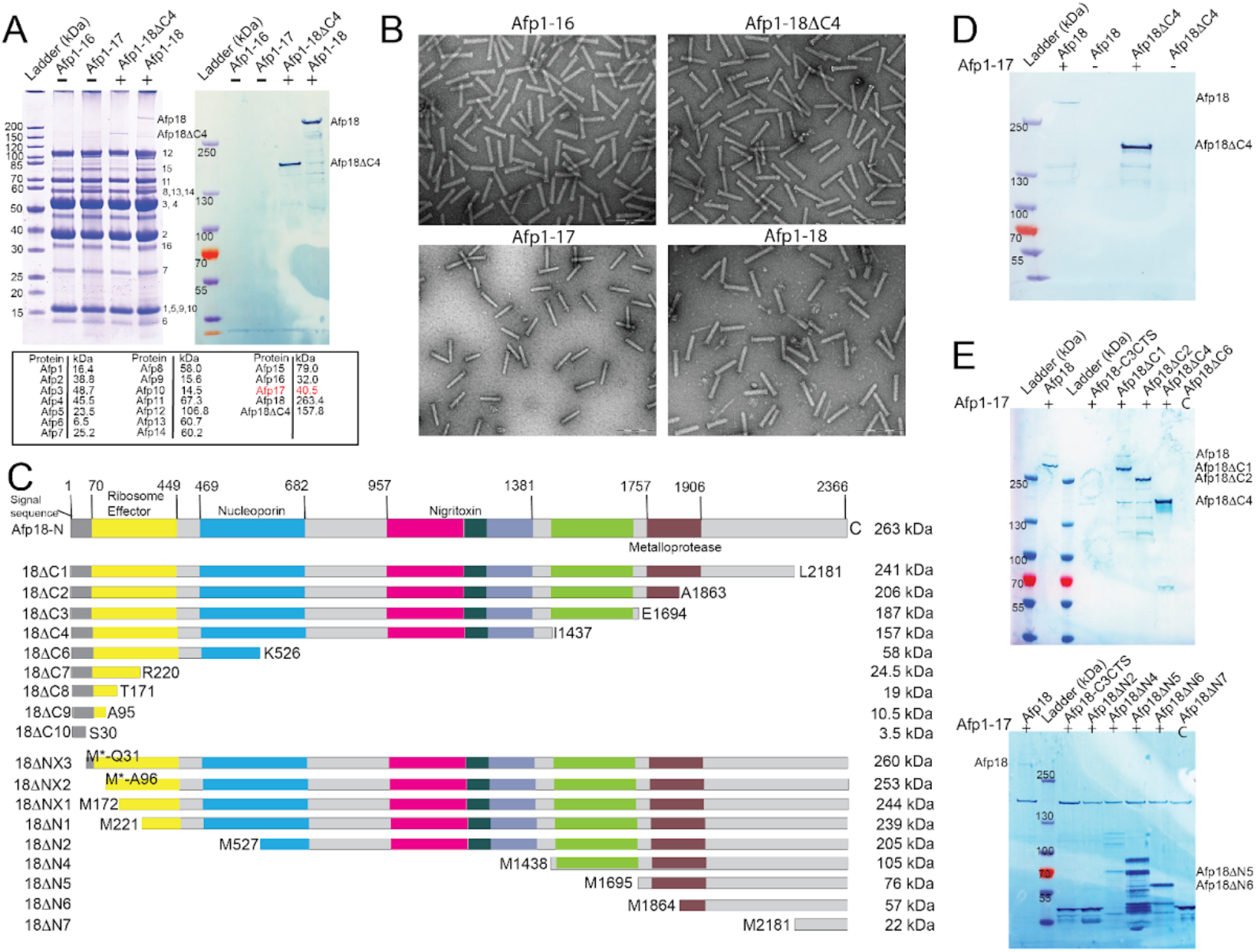
Truncation of Afp18 toxin shows stable toxin purification and particle assembly. **(A)** Left: Purified Afp particles using one plasmid-based expression (pBAD33 only); Afp1-18, Afp1-18ΔC4 (N-terminal half of Afp18), Afp1-17 and Afp1-16 stained with Coomassie (left). Right: immuno-detection of Afp18 in these samples. **(B)** Afp variants investigated for integrity and features using negative stain electron microscopy (scale bar 200 nm). **(C)** Systematic Afp18 truncation from N-terminal (ΔN-constructs) and C-terminal (ΔC-constructs) sites. At the Afp18 N-terminus, no signal sequence could be detected using state-of-the art detection programs (*29*, SignalP 6.0) around residue area 1-70 amino acids. **(D)** Immuno-detection blot of toxin, Afp18 and half truncated Afp18ΔC4 co-produced with Afp (Afp1-17). The toxin was produced in parallel without Afp to verify toxin co-purification with Afp particles (-Afp18/-Afp18ΔC4). **(E)** Immuno-detection blot visualizing co-production of C-terminal and N-terminal truncated Afp18. C-terminal truncations result in more stable toxin co-purification (top), where Afp1-17+Afp18 serves as co-expression control, and Afp18ΔC6 and Afp18ΔN7 constructs serve as antibody controls since they are outside of antibody recognition site (C). Toxin degradation profiles in the form of ladder-like bands (e.g., Afp18ΔN4, Afp18ΔN5) can be seen for N-terminal truncated Afp18 variants (bottom).

To investigate whether it is possible to co-express the cargo and Afp on a separate plasmid, co-expression of Afp particles (Afp1-17) and Afp18 on separate plasmids was tested. The Afp18 toxin can be successfully co-expressed and packed into the Afp particles, with comparable levels of co-purified toxin compared to the one-plasmid production approach (Fig. 2D). The two-plasmid approach makes exchange of Afp particle cargo quicker and more flexible, since cloning of large (>20 kb) plasmids can be challenging. Furthermore, for the co-production (Fig. 2&3), we developed a short particle purification protocol for high throughput screening of Afp18-manipulated variants (Materials and Methods). As a control, toxin and effectors were expressed without Afp particles to monitor soluble aggregation and to prove that particle-cargo co-production is successful, from here on called mock expression (Fig. 2D). The Afp18 toxin and Afp18ΔC4 (about half the size of wild type Afp18) can be co-produced in both the one plasmid and two-plasmid expression and particles show the same architecture (Fig. 2A&D). This indicates that the N-terminus is important for toxin packing and that both particle-toxin production protocols can be used.

N-terminal regions of PVC toxins were recently found to have a similar signal sequence (13–15), and it has been proposed that leader sequences can also be positioned along the whole protein toxin cargo (14). To examine what part of Afp18 toxin is required for packing, we designed N- and C-terminal Afp18 truncations (ΔN and ΔC), aided by domain detection (Fig. 2C&E) and secondary structure predictions (27, 28) using the two-plasmid expression approach.

The Afp18 toxin can be C-terminally truncated down from 264 kDa to a detectable minimum of 10.5 kDa (Fig. 2E, Fig. S7.). The C-terminally truncated Afp18 variants were stably co-produced until truncation Afp18ΔC6 (58 kDa) with negative mock expression, smaller C-terminal variants showed positive mock purification, indicating toxin aggregation or protein solubilization (Fig. 2E, Fig. S8.). N-terminal truncation of Afp18 results in toxin degradation, however Afp particle morphology is not affected (Fig. 2E, Fig. S9&10.). It appears that Afp18 can be more easily manipulated on its C-terminus and that manipulation of the cargo does not affect or impair Afp architecture (Fig. 2B, Fig. S10-S12.). As an additional indicator that the N-terminus plays a crucial role, all N-terminal Afp18 truncations resulted in pronounced toxin degradation (Fig. 2E).

The results strongly suggest a crucial region located at the Afp18 N-terminus, an N-terminal signal peptide (NtSP), for Afp cargo allocation and C-terminal attachment of novel toxins and effectors (Fig. 2A, D&E).

### Use of Afp18 N-terminal region as toxin and cargo delivery scaffold

We wanted to investigate if effector loading is conserved and if the Afp18 N-terminal region could function as a scaffold to attach other toxins and effectors, so called toxin-chimeras, of differing sizes and origins (Fig. 3). The particle and cargo were produced in a two-plasmid co-expression approach (Fig. 3A). Interestingly, *Yr*Afp17, and *P. luminescens* eCIS effector CyaA (PluDJC_08830) were co-purified without any Afp18 N-terminus attached, suggesting similar NtSP domains are present in these proteins (*30*) (Fig. 3B, Fig. S13.). The T6SS effector, Tse1, and T3SS effector ExoU from *P. aeruginosa* were both successfully loaded as Afp18 toxin-chimera, Afp18ΔC8-Tse1 and Afp18ΔC8-ExoU maintaining high temperature stability (31, 32) (Fig. 3C, Fig. S14.). As a control, the non-eCIS related toxin-chimeras were produced in parallel (mock expression) without Afp particle, to exclude false positive results through soluble toxin-chimera aggregates (Fig. 3C, Fig. S14.). The largest manipulated cargo tested was Afp18-sfGFP with a total size of 290.8 kDa (Table 1, Fig. S15.), indicating that cargo payload could be increased, at least to some extent, in molecular weight. Attachment of sfGFP to smaller Afp18ΔC4 truncation variant seems to enhance toxin solubility and results in a positive result in mock expression (Fig. S16.). However, for the full length Afp18-sfGFP no detectable soluble amounts without Afp could be produced (Fig. S15.). Interestingly, toxin *PAU_RS10120*, sharing structural homology with ABC toxins with RHS (rearrangement hot-spot) repeat toxins (detected using HHPRED), was expressed but not packed into the particle (Fig. S17.).

**Figure 3.**
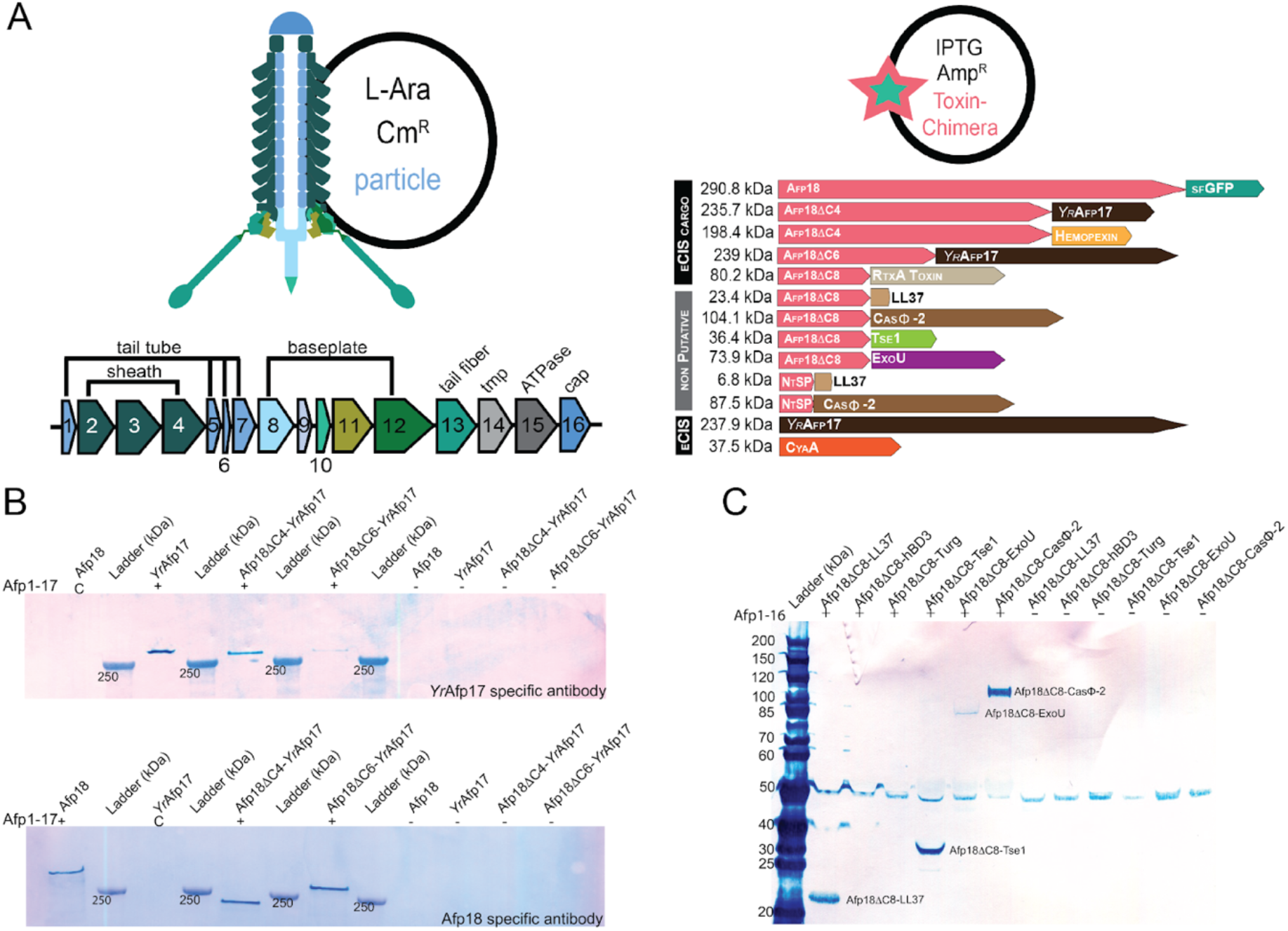
Afp particle and toxin-chimera co-expression results in successful attachment of modified cargo. **(A)** Schematic of *afp* gene cluster (a*fp1-16*) and toxin and toxin-chimera genes designed, selected to explore various types and sizes and origins of cargo that can be attached. The Afp particle is produced on a pBAD33 (Cm^R^) L-arabinose inducible vector and for quick cargo exchange, all cargos are co-expressed on a pET11a (Amp^R^) IPTG inducible vector. Packing of eCIS toxins (black bar) and non eCIS related cargos (grey bar), in the form of toxin-chimeras, are investigated. **(B)** Immuno-detection of successfully attached toxins and toxin-toxin chimeras into Afp1-17 compared to wild type Afp particle Afp1-17+Afp18 co-expression. **(C)** Immuno-detection of successfully Afp18ΔC8-effector chimeras with successfully attached T6SS and T3SS Secretion System effectors from *Pseudomonas aeruginosa*, ExoU and Tse1 and two eCIS unrelated cargos, Biggiephage hypercompact CRISPR-Cas protein CasΦ-2 and the human antimicrobial peptide, LL37. Other antimicrobial peptides, Human beta-defensin 3 (hBD3) and Turgencin (Turg), did not associate with the particle.

**Table 1.**
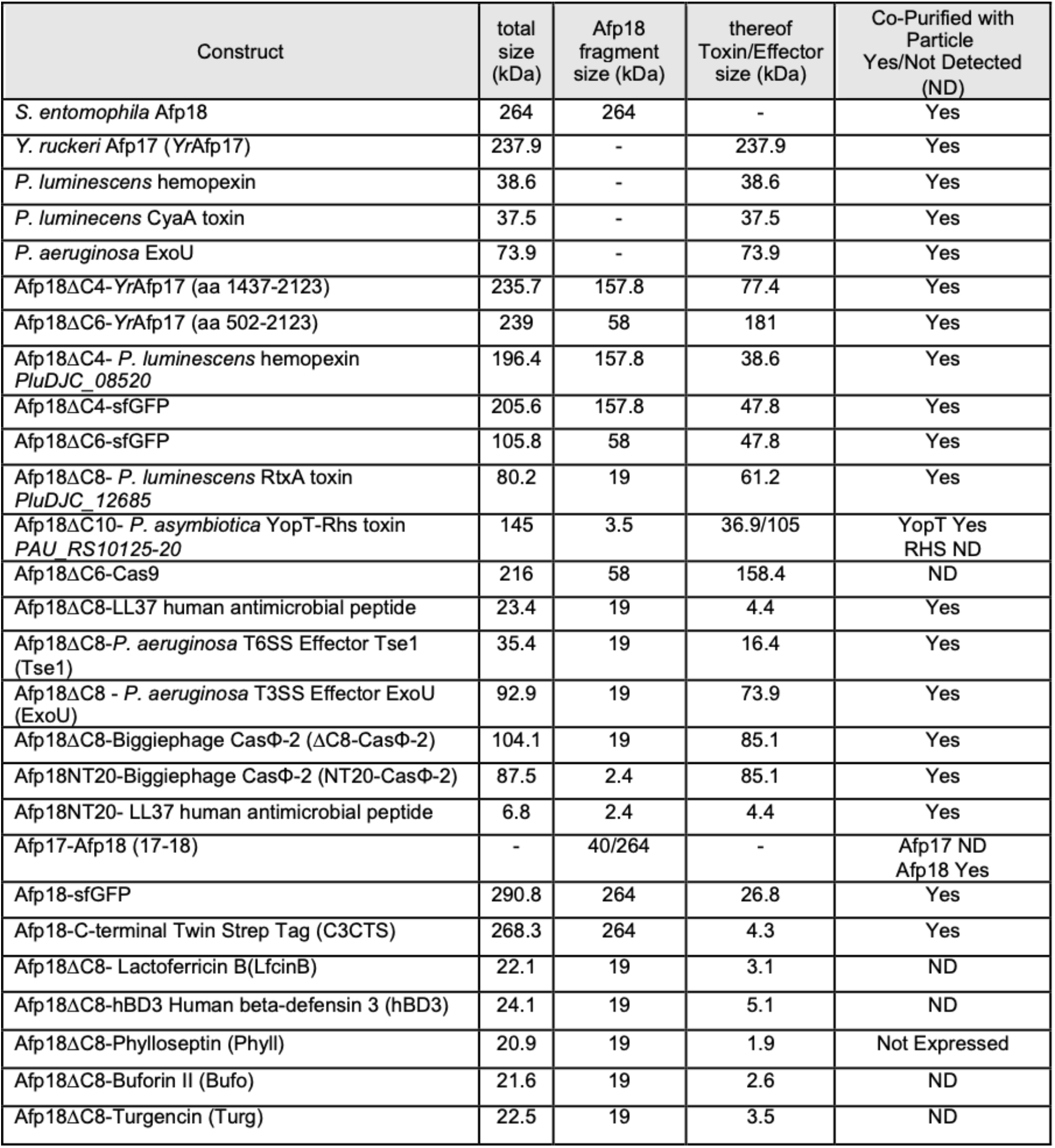
Overview of toxin and toxin-chimera proteins and results of co-purification with Afp. **(A)** *Y. ruckeri Yr*Afp17, *P. luminescens* hemopexin, CyaA toxin and *P. aeruginosa* T3SS effector ExoU effector, were co-purified without Afp18 as scaffold, Rhs toxin and Afp17 remnant toxin are two examples of not detectable (ND) toxin co-purification with the particle.

### Exploring non-eCIS related cargo for drug delivery purposes

To explore structural and biophysical limits of possible cargo, we pursued the attachment of antimicrobial peptides (AMPs). Cationic AMPs are a powerful tool to disrupt a broad range of membranes. Because of their small size, theoretically it would be possible to pack high amounts into the Afp particles. We selected AMPs with different (predicted or experimentally verified) structural features, including Cathelicidins LL37 hCAP18 (33) (LL37, α-helical), Lactoferricin B (34) (LfcinB, β-sheet), human beta-Defensin-3 (hBD3, mixed secondary structure), Phylloseptin (36) (α-helical), Buforin II (37) (helical-helix-propeller structure) and Turgencin (38) (α-helical). Apart from Phylloseptin, all other peptides could be expressed as C-terminal fusions to Afp18 NtSP (Fig. S17.). Out of this selection, only the human antimicrobial peptide LL37, attached to Afp18ΔC8, could be detected to be loaded into the Afp particle (Fig. 3C, Fig. S14.). Structural comparison using AlphaFold2 prediction and available structures of AMPs showed that the Afp18 NtSP is unstructured for all predictions and accessible (Fig. S18.). We observe a high number of cysteine residues for the candidates that were not purified along with the Afp particle. Resulting disulfide bridges and secondary structure differences could therefore possibly be a limiting factor for Afp cargo allocation. Alternatively, NtSP is shorter than the minimal packing sequence and actually depends on downstream structural features that are present in some but not all of the tried cargos to establish their efficient packing.

As a second, eCIS-unrelated cargo group, we pursued packing CRISPR-Cas gene editing enzymes into Afp, as cell-specific targeted delivery will lead to minimize off-target effects and more efficient gene editing. Packing of Cas9 from *Francisella novicida* and a hypercompact Biggiephage Casj12 (CasΦ-2) to Afp18ΔC6 and Afp18ΔC8 were attempted, respectively (22, 39). The hypercompact CasΦ-2 was chosen for its small size in case there is a limit to cargo size for packing. For Afp18ΔC6-Cas9, we did not see conclusive attachment, however, Afp18ΔC8-CasΦ-2 was clearly copurified with the Afp particle (Fig. 3C). Purified particles appeared fully formed and complete in architecture, with minimal amounts of incompletely assembled baseplates and high temperature stability (Fig. S14&S19.).

All Afp particle components (except Afp17), Afp18ΔC8-CasΦ-2/ExoU/Tse1 and LL37 effectors could be detected by immuno-detection and in solution mass spectrometry (LC/MS) (Fig. 4D, Fig. S20.).

**Figure 4.**
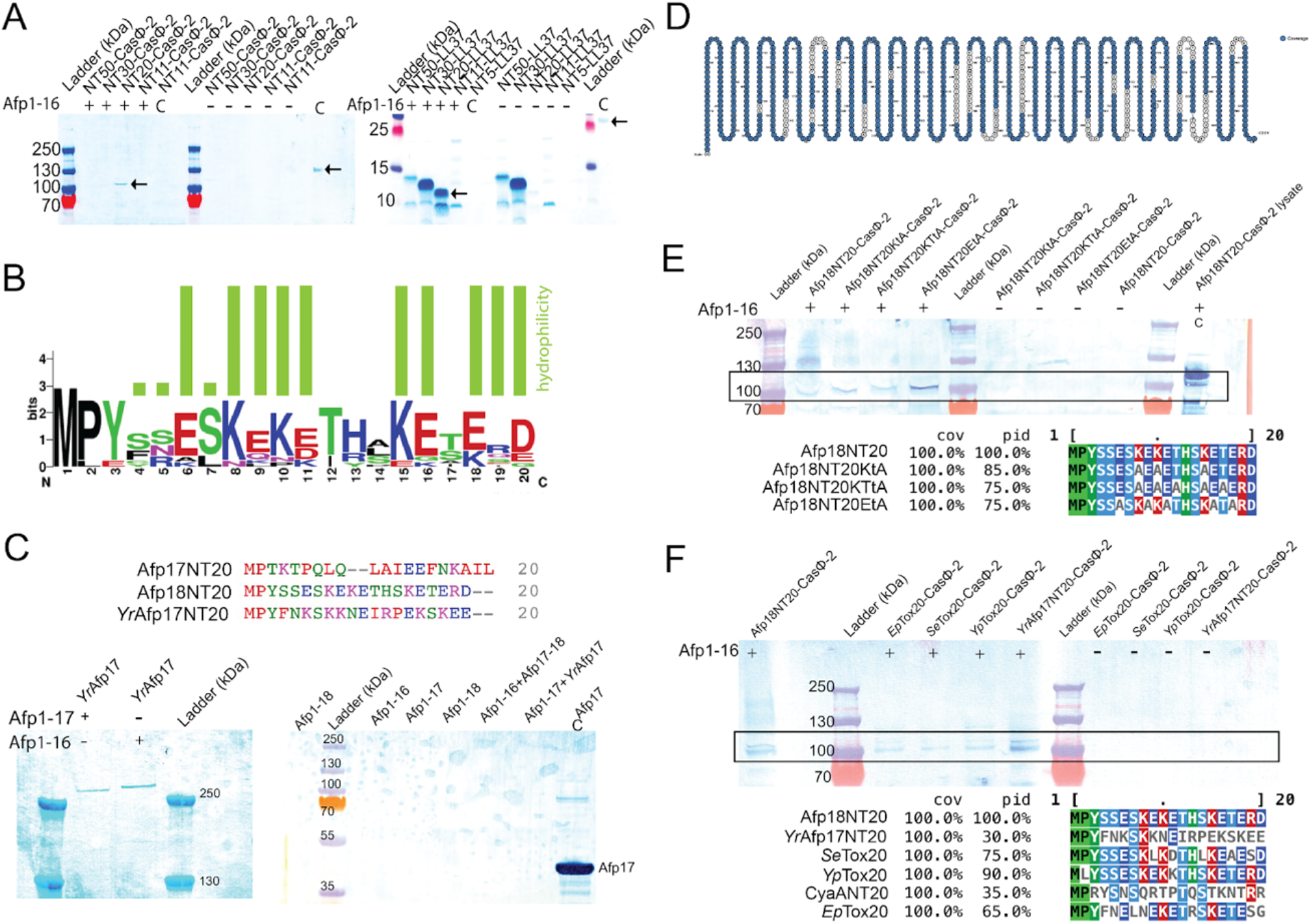
Characterization of NtSPs. **(A)** Truncation series of first 50 N-terminal (NT) amino acids of Afp18 and attachment of non-eCIS related cargo, CasΦ-2 and LL37. The Afp18NT20-effector chimeras, Afp18NT20-CasΦ-2 and Afp18NT20-LL37 all show successful Afp packing and no effector aggregation/solubilization in the control. **(B)** Afp18NT20 like N-terminal packing domains (NtSPs) found in *Erwinia*, *Yersinia*, *Salmonella* and related *Serratia* species and aligned against *Yr*Afp17 and Afp18NT20 termini using Clustal Omega. All effectors including homologous NtSPs are located downstream of an eCIS like particle. No conserved residues can be highlighted, however, there is an obvious presence of polar amino acids, visualized using WebLogo (https://weblogo.berkeley.edu/). Highly hydrophilic residue occurrence is indicated with green bars. **(C)** Comparison of two NtSPs that showed cargo packing, Afp18NT20 and *Yr*Afp17NT20, and one cargo that was not packed, Afp17NT20. Successfully packed cargo has NtSPs with high polar amino acid content and alignment-independent ACC calculations, using cross correlation and the Hellberg z-scale (42) to highlight NtSP packing domain characteristics. Immuno-detection blots confirm *Yr*Afp17 (YR17) packing without any Afp18 fragment and that Afp17 cannot be attached to the Afp particle but can be expressed and detected in expression lysates (control Afp17). **(D)** Mass spectrometry analysis confirming peptide coverage and presence of Afp18NT20 and CasΦ-2 in particle preparations. **(E)** Mutational analysis of Afp18NT20 and mutation of up to 5 hydrophilic amino acids did not abolish NtSP packing function, tested using immuno-detection blotting. Multiple sequence alignment of NtSPs highlighting protein identity (pid) and alanine substitutions using ClustalOmega and MView (43). **(F)** NtSPs from other species can pack CasΦ-2 into Afp, tested using immuno-detection blotting. The sequence alignment and analysis shows high and low protein identity.

For *in vivo* efficacy tests on *G. mellonella* larvae, we injected three toxin-chimera particles (Afp18ΔC8 -*P. aeruginosa* T3SS effector ExoU (PAExoU), Afp18ΔC8-LL37 human antimicrobial peptide and Afp18NT20-LL37) as well as Afp particles (Afp1-18, Afp1-17 and Afp1-16) (Fig. 3D). ExoU is an intracellular phospholipase targeting cell membranes (40) and has previously been used in similar experiments for the PVCs (13). Although we could observe a significant increase in killing of *G. mellonella* larvae (*Lepidoptera*, moths & butterflies) in SE1-16+Afp18ΔC8ExoU and SE1-16+Afp18NT20-LL37 compared to pBAD33 control already after 2 and 3 days, respectively (Fig. 3D) we observed a similar lethality without Afp present (Fig. S21). This suggests that the main killing effect in the coexpression tests could come from the excess toxins, which is inherent to our experimental setup. We acknowledge limitations to the experiment and the native toxicity of insect toxins, currently we have no means for normalization of toxin expression levels without Afp to toxin-loaded Afp particles (ratio toxin:Afp).

### The Afp18 N-terminal signal peptide (NtSP)

The results described above indicate conserved N-terminal sequence properties that are crucial for Afp loading. We further investigated what the minimal Afp18 packing sequence is, by creating shorter N-terminal truncations of the first 50 amino acids (50, 30, 20, 11 and 5 amino acids, NT50-NT5), and attachment of CasΦ-2 and LL37 as two non-eCIS related candidates, for better toxin detection (minimum 10 kDa in size) (Fig. 4A). For both Afp18-effector truncation series, Afp18NT20 (first 20 amino acids) fusions, Afp18NT20-CasΦ-2 and Afp18NT20-LL37 showed successful particle packing. A high abundance of polar amino acids is evident, among the first 20 amino acids of Afp18 and in other Afp18-like N-terminal sequences, however, conventional multiple sequence alignments did not pinpoint a consensus sequence (Fig. 4B&C). Our minimal tested and functional N-terminal signal peptide comprising 20 amino acids, will be referred to as Afp18NT20.

### Auto Cross Correlation and Covariance (CC) describing NtSP properties

Common effector properties were therefore investigated with alignment independent approaches. We investigated successfully loaded effectors using the cross covariance (CC) of amino acids in N-terminal regions using the Hellberg z-scale (42). The N-termini of Afp18, *Yr*Afp17 as well as other putatively packed eCIS cargo show highly negative values when investigating the z1 and z3 scale parameters (as related to hydrophilicity (z1), and electronic properties (z3)) and a >60% content of polar amino acids (R, D, E, H, K, S, T, Y) (Table 2, Fig. 4C). In contrast, Afp17 which is not packed shows highly positive CC value and low polar amino acid content.

**Table 2.**
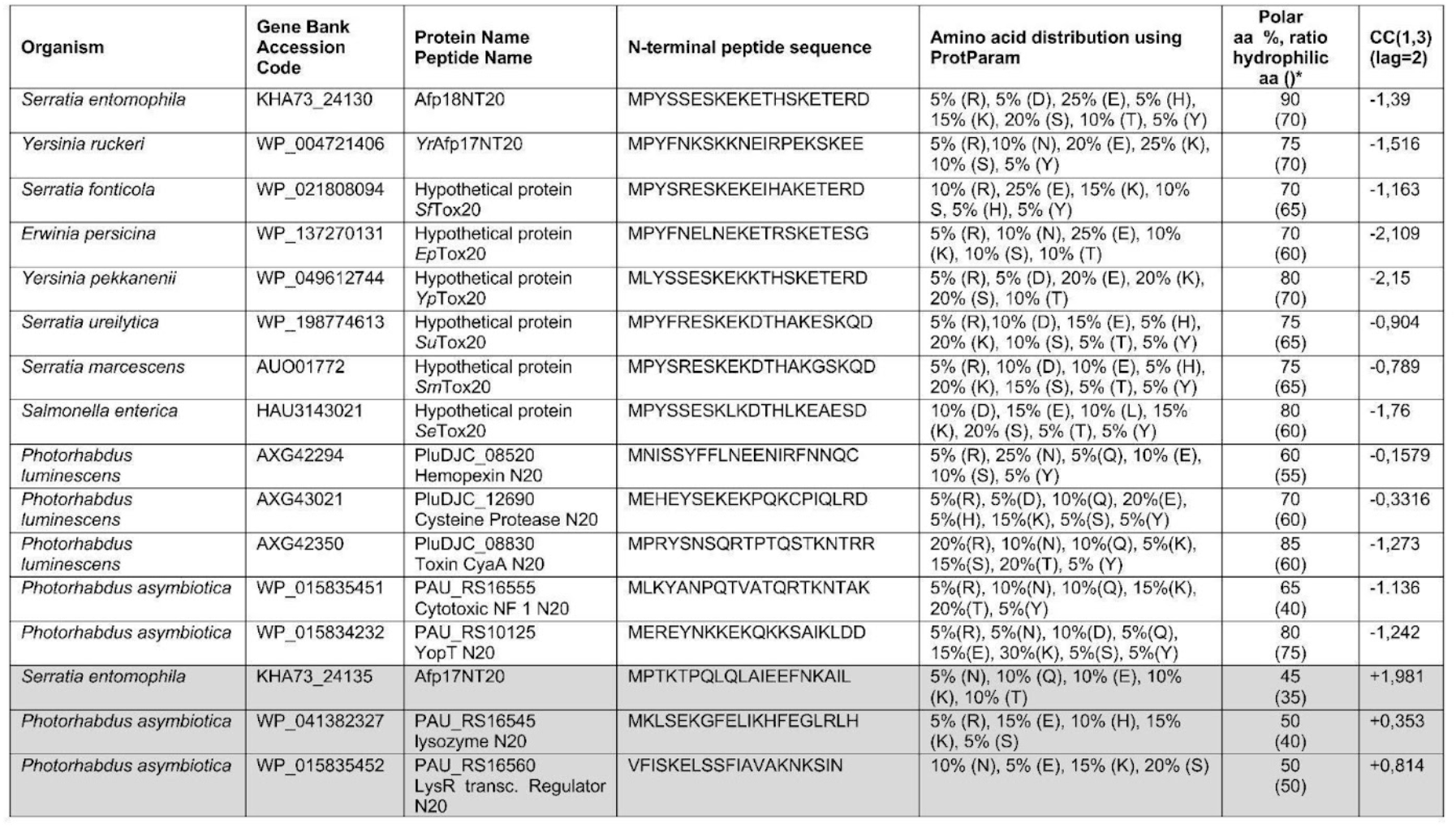
Analysis of NtSPs of effectors in other eCIS. Homology search using Afp18NT20 as search input revealed related effectors and eCIS particles in the species, *Serratia, Yersinia, Erwinia* and *Salmonella*. Investigation of NtSPs of known eCIS effectors in *P. luminescens* and *P. asymbiotica* show similar high polar amino acid content and negative CC values. Afp17NT20 is an example of an experimentally proven effector that was not associated with the Afp particle and NtSP that can not pack effectors (* calculated with peptide calculator https://www.bachem.com/knowledge-center/peptide-calculator/).

We searched for Afp18NT20 peptide homologs using the BlastP® suite (41), manually investigated each hit for presence of eCIS genetic elements upstream of the operon and found more eCIS particles with Afp18-like NtSPs (Table 2, Fig. 4B). When calculating the CC values for other eCIS related effectors at the N-termini, we find that they share similar physical chemical properties and a high percentage of polar amino acids. Similar N-terminal properties can also be highlighted for other eCIS effectors (Table 2).

### Mutational Analysis of Afp18 NtSP

Jiang *et al.* 2022 showed that point mutations within the N-terminal packing sequence of PVCs do not influence successful packing (13). We hypothesized that NtSP’s that have a high abundance of hydrophilic residues (Table 2, absence of hydrophobic patches) are quite resilient to mutations in particular residue positions, but rather depend on an overall hydrophilic (ζ, zeta for hydrophilic amino acids) signal peptide property. Therefore, mutational analysis of Afp18NT20 was carried out by deleting three lysine residues (ζ of 55%), lysine and threonine mutant (ζ of 55%), and a glutamic acid mutant (ζ of 45%) fused to non-eCIS cargo CasΦ-2, to investigate if specific amino acid patterns are required and if lower hydrophilicity abolishes packing capacity. Nevertheless, all mutation variants still packed CasΦ-2 confirmed by negative staining, immuno-detection blotting and mass spectrometry analysis (Fig. 4E, Fig. S22&S23.), suggesting that these mutations are not sufficient to block loading into the particle. (Mutated) NtSPs were confirmed as being present in particle preparations (Fig. S24) by immune-detection, in solution mass spectrometry and particle integrity confirmed over negative staining electron microscopy. As a negative control N-terminal sequences with low hydrophilic and hydrophobic patches, Afp17NT20 (ζ of 35%), ExoUNT20 (ζ of 40%), were shown to not pack CasΦ-2 (Fig. S25.).

### NtSPs of other species successfully pack CasΦ-2 into Afp

We wanted to investigate if NtSPs of different species can pack non eCIS related cargo into Afp. We performed a homology search (Table 2), and chose NtSPs which are also located close to gene clusters putatively encoding eCIS syringes (Table S1.). We then evaluated whether these sequences, *Se*Tox20, *Yp*Tox20, CyaANT20, *Ep*Tox20, *Yr*17NT20 can pack CasΦ-2. All NtSPs showed successful packing of CasΦ-2 into Afp, confirmed by immune-detection blotting and negative staining EM (Fig. 4F, Fig. S26.).

Mass spectrometry analysis confirmed particle components, NtSPs and cargo presence (Fig. S22&24.) Prediction by AlphaFold2 (23, 24) of NtSP-CasΦ-2 chimeras reveal CasΦ-2 structure prediction with high confidence and N- and C-termini predicted with low confidence and mostly unstructured, indicating that N-termini are most likely disordered protein regions that are not involved or impairing CasΦ-2 folding and accessible for protein-protein interaction for Afp packing events (Fig. S27).

### Prediction of NtSP patterns using logistic regression

The absence of packing of *S. entomophila* Afp17 highlights that direct genetic neighborhood is not sufficient for packing inside the Afp particle and that some proteins encoded close to syringe encoding-genes could be pseudo toxins or toxin remnants. To investigate whether we can predict which NtSP sequences lead to cargo packing, we gathered sets of sequences and labeled them as negative or positive based on whether they caused packing (see methods). We implemented several statistical models to predict whether a sequence was positive or negative, including a logistic regression model as well as a model using autocorrelation of the physicochemical properties of the amino acids (see methods). We found the logistic model to have a mean cross-validation accuracy of 93.0 %, better than the autocorrelation model (88.4 %), and a baseline model always predicting the most important class (‘negative’, 80.9 % accuracy), although this accuracy may be artificially high due to homology between sequences in the test/training split. Using a logistic classifier fitted on each residue in the sequence (accuracy = 96.0 %), we predicted the NtSP sequence “MPYSSNSKKNETHSKKNERD” (CC 1, 3 lag 2 = -1.5964) to have the optimal score suggesting that this sequence may merit more investigation.

### Non-syringe related cargo CasΦ-2 and ExoU Toxin-Effectors located within the tail tube

Jiang *et al.* recently showed that, for the closely related PVCs, effectors are located within the tube, probably unstructured (13). Structural analysis of the Afp particle and Afp18 cargo location was performed by cryo-EM and comparing experimental Afp maps at the baseplate for the particles Afp1-18, Afp1-17, Afp1-16, Afp1-18ΔC4 and Afp1-16+Afp18ΔC8-CasΦ2 and Afp1-16+Afp18ΔC8-ExoU as re-engineered cargo examples (Fig. 5, Table S2.). We did not observe any morphological differences among the particle preparations. The C6-symmetrized EM maps however show additional small density inside the tail tube when Afp18 or another cargo is present. The toxin could be present partially folded or unfolded, however, structural information is limited due to the low signal inside the particle, unstructured or partially structured toxin, and the applied C6 symmetry.

**Figure 5:**
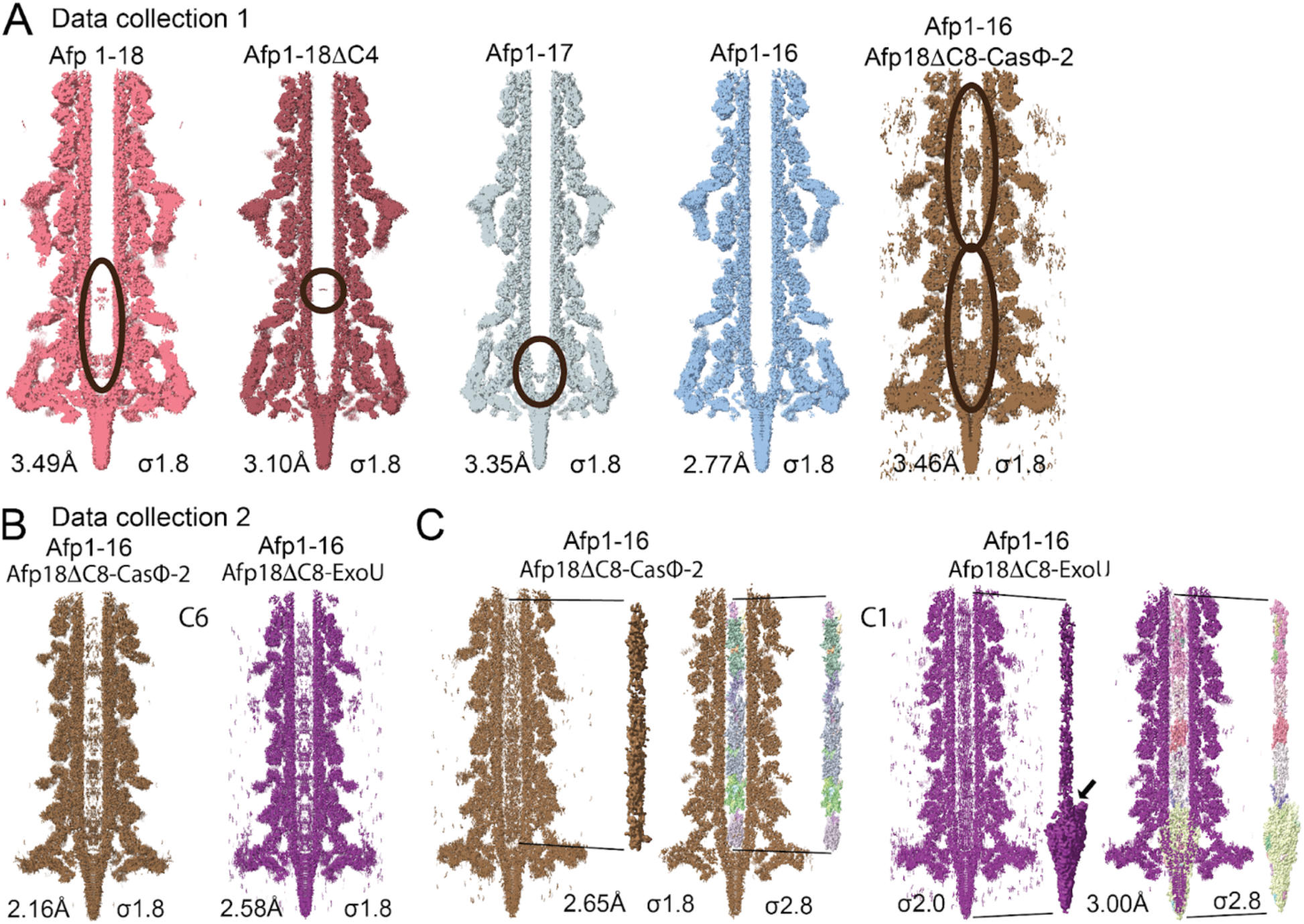
Cryo-EM analysis of Afp maps in native and modified states. **(A)** Overview of different Afp datasets collected on a Titan Krios microscope using comparable settings with a Falcon 3EC direct electron detector (data collection 1): all datasets collected using similar settings and with Falcon 3EC direct electron detector. From left to right: full length Afp (Afp1-18, red) shows small density along the lower third of the tail tube. Afp1-18ΔC4 (dark red), Afp18 truncated by 50%, shows diminished density inside the tube but remaining cargo close to the baseplate. Afp1-17 (grey) with small density close to the baseplate and tip entry (potentially Afp6 helices). Afp1-16 (blue) shows no density in the inner tube. Afp1-16+Afp18ΔC8-CasΦ-2 (brown) shows novel partially structured density appearing all along the tail tube. Afp maps were reconstructed to comparable high resolution (Fig. S32.) **(B)** Overview of different datasets collected on an upgraded Titan Krios microscope with a Selectors X imaging filter and a Falcon 4i direct electron detector (data collection 2): datasets collected on an updated microscope with Falcon 4i direct electron detector. High-resolution cryo-EM maps of two non-eCIS cargos loaded inside the Afp tail tube, Afp18ΔC8 fused to CasΦ-2 (brown) and to ExoU (purple). Density can be observed at various points inside the tail tube. Afp maps were reconstructed to comparable high resolution (Fig. S33.) **(C)** Refinement in C1 and segmentation was carried out with the same settings for both maps (threshold 0.2, dilation radius 5, and soft padding 5). For ExoU the density of the cargo appears connected to the central spike, and for CasΦ-2 the cargo density does not reach the central spike. Tail tube density was low pass filtered 6 Å and is highlighted in brown and purple, segmentation maps (not filtered) are highlighted (multi-color connectivity coloring) within the tail tube. Black arrow highlighting cargo-central spike connectivity for Afp18ΔC8-ExoU cargo.

We investigated the structure of the Afp particle with the effector cargo Afp18ΔC8-CasΦ-2 (104 kDa) because it might be at least partially structured inside the tube and the structure for CasΦ-2 is available, which would allow fitting of structural elements or model building in case part of the protein is structured in the tube (22). Compared to the empty (Afp1-16) Afp particle, Afp1-16+Afp18ΔC8-CasΦ2 shows enhanced pronounced density of Afp18ΔC8-CasΦ-2 within the tail tube at various locations (Fig. 5A). Investigating the distal (thought to be far furthest from the target membrane upon particle binding and contraction) end of Afp, by shifting the box size towards the cap, shows that density inside the tail tube appears all along the inner tube until the cap (Fig. S28&S29., Materials and Methods).

We attempted to improve our reconstructions of Afp particles with the effector cargo by collecting datasets on an upgraded microscope with improved detector (Table S3.). For Afp18ΔC8-CasΦ-2 and with Afp18ΔC8-ExoU particles, we could observe densities inside the tail tube in C6-and C1-symmetrized maps (Fig. 5B&C, Fig. S30.). We attempted to improve the density inside the tail tube, but no atomic model could be built for the cargo Afp18ΔC8-ExoU (Fig. S31.). The tube inner diameter (about 3 nm) is not large enough to hold fully folded cargo in that size (44). It was not possible to determine whether structural features represent Afp18ΔC8 (60 nm length, 30 nm width) or ExoU (about 60 nm diameter).

### Biotechnological Feasibility and Outlook of Afp Modification

The modification of eCIS holds promise for targeted delivery of molecules and drugs. We have shown here that by fusing the Afp18 toxin NtSP, comprising 20 amino acids, and sequence-related NtSPs from other species to other toxins and effectors, these can be successfully loaded into the Afp particle. In order to help build tools for targeted delivery, we attempted to modify the targeting as well as surface modification of Afp particles. We investigated whether replacing Afp tail fibers for fibers of another eCIS, and decoration of the particle sheath can produce intact Afp particles. Similar experiments have been successfully carried out e.g. for T6SS (tail sheath labeling) (45, 46) and for T7/T3-like phages where whole tail fibers or chimeric fibers have been swapped of phages with various host range (47), as well as for PVC eCIS where the putative target recognition domains have been replaced by elements recognizing novel targets with very high efficiency (15).

The Afp particle tail fibers could be successfully replaced with fibers from PVC particles, and as expected, tail fibers appear to be shorter than for wild-type Afp (Fig. 6). Similar complete particle morphology was observed when the Afp3 sheath protein was modified with mCherry on its N-terminus containing a 20 amino acid linker (Fig. 6).

**Figure 6:**
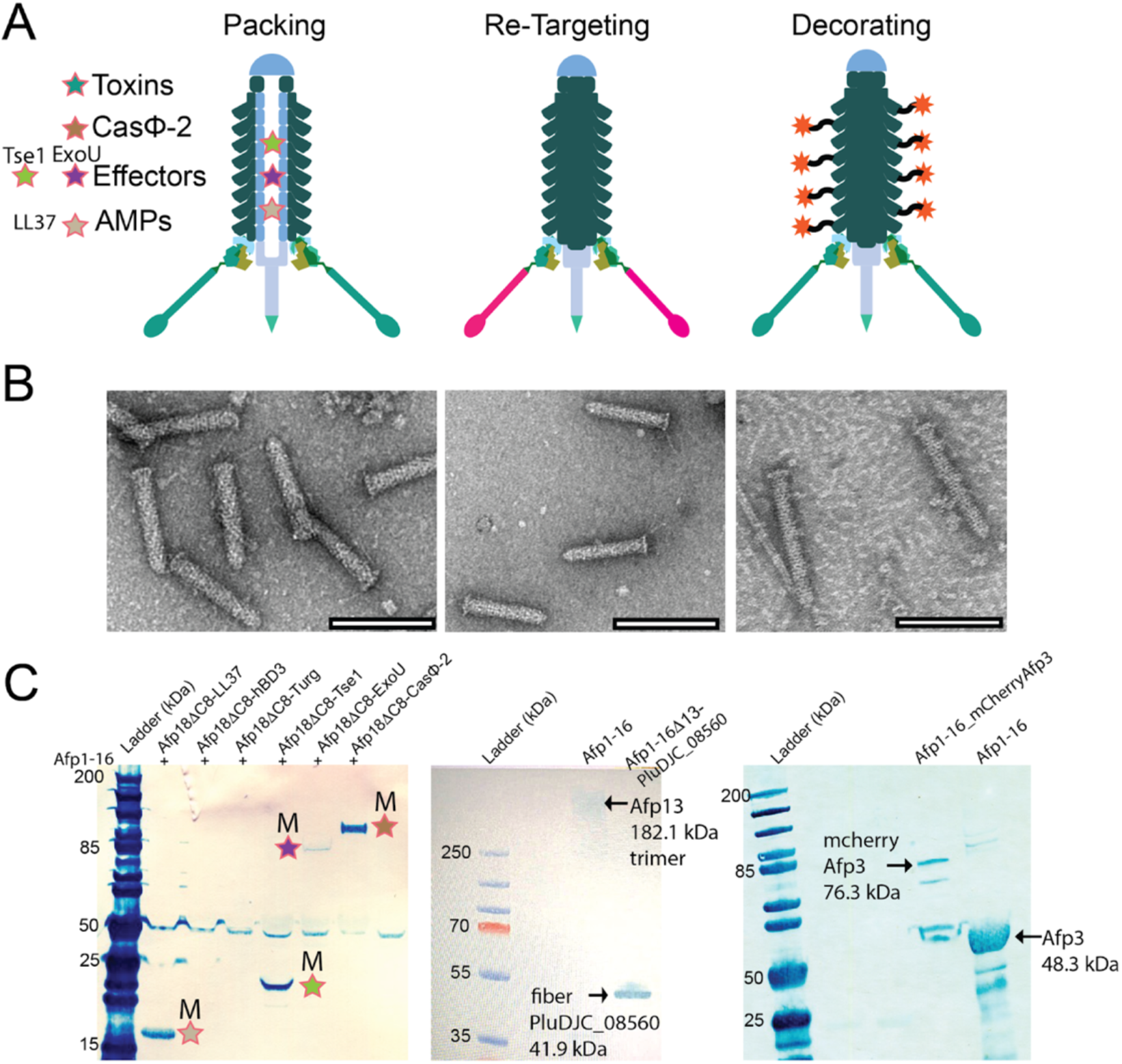
Feasibility of and potential of Afp as biotechnological toolbox. **(A)** Cargo packing, tail fiber exchange and sheath protein decoration. **(B)** Negative stain electron microscope images of modified Afp particles, show particle integrity and complete architecture (scale bars 200 nm) of modifications presented in above panel. **(C)** Immuno-detection of modified cargo, fibers and labeled sheath using specific antibodies, comparison of native and modified (M) Afp particles.

The successful fiber exchange and tail sheath labeling was confirmed by electron microscopy and immuno-detection. We validate that Afp particle architecture is intact for all three modification steps, toxin, fiber and sheath. The results underline the potential and stability of the Afp to serve as a biotechnological scaffold.

## Discussion

Targeted delivery in biotechnology and biomedicine aims to deliver effectors and drugs directly into specific cells while minimizing off-target effects. Bacterial nano-injection systems such as T6SS and T3SS as well as the eCIS PVC, have shown their potential for efficient delivery and modulation of host cell pathways (4–6, 15, 45, 46). However, several challenges remain, including effector loading, studies in animal models, and strain and payload activity *in vivo* (*48, 49*). In this study, we used cryo EM, structure prediction, mutational analysis, cross covariance, and protein design to investigate the packaging of different effectors into the Afp particle.

Our results confirm that the Afp particle is a promising candidate for targeted therapy. Previously, the presence of N-terminal packing domains for PVCs was demonstrated (13, 15). Here, we identified a 20-amino acid domain at the N-terminus of Afp18 as the likely minimal packing sequence in Afp (Fig. 4A). By fusing N-terminal signal peptides (NtSPs) from other species, we successfully loaded unrelated cargoes into the Afp particle (Fig. 4F). In addition, we used cryo EM, immuno-detection, and mass spectrometry to determine the presence and location of effector fusions that were packed inside the tail tube in an unstructured or partially structured conformation (Fig. 5).

Using alignment-independent cross-covariance calculations, we classified putative NtSP of other eCIS and proposed the presence of distinct physicochemical properties that enable efficient peptide packing (Table 2). We postulate that the observations and resilience of packing capability to single point mutations (13) and multiple mutations examined in our study (Fig. 4E) may be attributed to the physicochemical properties associated with efficient peptide packing. These properties likely play a crucial role in maintaining the packing capability of NtSPs. Furthermore, we propose that further optimization of the classifier could enable more reliable prediction of eCIS cargoes and a more efficient packing of cargoes through optimized NtSP sequences. Notably, limitations were observed in packing cationic antimicrobial peptides (AMPs), possibly due to the high abundance of cysteines in their sequences (Table 1). Additionally, we did not detect proteolytic cleavage after cargo packing (Fig. 4D). Interestingly, it was observed that PVC cargos undergo N-terminal cleavage (13), however it is not clear if cleavage is a necessary event for PVC loading. For Afp loading we do observe NtSPs to be present confirmed by immune-detection and mass spectrometry analysis, and toxins and modified cargos e.g. *Yr*Afp17 or Afp18-fusions show low degradation patterns on immune-detection blotting and presence of N-terminal peptides were validated using LC-MS excluding major degradation events at the NtSP region (Fig. 3C&4D, Fig. S21).

In the outlook on regression models for predicting NtSPs, we acknowledge the limitation of having a very limited number of positive and negative tested NtSPs to validate the model. To address this, a high-throughput method for testing could significantly enhance the model’s performance. Similar successful attempts have been made in the prediction of N-terminal effector sequences in T3SS, where comparable or even better scores were achieved (50).

Currently, our algorithm can predict in an alignment-independent way with a confidence level (96%) based on cross-covariance (CC) calculations. However, it is important to note that, despite homology reduction, the reliance on sequence homology may contribute to the accuracy of the predictions. Therefore, expanding the test set with a larger and more diverse collection of positive and negative NtSPs could improve the model’s performance without sequence homology impact. Of course, experimental data on packing of putative toxins would be the best training data, but is naturally very difficult to obtain.

Structural studies of the Afp particle have not yet confirmed the location of the Afp18 toxin, however, its favorable location inside the particle tube was proposed (12) and this is consistent with observations in PVC (13) and algoCIS (51). We used cryo-EM combined with mutagenesis to determine the location of the toxin within the tube. Presence of the toxin in our preparations was supported by immuno-detection and mass spectrometry methods. We conclude Afp18 and other large modified cargoes are packed inside the tail tube in an unstructured or partly structured ‘string of beads’ manner (Fig. 5), as suggested for PVC (13).

Recently, functional studies employing PVCs have demonstrated selective and efficient delivery of protein cargoes into tumor tissues, confirming a promising outcome for engineering the Afp for targeted drug delivery (13, 15). Similarly, we observed Afp1-18 particle lysates cause significant mortality and arrested larvae over time, compared to non-cargo controls (Afp1-16 and Afp1-18). Finally, further modification of Afp was shown to be feasible, by replacement of whole tail fibers and decoration of its sheath (Fig. 6).

The Afp particle shows promise as a highly temperature-stable candidate for the rational design of engineered injection systems. Further research is needed to address limitations, validate predictive models, and explore their potential applications for modification and synthetic biology advances. Overcoming these challenges will be important to unlock the potential of Afp particles as efficient and precise delivery systems for targeted delivery in research, biotechnological and biomedical applications.

## Materials and methods

### Experimental design

The objectives of this study were to develop optimized recombinant production of the *Serratia entomophila* Afp, to obtain insight into the toxin loading mechanism, and to attempt to load exogenous cargoes as well as to optimize this loading process.

### Cloning of *S. entomophila* Afp constructs

The natural plasmid pADAP (GenBank: AF135182) from *Serratia entomophila* Grimont *et al.* 1988 (https://www.ncbi.nlm.nih.gov/Taxonomy/Browser/wwwtax.cgi?id=42906) was prepared with a QIAGEN Plasmid Maxi Kit. The Afp gene cluster *afp1 - afp18* (encoding for Afp1-18) was cloned into a linearized, arabinose-inducible pBAD33 expression vector with chloramphenicol resistance (Cm^R^) (52) by PCR-amplified fragments with overlapping regions in each fragment (Fig. 1). The inserts and the linearized vector, 8.5 - 15kb in size, were amplified using the Platinum™ SuperFi™ PCR Master Mix (Invitrogen) and fragments then gel-purified using Monarch® Genomic DNA Purification Kit (NEB: #T3010S). The pBAD33 vector was DpnI (NEB: #R0176S) digested before gel purification. Fragments were assembled using the In-Fusion® HD Cloning Plus CE kit containing DNA in a 1:1:2 ratio, including the provided Cloning Enhancer. The reaction was incubated for 15 min at 37°C, followed by 15 min at 50°C and 5ul of the reaction mix was transformed into Stellar competent cells. Positive colonies were screened by colony PCR and restriction digest (BamHI or XbaI and KpnI) of the plasmid preparations. The *afp1-17* cluster was cloned as described for a. The constructs *afp1-18ΔC4* and *afp1-16* were cloned the same way as described above, but pBAD33-*afp1-18* and pBAD33-*afp1-17* served as a template, respectively, with two equally sized fragments in the In Fusion® assembly mix (primers Table S4). Engineered *afp* constructs replacing *afp13* with *P. luminescens* fiber *PluDJC_08560* (pBAD33 *afp1-16Δ13_PluDJC_08560*_fibre) and mCherry labeling of *afp3* at its N-terminus (including a flexible linker GSAGSAAGSGEF, pBAD33 *afp1-16*_mCherry-*afp3*) was carried out by PCR amplification of three fragments using pBAD33_*afp1-16* as template and amplification of inserts with overhangs into *afp1-16*. Fragments were gel purified and assembled using In-Fusion® HD Cloning Plus CE kit as described above. The full plasmid sequences were confirmed using Next Generation Sequencing (NGS), showing correct sequences of the whole clusters.

### Cloning of full toxin and toxin truncation constructs for co-expression and purification

The *afp18* toxin gene was amplified from pADAP plasmid (GenBank: AF135182.5, GenBank: CP082788.1) DNA preparations by PCR using the In-Fusion® assembly mix, into an ampicillin resistant pET11a vector, creating *afp18*-3CTS, with a C-terminal Twin Strep Tag (3CTS). *Afp18* in pET11a untagged, was amplified from *afp18*-3CTS with primers including a stop codon before the tag and closing the linear fragment using the KLD enzyme mix from New England Biolabs (NEB). Positive clones were confirmed with colony PCR, restriction digest and correct sequence validated by NGS. The *afp18* homolog, *afp17* (*Yrafp17*) encoded on the *Yersinia ruckeri* ATCC 29473 genome, was cloned and validated in the same way as *afp18* but using genomic *Y. ruckeri* DNA as a template, prepared with a Sigma gDNA GenElute® Bacterial Genomic DNA Kit (Sigma-Aldrich). The hemopexin toxin *PluDJC_08520*, part of a *Photorhabdus luminescens DJC* CIS cluster, was cloned and validated as described above *(P. luminescens DJC kindly provided by Prof. Ralf Heermann, University of Mainz)* (primers Table S4).

To investigate if the N-terminal or C-terminal part of Afp18 is responsible for cargo packing and which part the packing motif is in, we designed a series of N- and C-terminal truncation variants (C1-10 and N7-NX3, see Fig.2C). Truncation borders were chosen based on secondary and tertiary structure prediction programs Quick2D and HHPRED, respectively, provided by MPI Bioinformatics Toolkit (27) https://toolkit.tuebingen.mpg.de (Fig. S6.). The *afp18* truncation constructs were purchased from GenScript Gene Cloning Services, providing *afp18* plasmid as a template. No signal sequence could be predicted for the Afp18 N-terminal residues 1-70 using state-of-the-art programs that employ deep neural networks for signal peptide detection, Signal-P 6.0 server (29) DTU Health Tech, https://services.healthtech.dtu.dk/service.php?SignalP-6.0.

### Cloning of Afp18-toxin constructs for co-expression and purification

To investigate whether toxins of Afp related CIS can be fused and co-purified with Afp18 we designed a set of Afp18-toxin-chimeras. For cloning of homologous effectors, genomic DNA of *Photorhabdus luminescens DJC*, *Photorhabdus asymbiotica* ATCC43949 and *Y. ruckeri* ATCC29473 gDNA was purified based on a phenol-chloroform based protocol (53). The toxin genes *PluDJC_08520* (hemopexin), *PluDJC_12685* (RtxA toxin) from *P. luminescens DJC*, the *afp18* homologue *DJ39_RS03245* (*yrafp17*) from *Y. ruckeri* were cloned after the 3’ end of the DNA sequences encoding C-terminally truncated *afp18ΔC4*(I1437), *afp18ΔC6*(T171). These toxin chimeras were produced by linearizing plasmids encoding C-truncated Afp18 and PCR amplifying selected toxin-encoding DNA regions with 20 nt overhangs into the Afp18 vectors. The Afp18-toxin chimeras were assembled with the In Fusion® assembly mix, clones screened and confirmed as described for cloning of Afp18 (primers Table S4).

### Cloning of Afp18-effector constructs for co-expression and purification

The limit of *afp18* truncations to serve as a scaffold for co-purification and delivery of effector molecules was screened by cloning other secretion system effectors, including non-CIS related cargo, a short antimicrobial peptide (AMPs) and CasΦ-2 from Biggiephage, into the Afp18 C-truncation plasmid. We ordered synthesis and subcloning of Type VI secretion system effectors of *Pseudomonas aeruginosa* PAO1, *tse1* (gene: PA1844, Uniprot: Q9I2Q1), Type III secretion system effectors of *Pseudomonas aeruginosa* UCBPP-PA14, *exoU* (gene: *exoU*, Uniprot: O34208), codon-optimized (for bacterial expression) *casΦ-2* of Biggiephage (22), a short, non-CIS related AMPs, human *ll-37* (Uniprot: P49913, ‘LLGDFFRKSKEKIGKEFKRIVQRIKDFLRNLVPRTES’) *afp18ΔC8* from Genscript. The two non-CIS cargos *ll-37* and *casΦ-2* were cloned into the designed *afp18* N-terminal constructs including the first 50, 30, 20, 11, 5 and 2 amino acids, subcloning was ordered from Genscript.

### Cas**Φ**-2 fusion constructs of Afp18NT20 mutants and NtSPs of other species

Constructs encoding 20 amino acid NtSPs from other species *Yr*Afp17NT20 (*Yr*Afp17NT20:MPYFNKSKKNEIRPEKSKEE), *Se*Tox20 (*Se*Tox20: MPYSSESKLKDTHLKEAESD), *Yp*Tox20 (*Yp*Tox20: MLYSSESKEKKTHSKETERD), CyaANT20 (CyaANT20: MPRYSNSQRTPTQSTKNTRR), *Ep*Tox20 (*Ep*Tox20: MPYFNELNEKETRSKETESG), Afp17NT20 (Afp17NT20: MPTKTPQLQLAIEEFNKAIL), ExoUNT20 (ExoUNT20: MHIQSLGATASSLNQEPVET), and mutant variants of Afp18NT20, Afp18N20KtA (Afp18N20KtA: MPYSSESAEAETHSAETERD, lysines to alanines), Afp18N20KTtA (Afp18N20KTtA: MPYSSESAEAEAHSAEAERD, lysines, threonines to alanines), Afp18N20EtA (Afp18N20EtA: MPYSSASKAKATHSKATARD, glutamic acids to alanines) were synthesized and subcloned into pET11a_*afp18NT20-cas*Φ-*2* (replacing *afp18NT20*) by Genscript.

### Afp protein, toxin and effector-specific rabbit polyclonal antibodies

Afp particle, toxin and NtSP-specific polyclonal antibodies were designed using secondary and tertiary structure prediction programs Quick2D and HHPRED respectively, provided by MPI Bioinformatics Toolkit (27) (https://toolkit.tuebingen.mpg.de) and produced by Genscript (Table S5).

### Expression and purification of Afp particles

The pBAD33-*afp* plasmids were transformed into Electro Competent One Shot™ BL21 Star™(DE3) with pBAD33 expressing *afp1-18*, *afp1-18ΔC4*, *afp1-17* and *afp1-16* and plated on LB-Cm^R^ (25 µg/mL working concentration) plates. Colonies were picked and a starter culture of 10 mL LB-Cm^R^ was grown overnight at 37°C. The next morning, a growth culture was started in 900 mL media-Cm^R^ and induced at OD_600nm_ 0.6 – 0.8 with 0.2% L-arabinose, grown at 18°C for 18-22 hours at slow agitation.

After induction, cells are harvested and resuspended in 25 mL of lysis buffer (25mM Tris pH 7.4, 140mM NaCl, 3mM KCl, 200 μg/mL lysozyme, 50 μg/mL DNase I, 0.5% Triton X-100, 5mM MgCl2 and one tablet cOmplete™ Protease Inhibitor Cocktail from Roche) and incubated for 45 min at 37°C. The lysate is cleared for 45 min, 4°C and 18,000xg centrifugation. After clearing the lysate, the particles are pelleted in two ultracentrifugation (UC) rounds, each 45 min, 4°C, 150,000xg and resuspended first in 5 mL, then in 0.5 mL 1xPBS buffer. After the second round of UC, the particles are loaded on an OptiPrep™ gradient ranging from 40%, 35%, 30%, 25%, 20% and 10% prepared in 1xPBS and run for 20-24h, at 4°C at 150,000x*g*. Fractions are harvested in 0.5mL steps and particle location confirmed using SDS PAGE. Particle samples are pooled and dialyzed for 6 days at 6°C in 1xPBS, after which a last round of UC is performed and the particles resuspended in 0.5mL of 1xPBS. Quality of particles is investigated using SDS PAGE, immuno-detection western blotting and negative staining electron microscopy (EM). Particle quality and toxin levels are a reference for further experiments. Analysis of produced particle preparations using SDS PAGE and Coomassie staining, immune detection blotting, electron microscopy and mass spectrometry analysis was performed in one replicate, or more if specified.

### Co-expression and purification of toxins and Afp particle variants

Chemically competent One Shot™ BL21 Star™(DE3) with pBAD33 expressing *afp1-18*, *afp1-17* and *afp1-16* were prepared using a rubidium chloride based standard protocol. *Afp18*, afp18-toxin chimeras and *afp18*-effector chimeras were transformed into chemically competent *afp1-17* and *afp1-16* respectively and colonies selected for chloramphenicol and ampicillin resistance (Cm^R^, Amp^R^) (Fig. 3). As a control experiment, the *afp18* toxin/effector constructs were expressed without Afp particles, termed mock expression – ‘no particle’ samples – to monitor toxin co-purification or insoluble toxin purification. For high throughput co-expression studies, 200 mL of each plasmid combination was cultured and induced at OD_600nm_ 0.6 - 0.8 with 0.25 mM IPTG 30 min prior to 0.2% L-arabinose, grown at 18°C for 18-22 hours at slow agitation. The co-expression protocol was optimized for balanced IPTG/L-arabinose concentrations leading to a detectable toxin to particle ratio. After induction, cells were harvested and resuspended in 3mL of lysis buffer (25mM Tris pH 7.4, 140mM NaCl, 3mM KCl, 200 μg/mL lysozyme, 50 μg/mL DNase I, 0.5% Triton X-100, 5mM MgCl2 and one tablet cOmplete™Protease Inhibitor Cocktail from Roche) and incubated for 45 min at 37°C. The lysate was cleared for 45 min, 4°C and 18,000xg centrifugation. After clearing, the lysates were precipitated with 8% polyethylene glycol (PEG) 6,000 and 0.5M NaCl and slowly agitated overnight in the cold-room (6-10°C). The next day particles were collected with a centrifugation at 4,000xg for 20 min at 4°C and the pellet resuspended in 1mL ice cold 1 x PBS buffer and agitated for 4h in the cold room. Afterwards, remaining precipitation was pelleted for 45 min at 14,000xg and supernatant saved for analysis on SDS-PAGE. Then, the supernatant was ultracentrifuged 150,000xg for 45 min at 4°C to pellet the particles. Analysis of produced particle preparations using SDS PAGE and Coomassie staining, immune detection blotting, electron microscopy and mass spectrometry analysis was performed in one replicate, or more if specified.

### SDS-PAGE analysis

The particles were diluted in 1xPBS to equal concentrations for comparison on SDS-PAGE and Coomassie staining and for immuno-detection blots. The samples were supplemented with reducing Laemmli SDS sample buffer (250mM Tris-HCl, 8% SDS, 40% Glycerol, 8% β-merceptoethanol, 0.02% Bromophenol blue, pH 6.8), boiled for 5 min at 98°C, centrifuged at 14,000xg for 2 min and loaded on Invitrogen™ NuPAGE™ 4-12%, Bis-Tris gels and gels were run at 200 V for 40 min. The gels were stained with Instant Blue™ Coomassie Stain for 30 min and washed with water for several hours before evaluation.

### Immuno-detection blot analysis

The Afp particles were diluted in 1xPBS to appropriate concentrations for visualization for SDS-PAGE and following Immunoblotting and detection by toxin and Afp particle specific antibodies. The samples were prepared as described above (SDS-PAGE analysis) using Invitrogen™ NuPAGE™ 4-12%, Bis-Tris gels (for particles) or Invitrogen™ NuPAGE™ 3-8%, Tris-Acetate (for high molecular weight toxin analysis). The NuPAGE™ 4-12%, Bis-Tris gels were run as described above and the NuPAGE™ 3-8%, Tris-Acetate gels were run at 150 V for 70 min. Afterwards, gels were removed from the plastic shields and washed in water. NuPAGE™ 3-8%, Tris-Acetate gels were soaked for 10 min in 20% Ethanol to allow increased protein blotting. Proteins from respective gels were transferred on iBlot™ Transfer Stack, PVDF membranes for 7 min for Bis-Tris gels and for 10 min for Tris-Acetate gels using Invitrogen™ iBlot® Dry Blotting System. The membrane was washed in water and exposed to particle and toxin specific antibodies using iBind™ Western Devices. Antibodies and Western reagents were prepared using the Invitrogen™ iBind™ Solution Kit with antibody dilutions ranging from 1:100 and 1:1000. Membranes were exposed to TMB-D Blotting Solution (Kementec) followed by scanning and analysis.

### Mass spectrometry of in-Gel analysis of Afp18 toxin

Proteins were separated using precast 4–20% Tris-Glycine SDS-PAGE gels (1.0 mm thick) (Life Technologies, Carlsbad, CA). Protein gel was stained with Simply Blue SafeStain (Life Technologies, Carlsbad, CA) and protein bands of interest were cut out and subjected to in-gel trypsinization. The samples were reduced, alkylated and digested with Trypsin protease in the presence of ProteaseMAX surfactant (Trypsin enhancer) as described (54). Briefly, after reduction with DTT (Sigma) and alkylation with iodoacetamide (Sigma), gel pieces were dried and subsequently rehydrated in solution containing 12 ng/μL Trypsin Gold (mass spectrometry grade from Promega), 0.01% ProteaseMAX surfactant (Promega) and 50 mM ammonium bicarbonate. After 10 min incubation at room temperature, the rehydrated gel pieces were overlaid with 30 μL of 0.01% ProteaseMAX in 50 mM ammonium bicarbonate and incubated at 37°C for 3 hours with shaking at 800 rpm (Thermomixer, Eppendorf). The digestion reaction was transferred to a fresh tube, mixed with formic acid (1% final concentration of formic acid) and centrifuged at 14,000xg for 10 min to remove particulate material. Supernatant was stored at -20°C until LC-MS/MS analysis.

Tryptic peptides were separated on Hypersil Gold AQ C18 RP column 100 mm x 1 mm 3 µm 175 Å (Thermo Scientific) using UltiMate 3000 LC system (Dionex). Mobile phase A composition was 0.1% formic acid, 2% Acetonitrile, and mobile phase B was 97.9% acetonitrile, 2% H2O, 0.1% formic acid. A multi-step gradient was used at a constant flow of 0.15 mL/min. Mobile phase B was linearly increased from 5% to 11% over 5 min, then from 11% to 25% over 25 min and from 25% to 50% in 25 min. The ions were infused into MicroTOF QII mass spectrometer (Bruker) using an ESI source in positive mode. A precursor m/z range of 75-2200 was used followed by data dependent MS/MS acquisition of top 5 most abundant precursor ions in every full MS scan. Data analysis was performed using DataAnalysis 4.0 (Bruker Daltonics). The detected masses were calibrated using sodium formate cluster ions as an internal calibrant infused during sample loading stage of LC gradient. Peptides were identified using AutoMSn (signal intensity cut off at 250) and deconvoluted using peptides and small molecules preset. Detected peptides were submitted to an automated Mascot search for identification (55). The Mascot search parameters were as follows: 1) Taxonomy: All entries, 2) Database: Swissprot, 3) Enzyme: Trypsin, 2 miss-cleavages allowed, 4) Global Mod: Carbamidomethyl (C), 5) Variable Mod: Deamidated: (N,Q), Oxidation: (M), 6) Mass Tol. MS: 10 ppm, MS/MS: 0.05 Da.

### Mass spectrometry analysis of purified multiprotein assembly samples

Samples after purification using the method described under ‘*Co-expression and purification of toxins and Afp particle variants*’ and partly purified further as described in *‘Expression and turification of Afp particles’* (referred to as ‘pure’ samples) were analyzed for protein species content. 100 µL of room temperature 50 mM ammonium bicarbonate was added to 7.5 µg of purified proteins. Following this, 250 ng of sequencing-grade trypsin was added and samples were incubated overnight at room temperature with mild agitation. Samples were reduced and alkylated (using TCEP and chloroacetamide at 10 mM) for 30 min prior to peptide clean-up via high-pH C18 StageTip procedure. C18 StageTips were prepared in-house, by layering four plugs of C18 material (Sigma-Aldrich, Empore SPE Disks, C18, 47 mm) per StageTip. Activation of StageTips was performed with 100 μL 100% methanol, followed by equilibration using 100 μL 80% acetonitrile (ACN) in 200 mM ammonium hydroxide, and two washes with 100 μL 50 mM ammonium hydroxide. Samples were basified to pH >10 by addition of one tenth volume of 200 mM ammonium hydroxide, after which they were loaded on StageTips. Subsequently, StageTips were washed twice using 100 μL 50 mM ammonium hydroxide, after which peptides were eluted using 80 µL 25% ACN in 50 mM ammonium hydroxide. All fractions were dried to completion using a SpeedVac at 60 °C. Dried peptides were dissolved in 20 μL 0.1% formic acid (FA) and stored at −20 °C until analysis using mass spectrometry (MS).

Around 1 µg of digested proteins were analyzed (∼250 ng of peptide) per injection for each sample, as two technical replicates. In this paragraph, “Exp. 1” relates to Fig. S3, “Exp. 2” relates to Fig. S18, and “Exp. 3” relates to Fig. S19. All samples were analyzed on an EASY-nLC 1200 system (Thermo) coupled to an Orbitrap Exploris 480 mass spectrometer (Thermo). Samples were analyzed on 20 cm long analytical columns, with an internal diameter of 75 μm, and packed in-house using ReproSil-Pur 120 C18-AQ 1.9 µm beads (Dr. Maisch). The analytical column was heated to 40 °C, and elution of peptides from the column was achieved by application of gradients with stationary phase Buffer A (0.1% FA) and increasing amounts of mobile phase Buffer B (80% ACN in 0.1% FA). The primary analytical gradients ranged from 5 %B to 32 %B over 30 min for Exp. 1, 5 %B to 34 %B over 40 min for Exp. 2, and 5 %B to 38 %B over 40 min for Exp. 3. All gradients were followed by a further increase of 10 %B over 5 min to elute any remaining peptides, and followed by a washing block of 15 min. Ionization was achieved using a NanoSpray Flex NG ion source (Thermo), with spray voltage set at 2 kV, ion transfer tube temperature to 275 °C, and RF funnel level to 40%. Full scan range was set to 300-1,300 m/z, MS1 resolution to 120,000, MS1 AGC target to “200” (2,000,000 charges), and MS1 maximum injection time to “Auto”. Precursors with charges 2-6 were selected for fragmentation using an isolation width of 1.3 m/z and fragmented using higher-energy collision disassociation (HCD) with normalized collision energy of 25. Monoisotopic Precursor Selection (MIPS) was enabled in “Peptide” mode. Precursors were prevented from being repeatedly sequenced by setting dynamic exclusion duration to 50 s (Exp. 1) or 60 s (Exp. 2 and Exp. 3), with an exclusion mass tolerance of 15 ppm and exclusion of isotopes. For the 2^nd^ technical replicate of Exp. 1, dynamic exclusion was set to trigger only after attempting to sequence the same precursor twice within 10 s. MS/MS resolution was set to 45,000, MS/MS AGC target to “200” (200,000 charges), MS/MS intensity threshold to 230,000, MS/MS maximum injection time to “Auto”, and number of dependent scans (TopN) to 9.

All RAW files were analyzed using MaxQuant software (version 1.5.3.30). RAW files corresponding to Exp. 1, 2, and 3 (as described above, relating to Fig. S3, S18, and S19) were analyzed separately. Default MaxQuant settings were used, with exceptions outlined below. For generation of the theoretical spectral library, all expected full-length protein sequences were entered into a FASTA database. Digestion was performed using “Trypsin/P” in semi-specific mode (which allows non-specific cleavage on either end of the peptide), with a minimum peptide length of 6 (for Exp. 1) or 7 (for Exp. 2 and 3) and a maximum peptide length of 55. Protein N-terminal acetylation (default), oxidation of methionine (default), deamidation of asparagine and glutamine, and peptide N-terminal glutamine to pyroglutamate, were included as potential variable modifications, with a maximum allowance of 3 variable modifications per peptide. Modified peptides were stringently filtered by setting a minimum score of 100 and a minimum delta score of 40. First search mass tolerance was set to 10 ppm, and maximum charge state of considered precursors to 6. Label-free quantification (LFQ) was enabled, “Fast LFQ” was disabled, and “Skip normalization” enabled. iBAQ was enabled. Second peptide search was disabled. Matching between runs was enabled with a match time window of 1 min and an alignment time window of 20 min. For Exp. 2 and 3, matching was only allowed between same-sample technical replicates. Data was filtered by posterior error probability to achieve a false discovery rate of <1% (default), at the peptide-spectrum match, protein assignment, and site-decoy levels. The mass spectrometry proteomics data have been deposited to the ProteomeXchange Consortium via the PRIDE (56–58) partner repository with the dataset identifier PXD043850.

### Electron microscopy (EM)

Afp particle quality and integrity were investigated using negative-stain electron microscopy. For negative staining, aliquots of 4 μl of Afp samples were added onto copper grids with a continuous carbon support film. The grids were washed with distilled water, and stained with 2% uranyl acetate. The grids were dried at room temperature and analyzed on a Morgagni 268 transmission electron microscope at 100 kV. Cryo-grids of purified Afp particles were prepared with an FEI Vitrobot Mark IV at 4°C and 95% humidity in the environmental chamber. 4 μl of sample (0.5 mg/mL concentration) was applied onto freshly glow-discharged S373-7-UAUF UltrAuFoil QF - R2/2 (200 mesh). After 10s they were blotted using a blot force of -1. Cryo-grid screening was performed on a Tecnai G2 20 TWIN 200 kV transmission electron microscope.

For high resolution data collection, movies were collected using the automated acquisition program EPU (FEI, Thermo Fisher Scientific) on a Titan Krios G2 microscope operated at 300 kV paired with a Falcon 3EC direct electron detector (FEI, Thermo Fisher Scientific). Images were recorded in linear mode, at 75,000x magnification with a calibrated pixel size of 1.1 Å and under focus range of -0.5 to -2.0 μm (0.3 μm steps) with a dose rate of 67.24 e^-^/Å^2^/s, 35 e^-^/Å^2^ and total exposure time of 0.59 s, 23 fractions 6,500 exposures (Afp1-16); 69.87 e^-^/Å^2^/s, 39 e^-^/Å^2^, 0.59 s exposure time, 23 fractions 16,504 exposures (Afp1-17); 67.26 e^-^/Å^2^/s, 38 e^-^/Å^2^, 0.60 s exposure time, 23 fractions 5,445 exposures (Afp1-18ΔC4); 69.87 e^-^/Å^2^/s, 39 e^-^/Å^2^, 0.57 s exposure time, 23 fractions, 9,741 exposures (Afp1-18) (Table S2).

Datasets Afp1-16+Afp18ΔC8-CasΦ-2 and Afp1-16+Afp18ΔC8-ExoU were collected on the same Titan Krios G2 microscope operated at 300 kV but in the meantime upgrade with a Selectris X image filter and Falcon 4i direct electron detector (Thermo Fisher Scientific). Images were recorded by EPU software in counting mode at 165,000x magnification with a calibrated pixel size of 0.725 Å and under focus range of -0.5 to -2.0 μm (0.3 μm steps) and total exposure time of 2.11 s, leading to a final dose of 37 e^-^/Å^2^ and 45 e^-^/Å^2^, respectively. A total of 7,014 exposures (Afp1-16+Afp18ΔC8-CasΦ-2); and 11,638 exposures (Afp1-16+Afp18ΔC8-ExoU) were collected (Table S3).

### Cryo-EM data processing and analysis

All cryo-EM data processing was performed in cryoSPARC (59, 60) (Fig. S28). For all datasets, movies were motion-corrected using full-frame or patch motion correction. The CTF was estimated using patch CTF or CTFFIND4 (61). Micrographs were inspected for CTF quality, motion correction and ice contamination. For all reconstructions shown in this manuscript, details about particle numbers and reconstruction parameters can be found in Table S2&S3.

### Baseplate reconstructions

Particles were initially picked using blob picker (particle diameter 400-600 Å), extracted with a box size of 800 pixels and downsampled to 600 pixels and classified using 2D classification. Good classes containing base plates were then used in template-based picking. After 2D classification, *ab initio* models were constructed and used for homogeneous 3D refinement with C6 symmetry imposed. For datasets Afp1-18 and Afp1-18ΔC4, the *ab initio* model of Afp1-17 was used for homogenous 3D refinement, substantially improving map quality. Datasets Afp1-16+Afp18ΔC8-CasΦ-2 and Afp1-16+Afp18ΔC8-ExoU were processed as above but particles were extracted in 1100 box size and binned to 800. The Afp1-17 *ab initio* reconstruction was used for the homogenous refinement in C6 and in C1 symmetry.

### Cap reconstructions

For cap reconstructions, the box center of all baseplate reconstructions was shifted by 525 pixels towards the cap using the volume align tool in cryoSPARC. Particles were re-extracted with a box size of 800 pixels, reconstructed using homogeneous reconstruction only without particle alignment but with C6 symmetry imposed and a final homogeneous refinement with C6 symmetry imposed (Fig. S28. & Table S2&3.)

### Bioinformatic Investigation of NtSP like domains on eCIS cargos

#### Identification of conserved N-terminal packing motifs in homologous CIS particles

We used the Afp18NT20 sequence MPYSSESKEKETHSKETERD as input sequence for a protein homology (above 60%) or pattern search using BLASTP® to search for peptide homologs (raw output file: 79TBY202013-Alignment.txt, where 79TBY202013 is Blastp JobID). Manual investigation of each accession code (e.g., WP_049612744.1) for gene and genome location (NCBI nucleotide/genome database, www.ncbi.nlm.nih.gov) and validation of being next to homologous CIS particle. The homologous N-terminal packing motifs, accession codes, presence of CIS particles are summarized in Table S1. NtSP from Table S6 were aligned using Multiple Sequence Alignment server ClustalOmega and highlighting for amino acid abundance as logo (WebLogo, https://weblogo.berkeley.edu/, or Seq2Logo https://services.healthtech.dtu.dk/service.php?Seq2Logo-2.0, Fig. S34).

#### Alignment-independent cross covariance (CC) calculations and polar amino acid content

Since alignment-based methods cannot account for gaps in motifs and disrupt the alignment to highlight similar motif properties (a consensus sequence), we highlight parameters and qualification as packing motif using alignment-independent approaches, via cross covariance (CC) which is a transformation of peptide sequence into uniform vectors of principal amino acid properties described in z scales (42). Two vectors, characterized in that said packing motif result in auto cross covariance (CC) deviating lower than zero, where the first vector comprises the amino acid hydrophilicities (z1 scale) of each amino acid in said packing domain and second vector (z3 scale) comprising the electronic properties, represent the molecule’s charge and polarity, having a lag of 2 for said first vector or said second vector.

The covariance is in a preferred embodiment the cross-covariance, calculated in accordance with the following equation I:

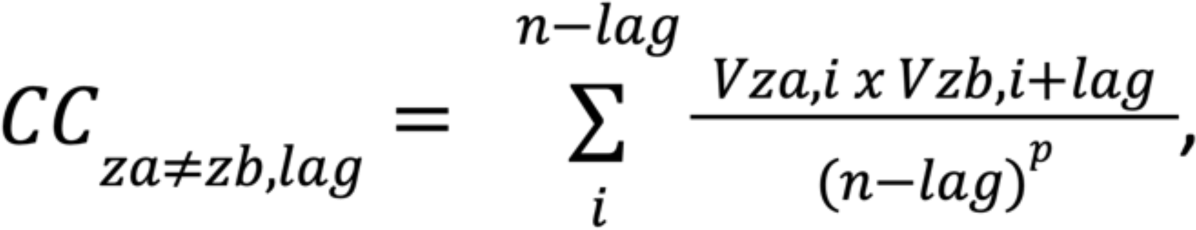

wherein CC is the cross covariances between the first z1 comprising amino acid hydrophobicity of each amino acid in said packing motif and z3 comprising the electronic properties of each amino acid in said packing motif, *i* is the position of each amino acid and is a number between 1 and 20, *n* = 20 is the number of amino acids comprised by the vector, *l* = 2 is the lag, p is the normalization degree and V is the descriptor value.

The VaxiJen 2.0 server offers CC calculations for bacterial peptides with ACC output for all z-scales and combinations and was used to investigate NtSP’s properties (http://www.ddg-pharmfac.net/vaxijen/VaxiJen/VaxiJen.html) (62).

We used ProtParam (https://web.expasy.org/protparam/) to calculate individual amino acid content and manually investigated total polar amino acid content along NtSP’s.

Potential NtSP’s with more than 60% of polar amino acids, preferably balanced and interspersed distribution of positive polar (lysine (K), histidine (H), arginine (R)) and negative polar (glutamic acid (E), (D)) supported by other polar amino acids asparagine (N), serine (S), threonine (T), glutamine (Q) in optimized sequence distribution to achieve high negative ACC1,3 (lag=2) values. We investigated the N-terminal packing domains for packing motifs of *P. luminescens* and *Photorhabdus asymbiotica* CIS particles (Table. S6). Particle related cargos are highlighted with their negative CC1,3 (lag = 2) values and more than 60% polar amino acid content.

### Regression models for predicting cargo packing from NtSPs

We constructed a set of ‘positive’ and ‘negative’ NtSPs, i.e. those we believed would lead to packing and those we did not believe would cause packing. In the positive dataset we included the 6 natural NtSPs for which we have experimental evidence of packing, 3 NtSPs obtained from mutational analysis, and 6 prospective NtSPs found by homology search of public databases (see below). The negative dataset consisted of 3 sequences experimentally seen not to cause packing, 4 negatives found by mutational analysis, 48 antitoxins obtained from a search for known *E. coli* K12 toxin antitoxin systems in UniProtKB and 21 toxins from types III, VI and VII secretion systems which we assume to be not related or too distantly related to the eCIS to be able to cause packing. The positive dataset was homology reduced to 90% identity. We used three-fold cross validation to test various models implemented in scikit-learn v1.3.0 to predict whether a sequence was positive or negative: A naive model always predicting the most common class (accuracy = 80.9 %), a simple logistic model that used the count of each amino acid as input (accuracy = 93.0 %), a model that uses auto cross correlation of physico-chemical properties from Hellberg et al. (42) (accuracy = 88.4 %), and a logistic model that used the full sequence represented as a one-hot matrix as input (accuracy = 96.0 %). From the latter model, the weights could be extracted for each position to obtain the sequence that maximized the predicted probability of packing. However, we note that a homology reduction threshold of 90% means that sequences in our dataset were still homologous, such that all models suffered from data leakage between the training and validation splits. For that reason, we have low confidence in the assessment of our models’ accuracy. Indeed, the models presumably just learned to recognize any sequence that looks like Afp18NT20, variants of which comprise much of the positive dataset, hence why the “optimal” sequence shares 15/20 aa with this sequence. Setting a stricter homology reduction threshold of 50% reduces the size of our positive dataset to just 4 sequences, too low for validating a statistical model. All toxins found using homology search are provided in the supplementary as fasta files (possible_toxins.faa).

### Bioinformatic investigation of eCIS like regions in public databases

To search for putative eCIS systems, we curated a set of marker proteins which we expected to be present in all eCIS systems of the families afp1, afp5, afp11 and afp15. To do this, we gathered a list of bacterial strains with eCIS subtype Ia based on Chen *et al.* (3), (Supplementary Table 3), and extracted all proteins of the above families from these strains, as well as the *Serratia* and *Yersinia* species from which we had experimental evidence, from dbeCIS (http://www.mgc.ac.cn/dbeCIS/). We then searched for these protein against NCBI’s databases env_nt and nt_prok (version 2022-06-14) using tblastn v2.13.0 (min identity 25%, coverage 50%) and extracted all DNA loci that were within 50 kbp of at least one member of all marker protein families. We searched for homologs to our experimentally validated toxins in these loci using tblasn with the same parameters, after which we extracted the 20 N-terminal amino acids, and homology reduced these with a 90% identity threshold. The search yielded 8 new potential NtSPs (Table S7).

### Afp particle efficacy on *Galleria mellonella* larvae

*E. coli* BL21 star cells carrying the pBAD33 constructs encoding *afp1-16*, *afp1-17*, *afp1-18* and empty pBAD33 (used as a control) were grown and induced as described in section ‘Expression and Purification of native Afp particles’. Thereafter, the cells were collected via centrifugation, 5,000 rpm for 20 min and washed 3 times with PBS buffer. Protein extraction was performed via sonication followed by centrifugation, 5,000 rpm for 20 min, and filtration using a 0.2 µm filter to clear cells debris. To ensure that the syringe and toxin components were produced and present in the protein lysate in about the same amounts, SDS-PAGE and immuno-detection against toxin and Afp particle sheath was performed. For testing Afp particle toxicity *in vivo*, 10 *Galleria mellonella* larvae were injected with 30 µl of filtered protein lysates of the respective Afp constructs into the posterior proleg. 30 µl of PBS buffer were injected as a control group to ensure that the solution used for the nanoparticle extraction was harmless to the larvae. The injected larvae were kept at 30 °C and observed for 13 days. Experiments were stopped when controls were fully evolved to moths. Phenotypic interpretation was carried out as follows: ‘Dead larvae’ are not responsive upon pinch stress and present a dark color. ‘Arrested larvae’ are slightly responsive upon pinch stress, however, do not progress in their development to moths in comparison with the control groups. ‘Alive larvae’ are responsive upon pinch stress and develop into moths in the 13 days of experiment. Three independent experiments were performed. Dead and arrested larvae were plotted as percentages using Prism9 as individual values of the three independent experiments, as well as mean and standard deviation. A two-way ANOVA with Dunnett multiple testing (Afp particles compared to pBAD33 control) was performed in Prism9 (P values: 0.0332 (*), 0.0021 (**), 0.0002(***)). The family-wise alpha threshold and confidence level was 0.05 and 95%, respectively.

For testing the effect of Afp particles loaded with toxin-chimeras, the workflow was the same as described above with the following differences. Additional samples: pBAD33 constructs with *afp1-16* co-expressed with *afp18ΔC8* - *P. aeruginosa* Type III SS Effector *exoU* (PAExoU), *afp18ΔC8-LL37* human antimicrobial peptide and *afp18NT20-LL37* human antimicrobial peptide, respectively. Injected larvae were observed for 7 days and only the number of dead larvae was counted since toxin-chimeras led to the death of all 10 larvae after 7 days. Dead larvae were plotted as percentages using Prism9 as individual values of the three independent experiments, as well as mean and standard deviation. A two-way ANOVA with Dunnett multiple testing (Afp particles compared to pBAD33 control) was performed in Prism9 (P values: 0.0332 (*), 0.0021 (**), 0.0002(***)). The family-wise alpha threshold and confidence level was 0.05 and 95%, respectively.

## Statistical Analysis

Sequences were obtained from the National Center for Biotechnology Information, Uniprot or dbeCIS (http://www.mgc.ac.cn/dbeCIS/). Multiple sequence alignments and analysis were performed using Clustal Omega and MView. Sequence logo was created using Weblogo (https://weblogo.berkeley.edu/). Two-way ANOVA with Dunnett multiple testing was performed to confirm statistical significance at 95% confidence of samples compared (P values: 0.0332 (*), 0.0021 (**), 0.0002(***)).

## Acknowledgments

We thank Victor Klein De Sousa for support with AlphaFold2 and the Danish Cryo-EM Facility at the Core Facility for Integrated Microscopy (CFIM) at the University of Copenhagen and Michael James Johnson and Nicholas Heelund Sofos for training and support. Thanks to Eleonora Nigro for her input on scientific illustrations (Scientific Illustrator & Live Drawing Events www.eleonoranigro.com.)

## Contributions

N.M.I.T. and E.M.S.R. conceived the project and designed experiments. E.M.S.R. and R.N.E. set up the purification protocols. E.M.S.R. created the mutants and together with R.N.E. performed biochemistry experiments and most of the analysis of biochemistry and biochemical data. E.M.S.R. prepared cryo-EM samples, EM grids and together with T.P. collected the cryo-EM images. E.M.S.R., M.P.R. and C.K. performed the rest of the cryo-EM processing and cryo-EM map analysis. L.M.A. and R.N.E. supported cryo-EM data analysis and data submission. Larval *in vivo* efficacy assays were carried out by A.R. and K.G. in consultation with R.H. E.M.S.R. and M.P.R. prepared samples for mass spectrometry analysis and I.A.H. carried out mass spectrometry validation in consultation with M.L.N. I.P. performed and analyzed in-gel digest and mass spectrometry analysis. J.N.N. searched NCBI for potential new eCIS and did the regression modeling in consultation with S.R. The global results were discussed and evaluated with all authors. E.M.S.R. and N.M.I.T. coordinated and supervised the project. E.M.S.R. wrote the manuscript with input from all the authors.

## Data and materials availability

All data needed to evaluate the conclusions in the paper are present in the paper or the supplementary materials. Cryo-EM maps were deposited in the EMDB. The final 3D maps of baseplates from data collection 1: Afp1-18 (EMD-18524), Afp1-18ΔC4 (EMD-18526), Afp1-17 (EMD-18551), Afp1-16 (EMD-18552), Afp1-16+Afp18ΔC8-CasΦ-2 (EMD-18525), and from data collection 2: Afp1-16+Afp18ΔC8-CasΦ-2 (EMD-18527), Afp1-16+Afp18ΔC8-ExoU (EMD-18553) in C6 and Afp1-16+Afp18ΔC8-CasΦ-2 (EMD-18528), Afp1-16+Afp18ΔC8-ExoU (EMD-18580) in C1 symmetry, and from Afp-caps in C6 symmetry from data collection 1: Afp1-18 (EMD-18530), Afp1-Afp18ΔC4 (EMD-18531), Afp1-17 (EMD-18575), Afp1-16 (EMD-18576), Afp1-16+Afp18ΔC8-CasΦ-2 (EMD-18532), and from data collection 2: Afp1-16+Afp18ΔC8-CasΦ-2 (EMD-18577), Afp1-16+Afp18ΔC8-ExoU (EMD-18579) in C6. More cryo-EM data collection details are shown in Table S2 and Table S3.

Results from the in-gel digest and LC–MS data analysis are attached in Supplementary Material and as pdf, 2019-08-06_EMR_band1-TD_Mascot-NCBIprot.pdf. The mass spectrometry proteomics data have been deposited to the ProteomeXchange Consortium via the PRIDE partner repository with the dataset identifier PXD043850. LC–MS data used to generate tables and figures has been provided as a .xlsx Source Data file as MS_Merged_Supplement.xlsx. Raw data from experiments of Afp particle efficacy on *G. mellonella* larvae has been provided as .xlsx file Larvae_assay_Afp_constructs_20230718.xlsx. AlphaFold models are represented in Supplementary Material.

## Code availability

The code used for the regression models and searching the NCBI public databases is publicly available at https://github.com/jakobnissen/ecis_search.

## Supplementary Materials

Supplementary material is attached as *.docx file: ‘advances_supplementary_materials_template_Rebrova-et-al-2023.docx’.

