## Supplementary Material for "N-terminal toxin signal peptides efficiently load therapeutics into a natural nano-injection system"

Steiner-Rebrova *et al.*

**This PDF file includes:**

Figs. S1 to S34
Tables S1 to S7
References (1 to 4)

**Supplementary Material**

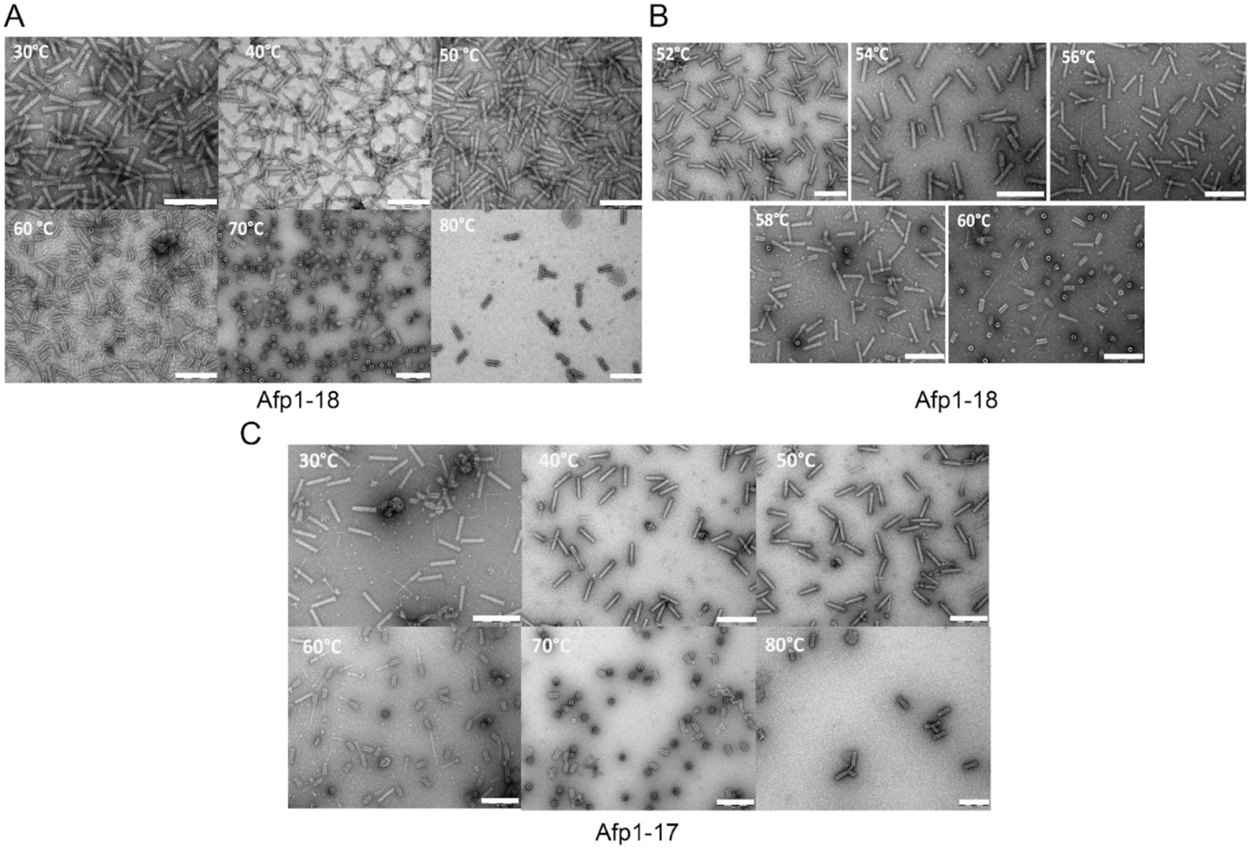


**Fig. S1.**

**Afp temperature stability for particles including and excluding Afp18**

Particles with toxins (Afp1-18) and without Afp18 toxin (Afp1-17) were diluted in PBS to about 0.2 mg/mL concentrations and a 20 ml sample heated for 10 min at respective temperature. 4 µl of the sample was used for negative staining electron microscopy and particle morphology investigated (scale bar 200 nm). (A) Afp particles contracted at temperatures higher than 60℃. (B) Smaller temperature steps investigated between 50-60℃ showing that Afp particles start contracting between 58-60℃. (C) Afp1-17 particles show similar stability behavior with contracted particles at 60℃.

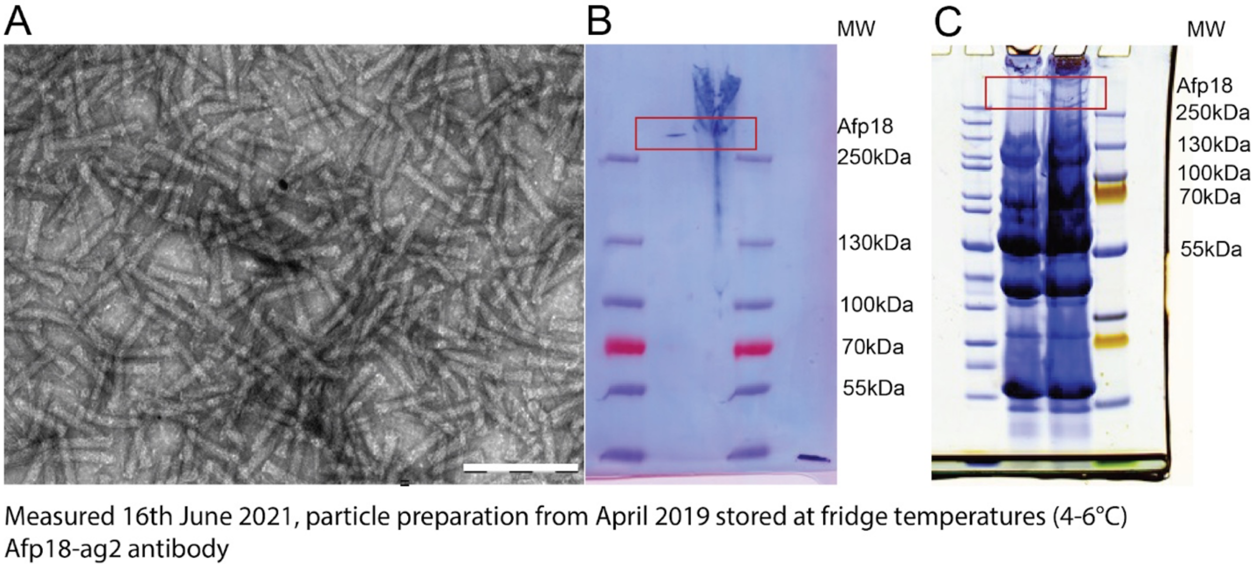


**Fig. S2.**

**Long-term stability of Afp preparations**

Syringes are intact after more than 2 years storage at fridge temperatures (4-6°C) and toxin was detected. (A) Negative staining electron microscopy (EM) image of 2 year old particles (scale bar 200 nm). (B) Immune-detection blotting of Afp18 toxin using Afp18-Ag2 antibody and (C) Coomassie gel of particles and high molecular weight (MW) toxin band highlighted (red box). Particle preparation from April 2019 and measured June 2021.

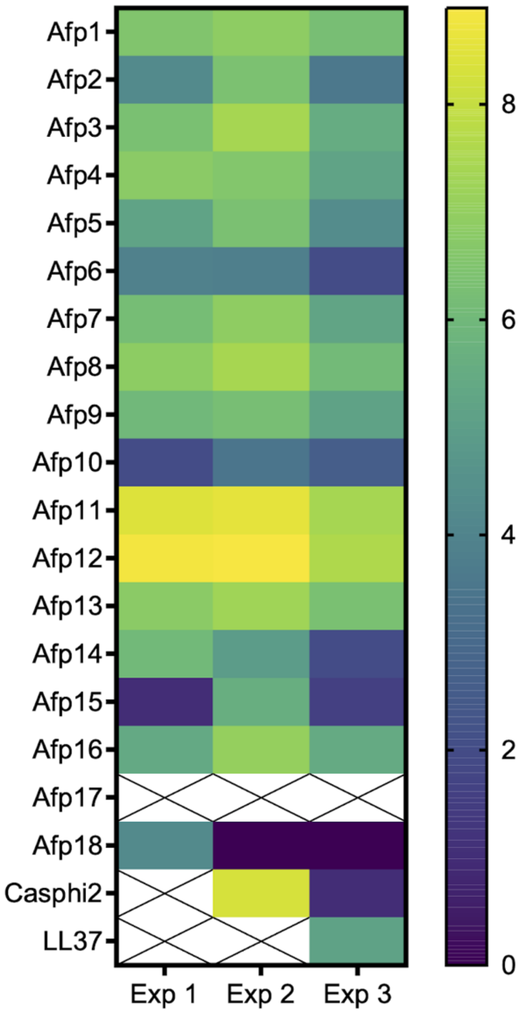


**Afp18 toxin sequence coverage**

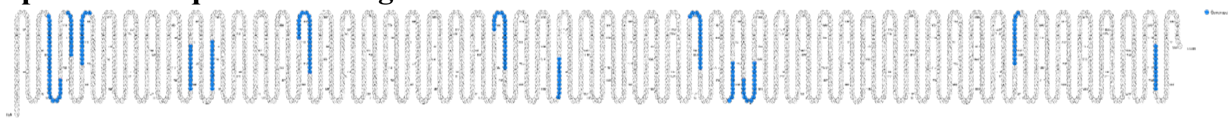

**Fig. S3.**

**Mass spectrometry analysis of Afp particle preparations**

Heatmap visualization of mass spectrometry analysis of particle assemblies Afp1-18 (Exp 1), Afp1-16+Afp18ΔC8-CasΦ2(Exp 2, Casphi2), and Afp1-16+Afp18ΔC8-LL37 (Exp 3). Afp17 could not be detected in any of the three particle preparations; crossed out cells indicate no detection. The heatmap visualizes $log_2$-transformed spectral (MS/MS) counts of peptides mapping to each protein, *n*=2 summed technical replicates. Afp18 sequence coverage is visualized below (blue, sequence covered by peptides).

### Spectrum Analysis Report

**Date: 08/09/2019 Time: 11:09**

**FileName: C:\Data\mass spect data\Aug-2019\20190806_225_EMR_band1-TD_GA5_01_11061.d\20190806_225_EMR_band1-TD_compounds.xml**

#### Sequence data:

hypothetical protein [Serratia entomophila] WP_010895820.1

| | | | |
|---|---|---|---|
| Intensity Coverage: | 0.7 % (26863 cnts) | Sequence Coverage MS: | 53.8% |
| Sequence Coverage MS/MS: | 53.8% | pI (isoelectric point): | 6.7 |

```
        10         20         30         40         50         60         70         80         90        100        110        120
MPYSSESKEK ETHSKETERD NADPVFQRVS QGGVGVSPPD EGGDLSAGSN NDHLFAFIRE THLEEAQEFR LKGYNSQVPE VDEGLLTQLR DDLAAAKSRQ YVQASVGGPL GSRWLLEQIG
       130        140        150        160        170        180        190        200        210        220        230        240
KIERLTDKFS DEVIKRWVKH ELPRKNVGEA AGTELGSRRR AQSLARRLES TMQTLIRVVH TLHAMDDYST PEKRHDMLVA ELTGAFSSSD KNTVIQALGR MTAETSGHAA WASLMQARTA
       250        260        270        280        290        300        310        320        330        340        350        360
AWEAGKSATV DAMIATSQHE AEKANKKARE LQGDSQAFFS HVAAYLQSLS SDLAKVSINM GQSTSSSTIR NEAEKASELQ GDSQTFVSHV AAYLQSLPSD LAKVPINMWQ NMDFVKTKLQ
       370        380        390        400        410        420        430        440        450        460        470        480
AQTAVAGMKV HAHRLVRIAQ HGHSTKTPAK DPEQVVTDSI IRSILWLWQQ PAIKIQCAST ALLSKVGELK KIEGILASYA VTDELGNEQE RRNPPGEEDL DAQLREWVSA SLKQENPENQ
       490        500        510        520        530        540        550        560        570        580        590        600
RAVKVVTLER LLGGDIASAR NLVERLGKAE EGIQNLLRRQ QVAVLKMSVE RLPAAVSSLK AVDKLLPDLA GDLTASIAAL SNALQAAEYP ARDFSEAKIQ AGYAQLWATK VKESLSAASA
       610        620        630        640        650        660        670        680        690        700        710        720
RLTERPLDEH SRGARLAKHW ANLAKERSVG NYPPPDAQQV LASLKEQGLL AGTLSTGDPA GYLFATRLAG ELENAGNGEL RLPMSPEQYV ALEKGLVEYI VKWGQRRISQ GVTRIVIELS
       730        740        750        760        770        780        790        800        810        820        830        840
FEQALDTVSF NVSKLFRLPY KVLKASIKIP YNVNKVNNYT MPGHDKPYKA IYGLLGKKLK QLGFNLLTAP VPGVMKLVAG TGITAGAGVH NLRVENREKT FSAVYQHVVE GKQSEKIKMD
       850        860        870        880        890        900        910        920        930        940        950        960
SVKSMIFDSV LDTATTAAFK GGRRAWQSGR NKNDIEINEV TLSQAQQRQV AANNVDTAWG NAAANERKVE PVSHSSPLQQ QTDVRPFSDK SDQDAAQPRV RKRAVADMAP PPPHQWHNNI
       970        980        990       1000       1010       1020       1030       1040       1050       1060       1070       1080
PANTTLQSEH FDFDRDIRYQ GFSDEKKKQT YLHGIQLVLL QIQNDGRFAQ KIRNNAYLAS IGVSNLVPVD IEGYKLKNTF LLPDSPSAKS GVIVRLDSEN PYYYVGKGSD LLEDIKWAMP
      1090       1100       1110       1120       1130       1140       1150       1160       1170       1180       1190       1200
HDANEQKYEY MPDSDGNAHF YRLRNGFDIF NDIRMGNSSF EYFFNYNNPE PISITRLSSY LANTIEGDYK NGSSPPKNKL LISRAITGAH IPDPGVTATA AGYHIEFTWD NLTSAEYLRS
      1210       1220       1230       1240       1250       1260       1270       1280       1290       1300       1310       1320
FSRPFSTLSG EMQLVSSSIN GETIQETELH VHQAEYIGSW VDATVGAITS FTPEGWVLNT TLSAADIAAD LTEDKDPDPM AVAGLVVGCI PGGRIAAKVG KFTRIGGKVV KYGLMLGNKA
      1330       1340       1350       1360       1370       1380       1390       1400       1410       1420       1430       1440
VDLAIVGKSV KIAVDTGDPL AIYKALLISG IGTTNSYDMA KNMSSKLKIS KKLEDSVQLD KPLQNLRGGS KRPEEGESTS QKRSQPDESP ENSQEAVYSG PSGRARVTFD QTTEHHIMLN
      1450       1460       1470       1480       1490       1500       1510       1520       1530       1540       1550       1560
RFSRLVLARS FAEKRIQSFE HSSSKFKSSD EFKVDKSIPL DAQNASLNDM ASKTDNLINQ LEQFTGDKKE SISANLKNIQ LMIGDIMENR NNRNDNLNFH TYVLVGKNTT YDEISNYYGF
      1570       1580       1590       1600       1610       1620       1630       1640       1650       1660       1670       1680
AEFKFSINNG NLTLNNLIGH PYVVINKYPE FRDYLIRNGV ITQPELEPYN IKNVARYLGS KAVSSEIGYY ESIPATKVKI LSFTGANPIT QKLGNTLQSI SDLFSKKRFS DEPNDAITQD
      1690       1700       1710       1720       1730       1740       1750       1760       1770       1780       1790       1800
ELFTRQTDLY TYDEMINHLK TIVRTTFTPV KPIEEISSEK TDAAIKGIMT NGDAEIISEN IIESKNSLDH DTSYFSHEVI SAVDRAYEKV NSVKSLFERA ESQPEIKSQL KSLINEATGL
      1810       1820       1830       1840       1850       1860       1870       1880       1890       1900       1910       1920
NDEVIIDIDK VDVSSISDMA YNRFKDNINT LHDYLSKQKS GGYTSFVLFK YRQFNPSKPD AFAMPYDSKK RIFLAMLPEN QRHGGLVDTV THEASHNSSY TLDHTYIGNA RSNTGSFPSS
      1930       1940       1950       1960       1970       1980       1990       2000       2010       2020       2030       2040
FEPPTRDSRY FYENERVGKL SAKYALDLPE NANITQNQLA QADILLRNSR LTKADTLLNA AEYNAYMIDV LSRVKINGSH IEFEENSSRS KRSVSGGNHK LDDIIILASM NATLPRREGG
      2050       2060       2070       2080       2090       2100       2110       2120       2130       2140       2150       2160
EFESSTFKKT LTEFINKGTT SSQVDPVYAS YLPAFNTIVD SEGITLQQVK DLKSALPATG NDLRAQARYN YAVLQYLSGS AGIKLPIDTD VIRNSINDDL NIASCVDNCK KAASQRLELL
      2170       2180       2190       2200       2210       2220       2230       2240       2250       2260       2270       2280
KAVPEYLSGP EAIHAQIDTN LMINRIKNTL KHENYTYDLK VNERDYLLLQ LEQKYIASSK PPKVGAQKWS ADINKEINYL CDEIEEKKRE VVCWENKKNE SSRRLDLWST ATKYLSDPKG
      2290       2300       2310       2320       2330       2340       2350       2360       2370
IKPPIDTDVI LNNINDVLNH AGYVYNYKVR EKDNLEKKLS RTPLTSQIEV NRLQTEIAKR YRLIDYWGKK ADETLLMFIL WRNVFG
```

#### Display Parameter:

| | | | | | |
|---|---|---|---|---|---|
| Sequence Name: | hypothetical protein [Serratia entomophila] WP_010895820.1 | | | | |
| MH+ (mono): | 1.008 | MH+ (avg): | 1.008 | Threshold (a.i.): | 0.000 |
| Tolerance (Da): | 0.050 | Number of Peaks: | 615 | | |

#### Peaklist:

| Peak | Mass | Intensity | Peak | Mass | Intensity | Peak | Mass | Intensity |
|---|---|---|---|---|---|---|---|---|
| 1 | 238.212 | 5169.000 | 2 | 239.198 | 1849.000 | 3 | 256.222 | 8208.000 |
| 4 | 256.263 | 1106.000 | 5 | 261.115 | 1244.000 | 6 | 268.243 | 3777.000 |
| 7 | 279.150 | 701.000 | 8 | 284.318 | 1724.000 | 9 | 285.271 | 6123.000 |
| 10 | 288.285 | 3059.000 | 11 | 288.286 | 3059.000 | 12 | 293.202 | 982.000 |
| 13 | 295.184 | 1154.000 | 14 | 295.187 | 802.000 | 15 | 297.237 | 4182.000 |
| 16 | 299.160 | 1255.000 | 17 | 299.170 | 692.000 | 18 | 301.137 | 12125.000 |
| 19 | 301.204 | 3122.000 | 20 | 304.281 | 1579.000 | 21 | 304.291 | 1411.000 |
| 22 | 308.205 | 9786.000 | 23 | 309.200 | 1364.000 | 24 | 311.253 | 1206.000 |
| 25 | 312.357 | 1631.000 | 26 | 313.237 | 789.000 | 27 | 317.109 | 1025.000 |
| 28 | 320.193 | 1946.000 | 29 | 321.234 | 858.000 | 30 | 321.317 | 676.000 |


Copyright © 1999-2008 Bruker Daltonik GmbH

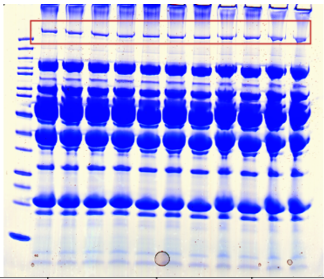

| Date | Sample ID | Type of MS analysis | Mascot ID | Mascot score | Seq coverage (User), % | Coverage range | Contaminants | Comments |
|---|---|---|---|---|---|---|---|---|
| 1-Aug-19 | EMR_03 | Intact protein | WP_010895820.1 Hypothetical protein [Serratia entomophila] | 5655 | 53,80% | 29-2339 aa | No other proteins were indentified in the digest of the SDS-PAGE band | Mascot search against NCBIprot database indentified Hypothetical protein [Serratia entomophila] WP_010895820.1 with high confidence (Mascot score 5655). Mapping of the detected tryptic peptides onto the expected protein sequence gives 53.8 % with covergade range 29-2339 aa. |

**Fig. S4.**

**In-gel digest and mass spectrometry (LC-MS) analysis of Afp18 toxin band**

Provided gel for in-gel digest (Afp18 toxin highlighted with red box) and mass spectrometry analysis of Afp1-18 preparation used for cryo-EM. Afp18 could be identified with 53.8% sequence coverage. Protein coverage from amino acid 29-2339 with high confidence (Mascot score 5,655).

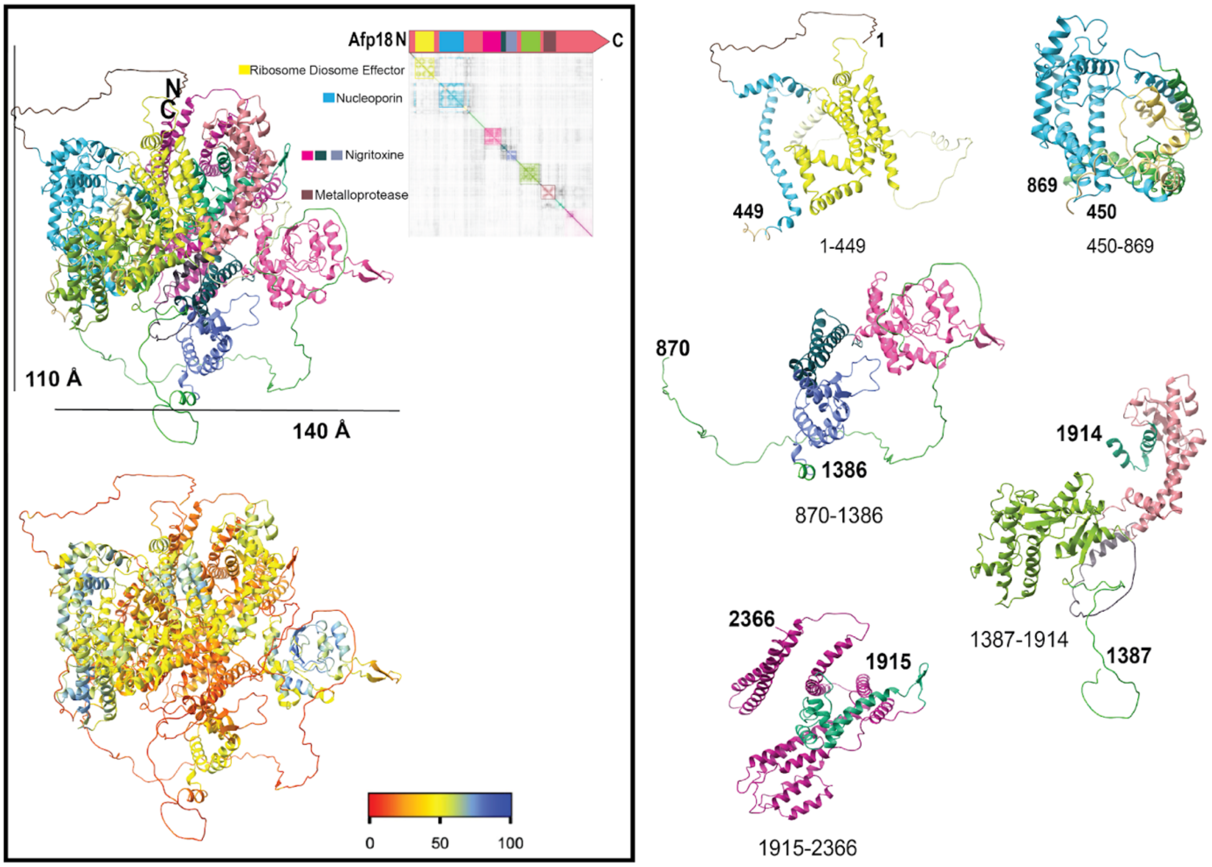


**Fig. S5.**

**Afp18 structure prediction using AlphaFold2**

Afp18 appears to have a pearl-chain like structure with large number of disordered regions, interspersed by rigid domain cores highlighted with a community clustering approach (https://github.com/tristanic/pae_to_domains, top left) that extracts protein domains from a predicted aligned error (PAE) matrix and and per-residue confidence metric (pLDDT) coloring (bottom left) in ChimeraX. Clustered domains are represented and structural homology using HHPRED can be found to Ribosome Disome Effector (yellow), Nucleoporin (blue), an insecticidal toxin, Nigritoxin (pink, dark green dark blue), and a Metalloprotease (brown). For better visibility of structural features, respective domains are shown on the right.

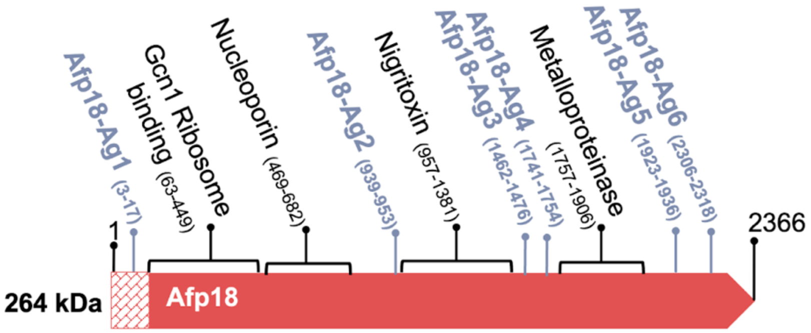


| HHPRED | Effect | PDB *[Ref]* | Probability (%) | aa area |
|---|---|---|---|---|
| *Gcn1 Ribosome Disome effector Saccharomyces cerevisiae Ribosome, Disome, GCN1, Translation, GAAC, ISR, Rbg2, Gir2; HET: 5CT; 3.9A {Saccharomyces cerevisiae S288C}* | Ribosome, Diosme Binding | 7NRC_A *[1]* | 56.22 | 63 - 449 |
| *Nucleoporin NSP1; nuclear pore complex, inner ring, protomer, Saccharomyces cerevisiae, TRANSPORT PROTEIN; 3.71A {Saccharomyces cerevisiae}* | Nucleopore for transport into eukaryotic cells | 7WOO_L *[2]* | 73.47 | 469 - 682 |
| *Nigritoxine; toxin, new fold, arthropod; 2.1A {Vibrio nigripulchritudo}* | *Bacterial toxin against crustaceans and insects* | 5M41_A *[3]* | 100 | 957 - 1381 |
| *NEUTRAL PROTEASE II; METALLOPROTEINASE, ZINC, NEUTRAL PROTEASE II, HYDROLASE; HET: EDO; 1.0A {ASPERGILLUS ORYZAE} SCOP: d.92.1.12* | Bcterial toxin | 1EB6_A *[4]* | 97.21 | 1757 - 1906 |

**Fig. S6.**

**Afp18 HHpred structural similarity search and antibody (Ag) design**

Designed polyclonal antibodies for Afp18 tracking Afp18-Ag1-6 are used to track toxin presence (blue). The HHpred search result of the first op hits are summarized in the table below.

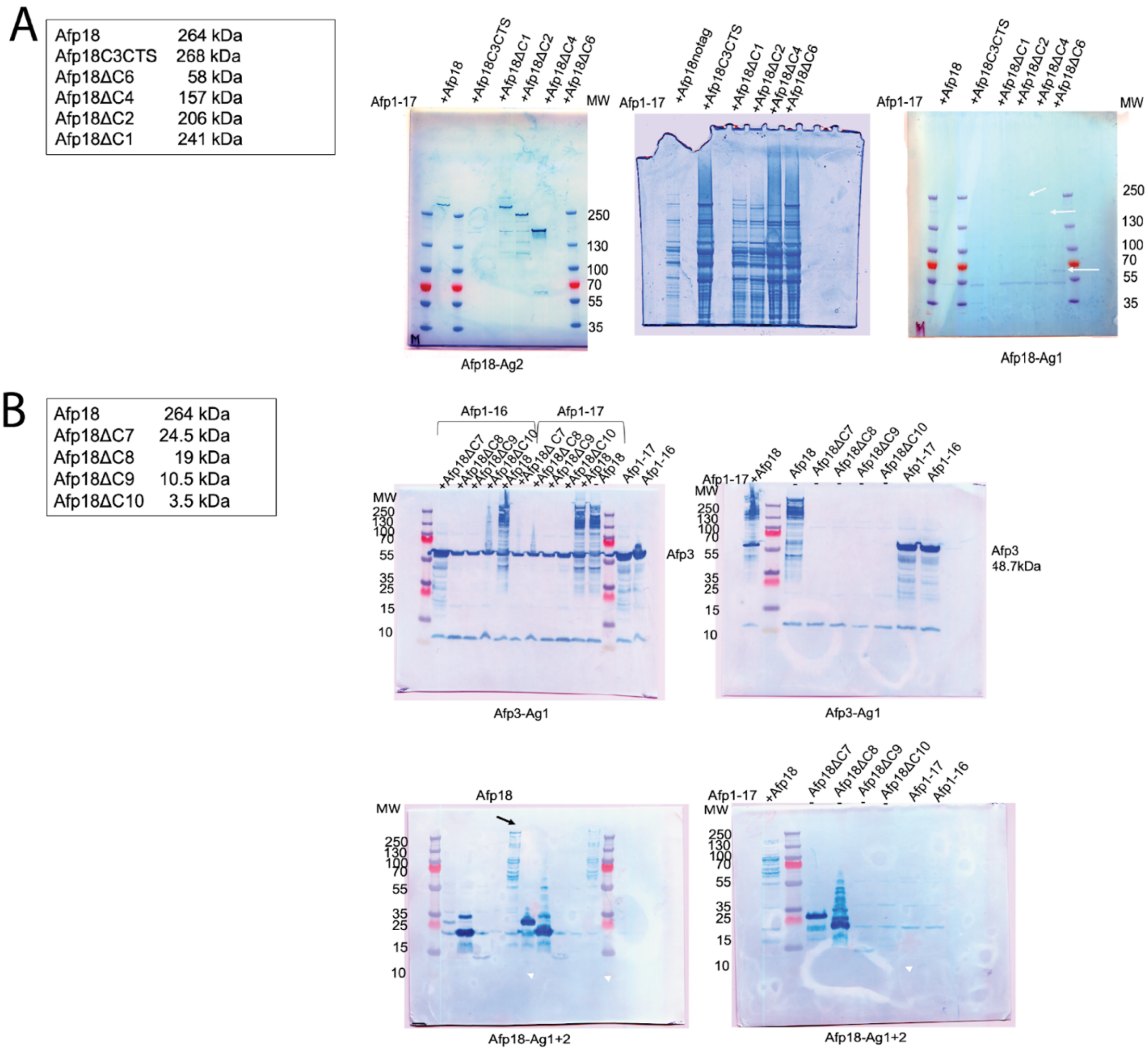


**Fig. S7.**

**Afp18 toxin C-terminal truncations and Afp particle co-production and mock expression**
(A) Afp18ΔC constructs co-produced with Afp syringe Afp1-17 and toxin detection after PEG precipitation and UC. (B) Afp18ΔC constructs co-produced with Afp syringe Afp1-17 and Afp1-16 and toxin detection in lysate samples. Detection limit for C-terminally truncated Afp18 toxin is Afp18ΔC9 (10.5 kDa).

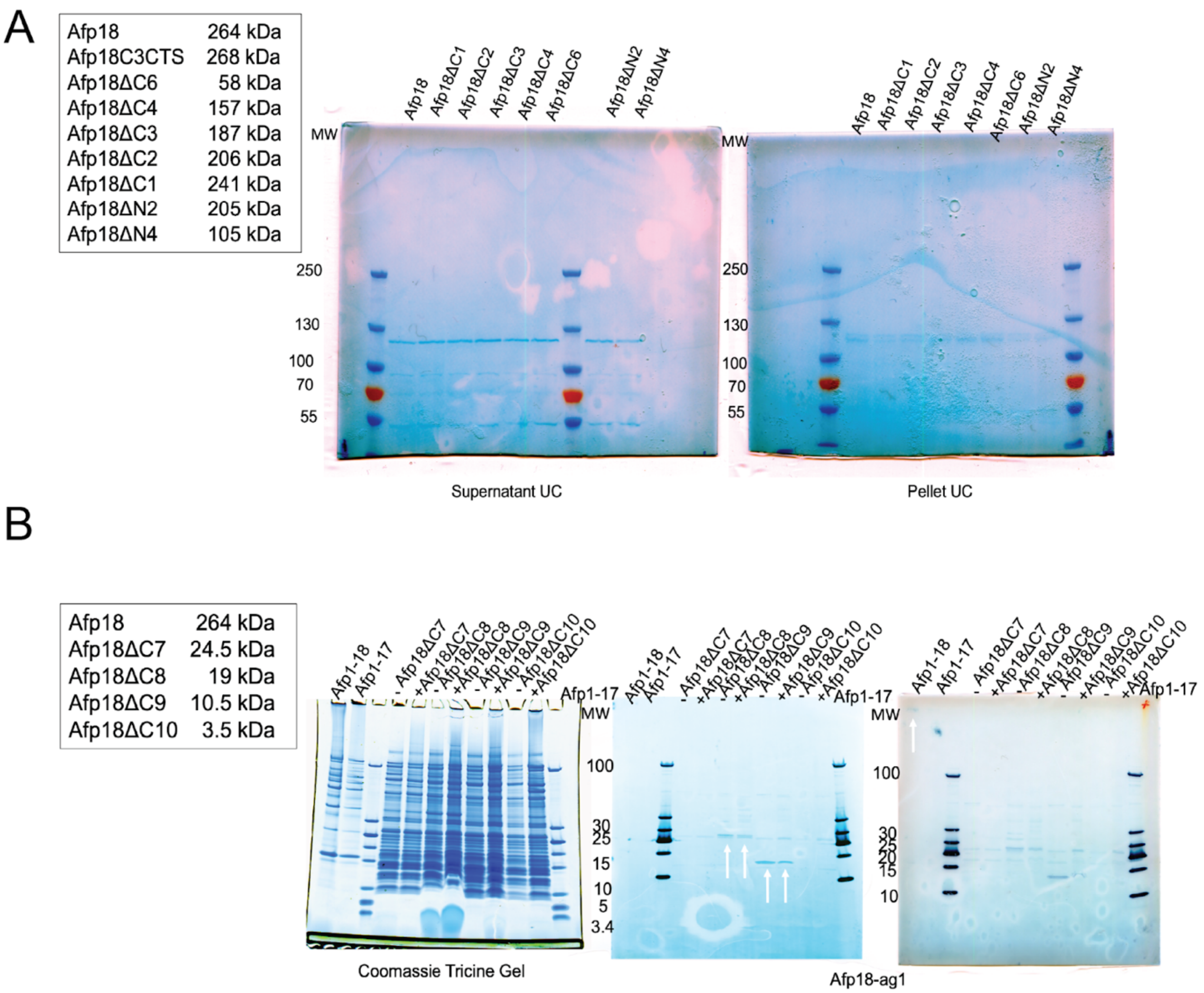


**Fig. S8.**

**Afp18 toxin C-terminal truncations and Afp particle co-production and mock expression**

(A) Production of Afp18ΔN and -ΔC constructs and investigation of mock expression. No toxin co-purification level could be detected. (B) Immune detection and Coomassie staining using tricine gels for low molecular weight proteins. White arrows indicating that small Afp18ΔC8 and C9 variants show positive mock expression and Afp1-18 as a control (Afp18-Ag1 used for detection).

| | |
|---|---|
| Afp18 | 264 kDa |
| Afp18ΔN1 | 239 kDa |
| Afp18ΔNX1 | 244 kDa |
| Afp18ΔNX2 | 253 kDa |
| Afp18ΔNX3 | 260 kDa |

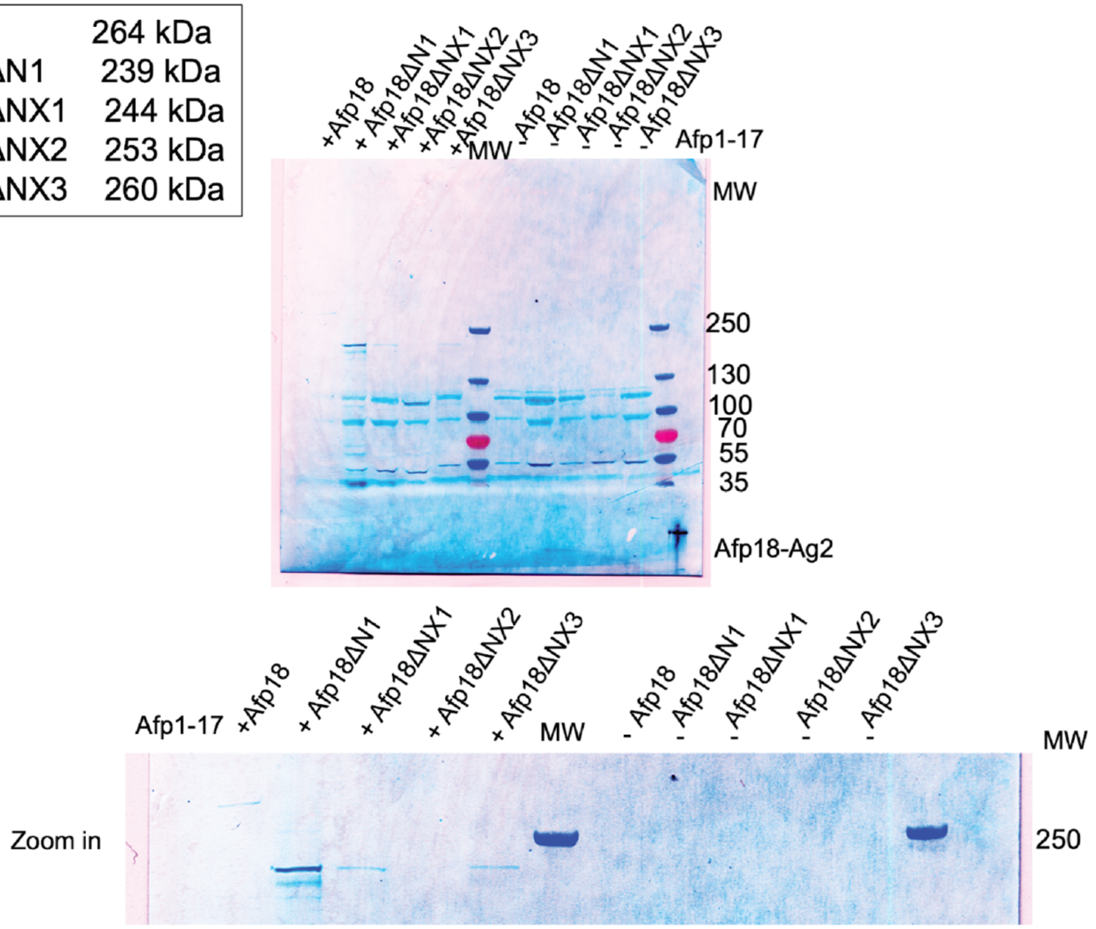


**Fig. S9.**

**Afp18 toxin N-terminal truncations and Afp particle co-production and mock expression**

Production of Afp18ΔN particles and mock expression and purification. Samples analyzed on immune detection blotting are after PEG precipitation and an ultracentrifugation step (UC). Respective Afp18ΔN toxin bands are of lower molecular weight than expected.

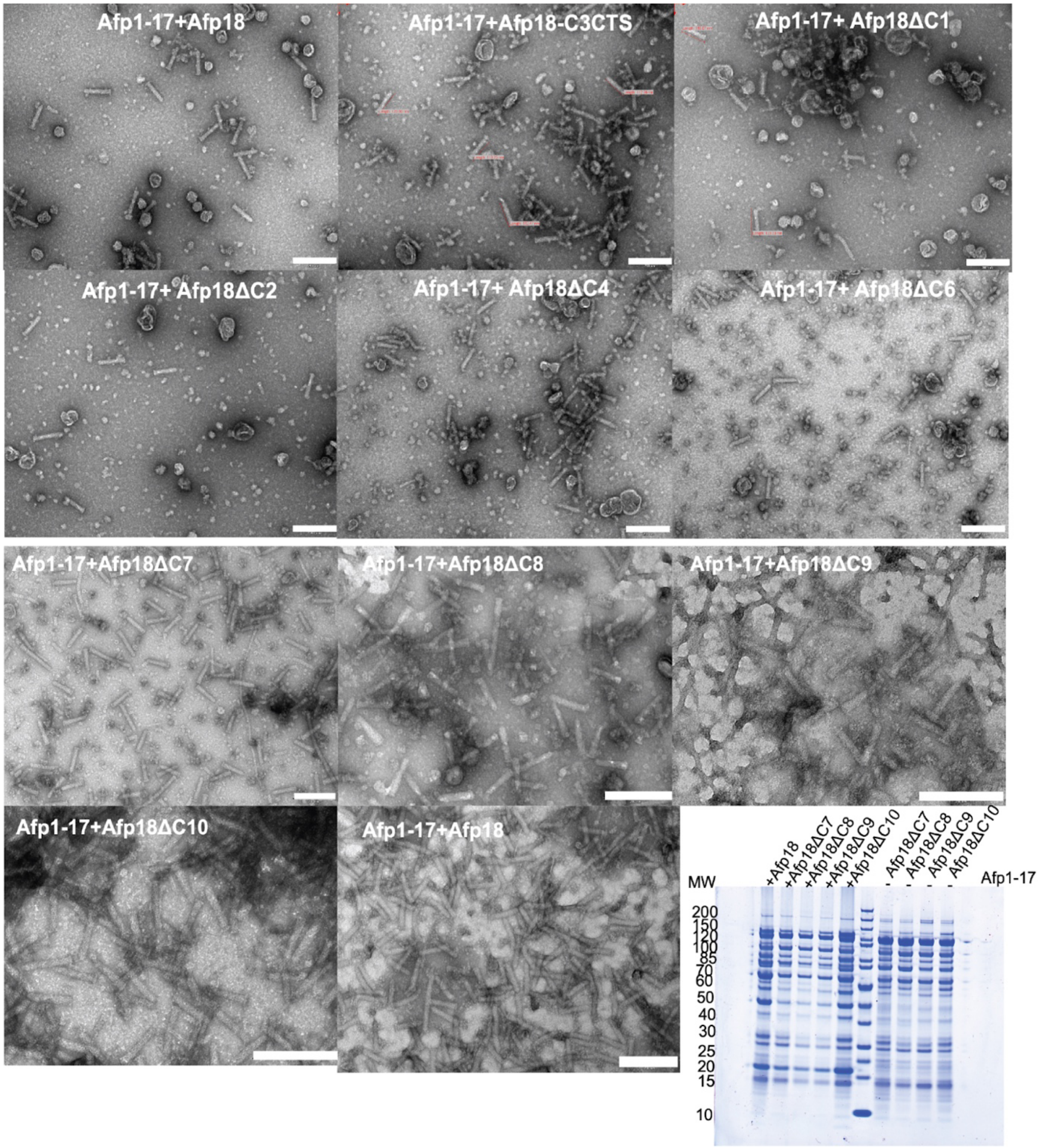


**Fig. S10.**

**Afp18ΔC toxin truncation and particle validation using negative staining EM**

All particles show intact morphology (scale bar 200 nm). Proteins are analyzed as well using Coomassie staining (bottom right corner).

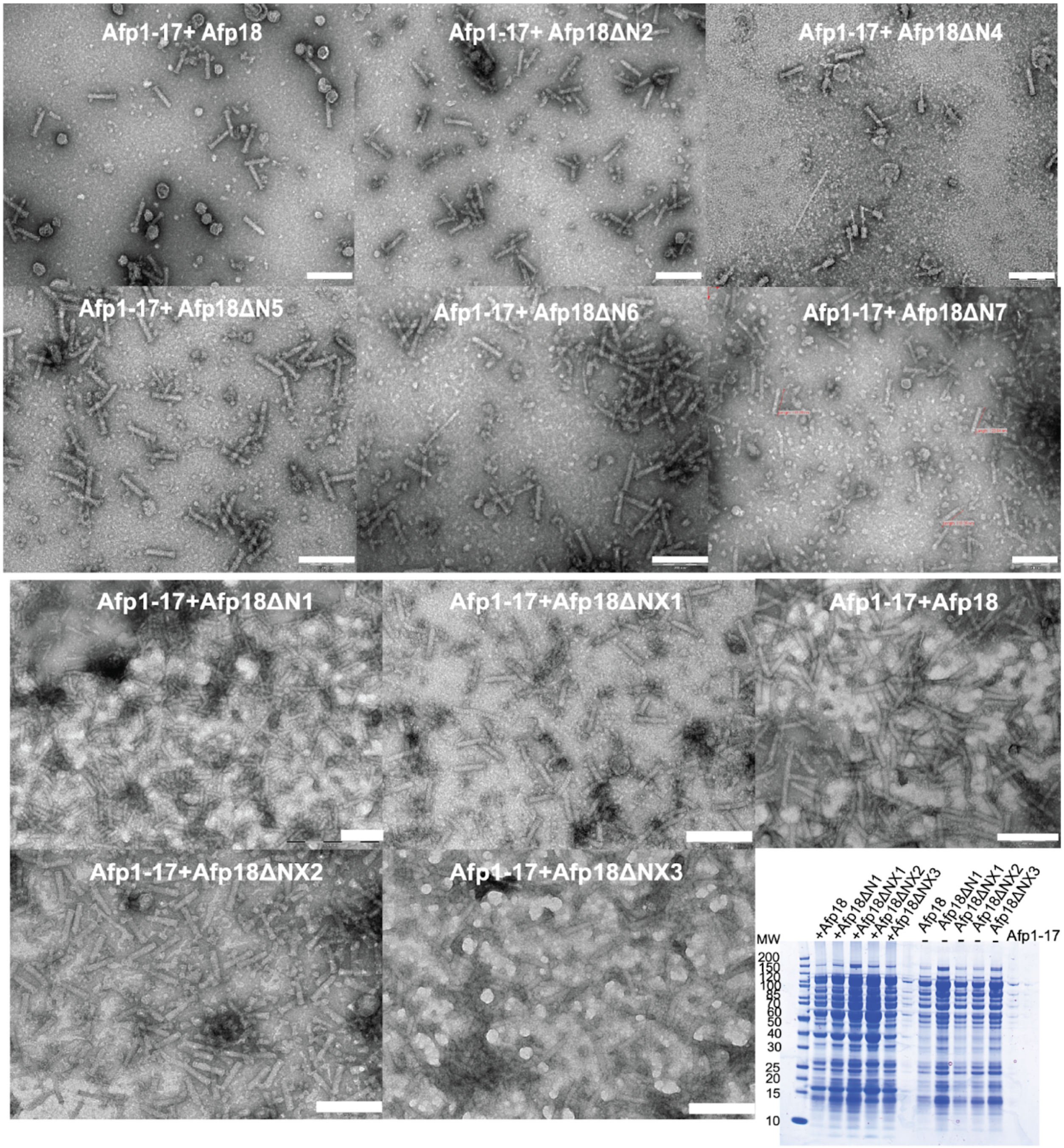


**Fig. S11.**

**Afp18ΔN toxin truncation and particle validation using negative staining EM**

All particles show intact morphology (scale bar 200 nm). Proteins are analyzed as well using Coomassie staining (bottom right corner).

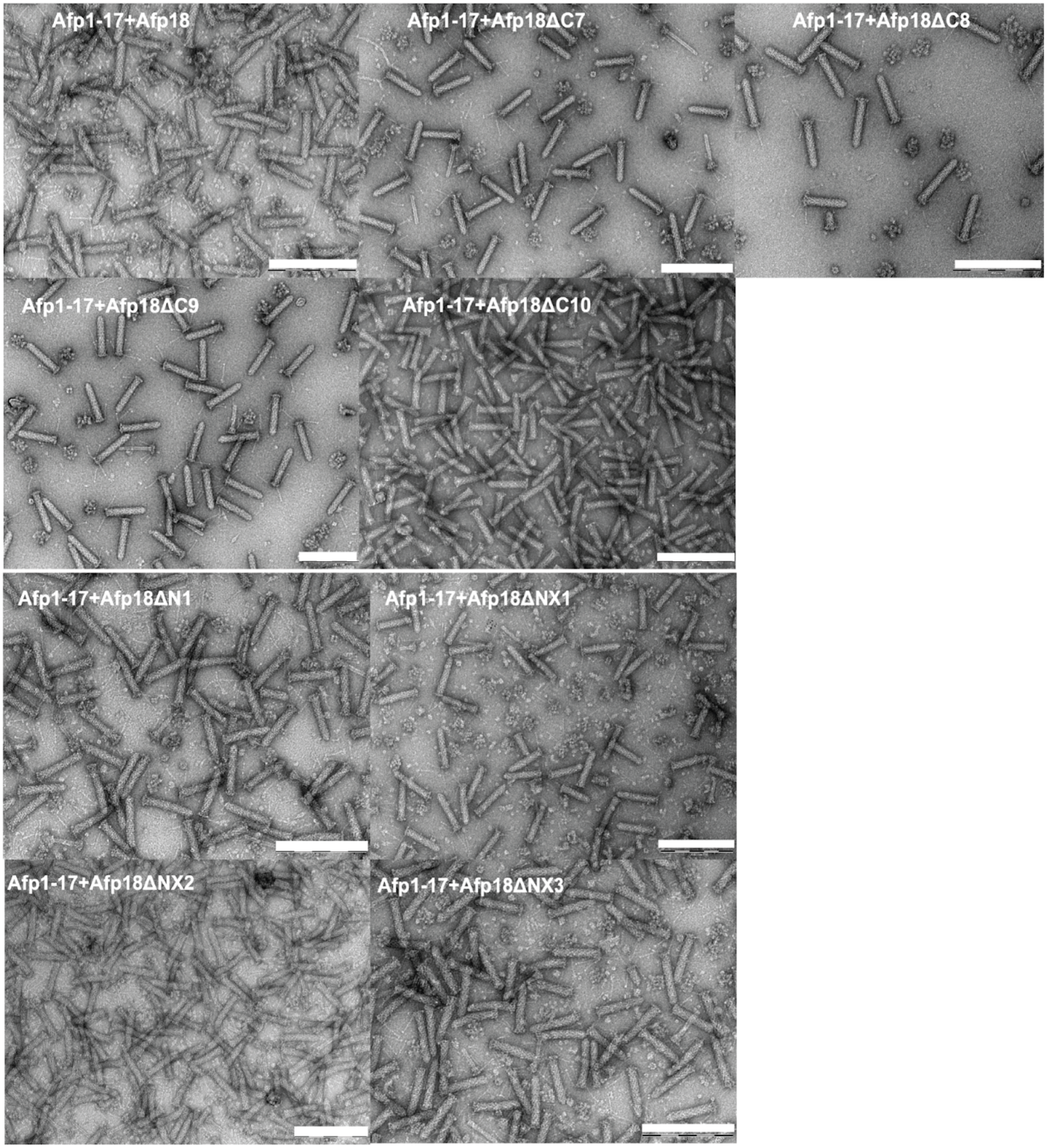


**Fig. S12.**

**Afp18ΔN and ΔCtoxin truncation and particle validation using negative staining EM**
Particles have been further purified after PEG precipitation and UC (Fig. S10, S11) using gradient purification to 'pure' samples qualified for potential cryo-EM analysis.Afp18 toxin N/C-terminal truncations and Afp particle productions and syringe validation using negative staining EM.

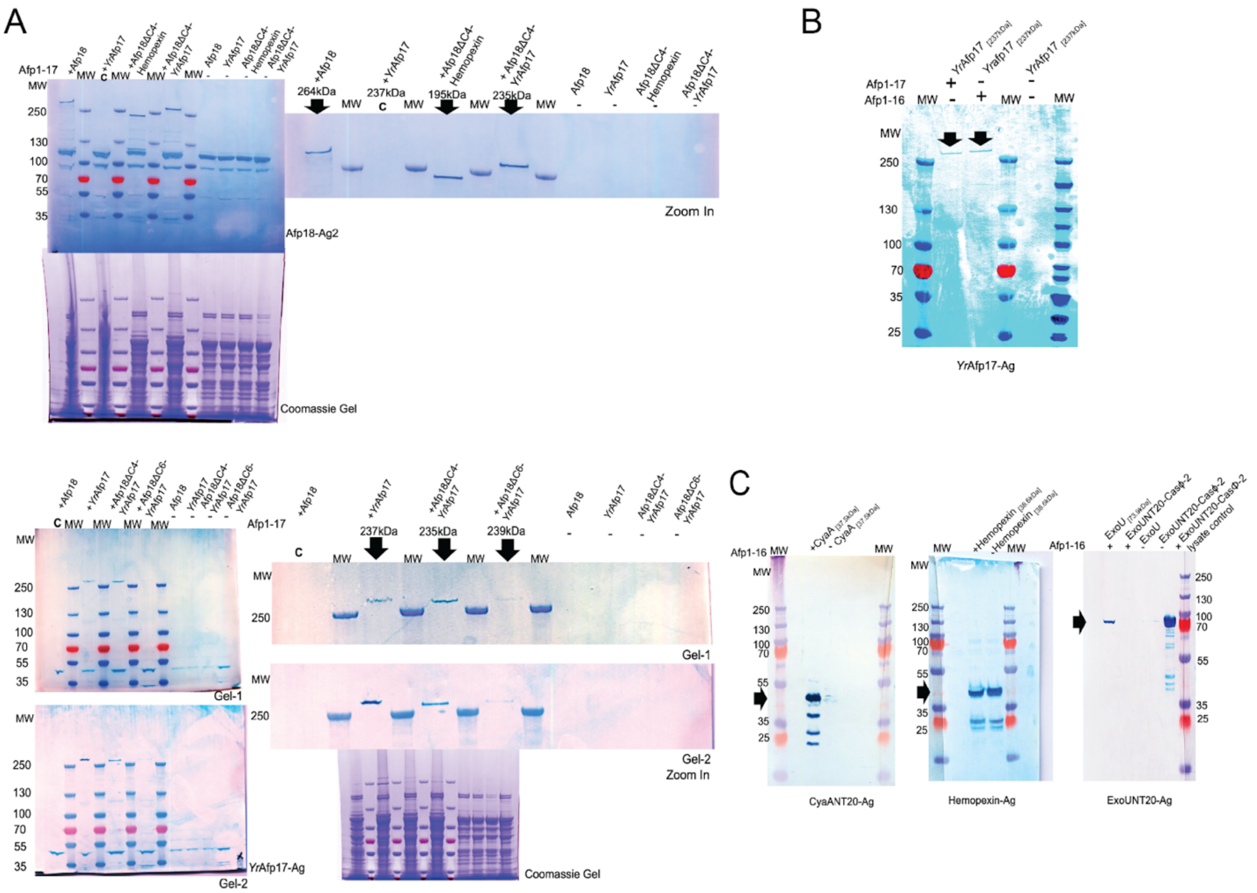

**Fig. S13.**

**Detection of toxins and Afp18ΔC-chimera loading into Afp**

(A) Different Afp18ΔC truncations were used to load toxins of other eCIS. (B) *Y. ruckeri* Afp17, *Yr*Afp17 (Afp18 homologue) was co-purified with Afp and without Afp18 (fragments) as a scaffold. (C) *P. luminescens* toxin CyaA (PluDJC_08830), *P. aeruginosa* ExoU and *P. luminescens* Hemopexin (PluDJC_08520) were co-purified without chimera formation with Afp18, suggesting that similar NtSPs are present on these putative effectors. Note that for hemopexin the mock shows protein being purified without the Afp syringe. Coomassie gels are labeled and immune-detection blots labeled with respective antibodies (Ag) used.

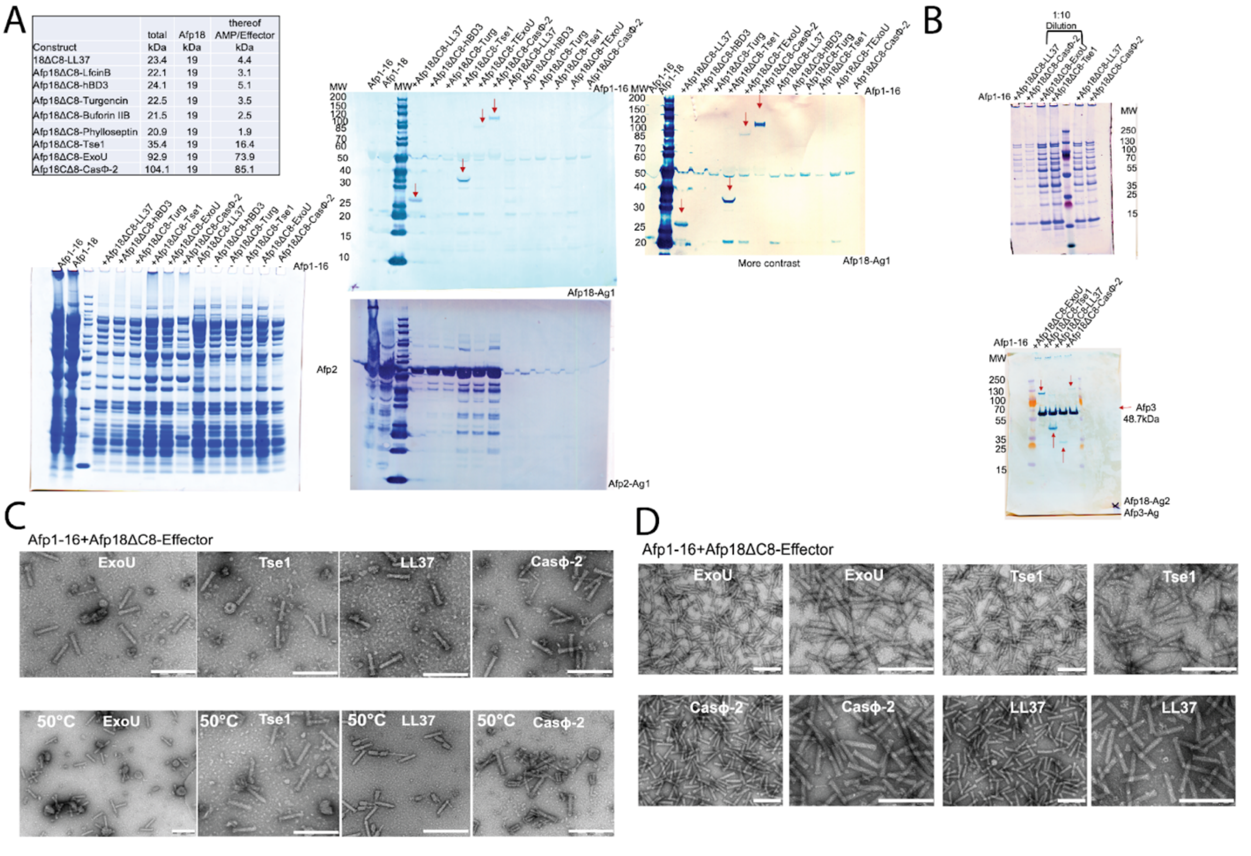


**Fig. S14.**

**Afp18ΔC8-chimera loading into Afp and validation using immune-detection blotting and negative staining EM**

(A) Coomassie gels and immune detection blots of Afp1-6 and Afp18ΔC-effectors (Cathelicidins LL37 hCAP18 human antimicrobial peptide (LL37, α-helical), Lactoferricin B (LfcinB) (β-sheet), human beta-Defensin-3 (hBD3, mixed secondary structure), Phylloseptin (Phyll, α-helical), Buforin II (Bufo, helical-helix-propeller structure) and Turgencin (Turg, α-helical) produced, confirming syringe and effector presence. T6SS effector, Tse1, and T3SS effector ExoU from *P. aeruginosa* were both successfully loaded. As a control, the non-eCIS related toxin-chimeras were produced in parallel (mock expression, no Afp particle) without Afp particle, to exclude false positive results through soluble toxin-chimera aggregates. (B) Coomassie gel and immune detection blot of final 'pure' particle preparations confirming syring and effector presence. (C) Investigation of temperature stability ($T_{stabil}$) of Afp18ΔC8-effector particle preparations after PEG precipitation and ultracentrifugation (UC) revealing particles are intact after 10 min exposure at 50°C (scale bar 200 nm). (D) 'Pure' particle preparations from (B) were investigated for Afp morphology using negative staining EM (scale bar 200 nm).

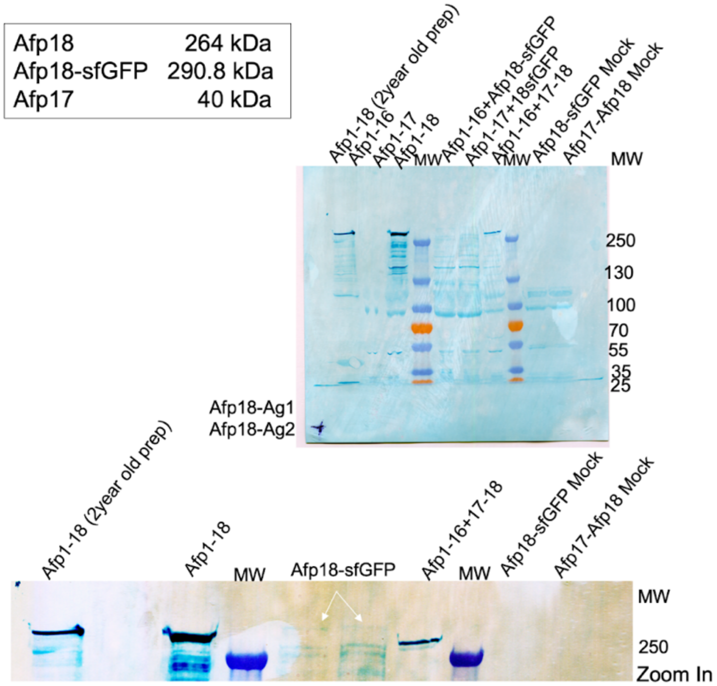


**Fig. S15.**

**Investigation of large cargo packing into Afp**

The largest manipulated cargo tested was Afp18-sfGFP 290.8 kDa shows faint cargo bands when produced together with Afp1-16 and Afp1-17, but not in the mock expression of the toxin only. This indicates that cargo payload could be increased beyond the native 264 kDa Afp18 toxin size. As a stability control the 2 year old particle prep is analyzed and full length Afp18 can be detected in good amounts compared to a fresh Afp1-18 particle preparation.

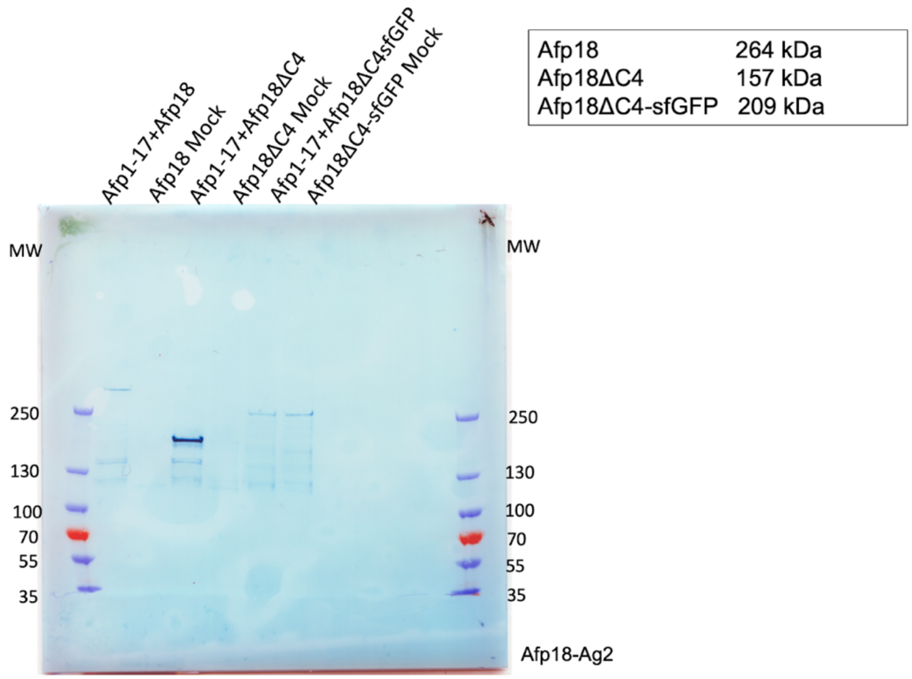


**Fig. S16.**

**Afp particle and Afp18 toxin, Afp18ΔC4 and Afp18ΔC4-sfGFP co-production analysis**

Immune detection blot with Afp18 specific antibodies reveal that attachment of sfGFP to smaller Afp18ΔC4 truncation variant solubilizes the toxin and gives a positive result in mock expression which could be used to solubilize parts of Afp18 for purification and detection.

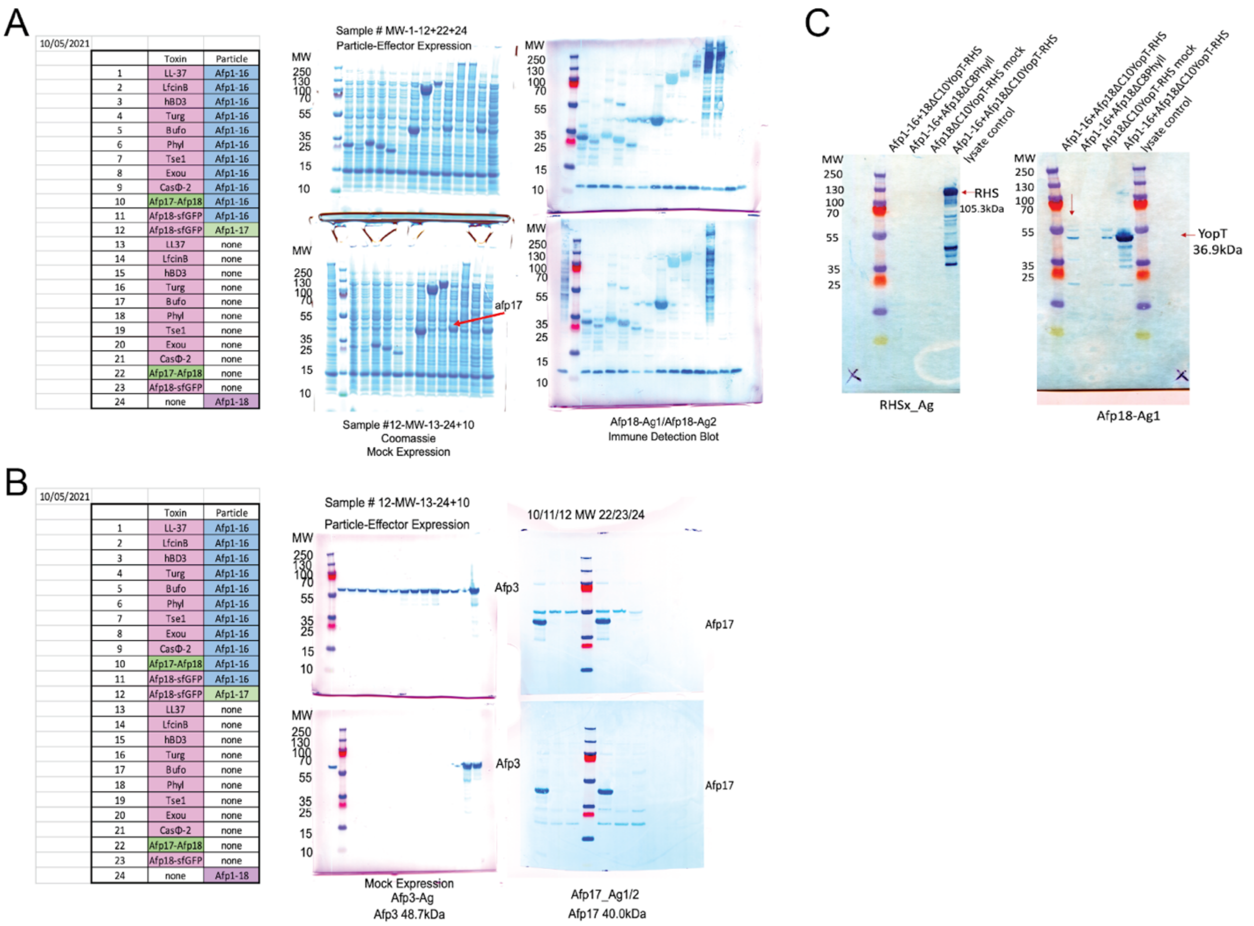


**Fig. S17.**
**Afp and effector-chimera co-expression and mock expression detection**
(A) Afp18ΔC-effectors, Cathelicidins LL37 hCAP18 human antimicrobial peptide (LL37), Lactoferricin B (LfcinB), human beta-defensin-3 (hBD3), phylloseptin (Phyll), buforin II (Bufo) , turgencin (Turg), Biggiephage CasΦ-2, T6SS and T3SS effectors Tse1 and ExoU co-produced with Afp, confirming syringe and effector presence expression levels detected in lysates using Coomassie staining and immune detection blotting. Sample number and co-expression combinations are summarized in the table to the left and loaded sample numbers are indicated above gels (Molecular Weight, MW). Afp18-sfGFP and Afp17-Afp18 were investigated if larger toxins can be packed and if toxin production levels can be increased. (B) Afp18ΔC8-effector expression levels detected in lysates using Coomassie staining and immune detection blotting Afp particle specific antibodies. Sample number and co-expression combinations are summarized in the table to the left and loaded sample numbers are indicated above gels (L, protein ladder). Particles and effectors were expressed, except for Afp18ΔC8-phylloseptin, where expression was not detected at this stage. (C) Immune-detection of purified Afp with YopT-Rhs and phylloseptin as cargo. Rhs was expressed however not co-purified with Afp1-16. Particles with Afp18ΔC8-phylloseptin were further purified to see whether concentrating and purifying particles would show positive cargo loading, however we could not detect Afp18ΔC8-phylloseptin.

| | | |
|---|---|---|
| LL37 AMP | UniProtKB - P49913 (CAMP_HUMAN) | LLGDFFRKSKEKIGKEFKRIVQRIKDFLRNLVPRTES |
| Lactoferricin B (LfcinB) | UniProtKB - P24627 (TRFL_BOVIN) | FKCRRWQWRMKKLGAPSITCVRRAF |
| Human b-defensin-3 (hBD3) | UniProtKB - Q5U7J2 (Q5U7J2_HUMAN) | GIINTLQKYYCRVRGGRCAVLSCLPKEEQIGKCSTRGRKCCRRKK |
| Turgencin B | UniProtKB - C0HLN6 (TURB_SYNTU) | GIKEMLCNMACAQTVCKKSGGPLCDTCQAACKALG |
| Buforin-IIb | UniProtKB - P0DQL2 (H2A_SPHLA) | RAGLQFPVGRLLRRLLRRLLR |
| Phylloseptin-PHa | UniProtKB - A0A2I2KHV2 (A0A2I2KHV2_PITHY) | FLSLIPAAISAVSALANHF |

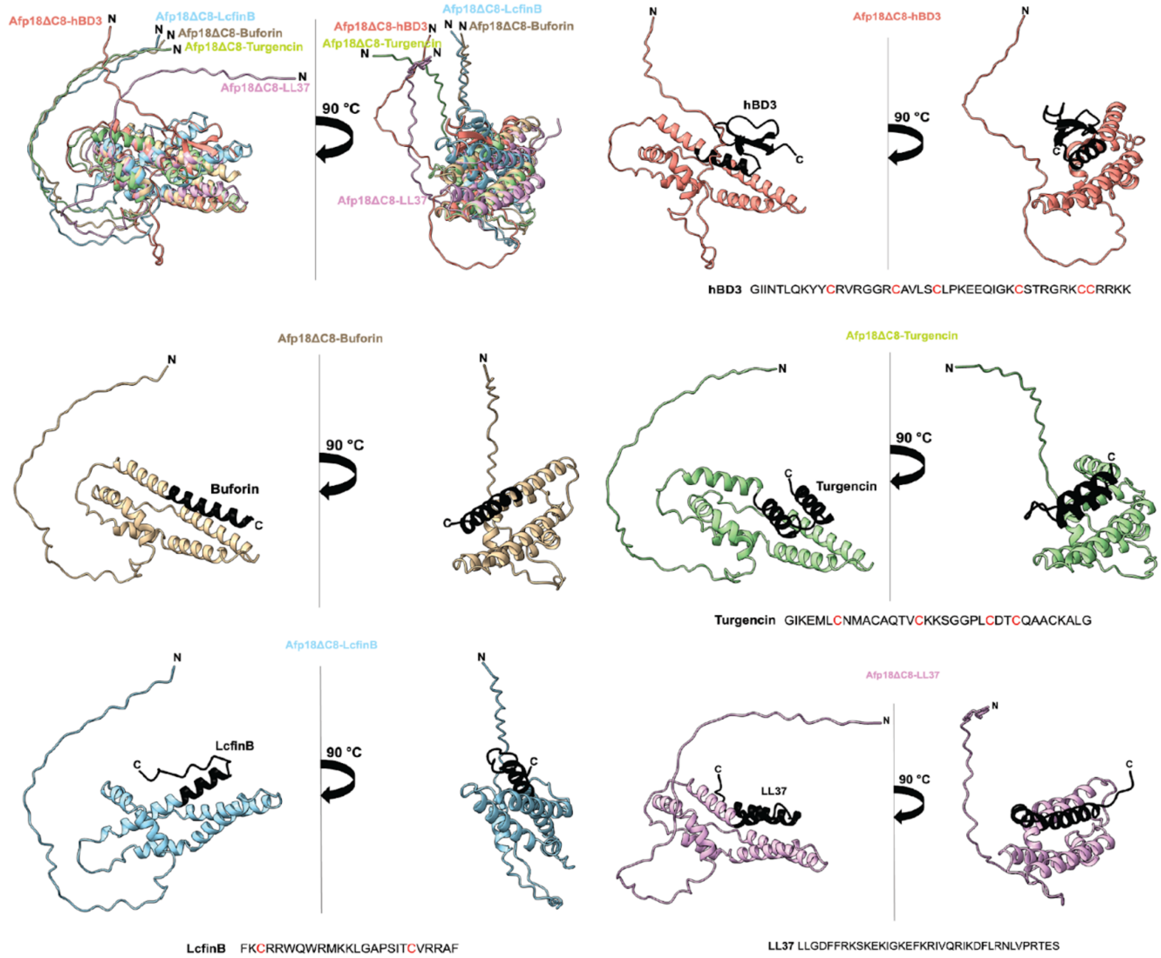


**Fig. S18.**

**Superimposition of Afp18ΔC8-AMP structure prediction using AlphaFold**

Alphafold2 prediction compared to available experimental structural data, AMPs are highlighted in black. All N-terminal regions are predicted to be unstructured and accessible. All AMPs are structured and superposition of all constructs show that cargo appears to have all similar globular size.

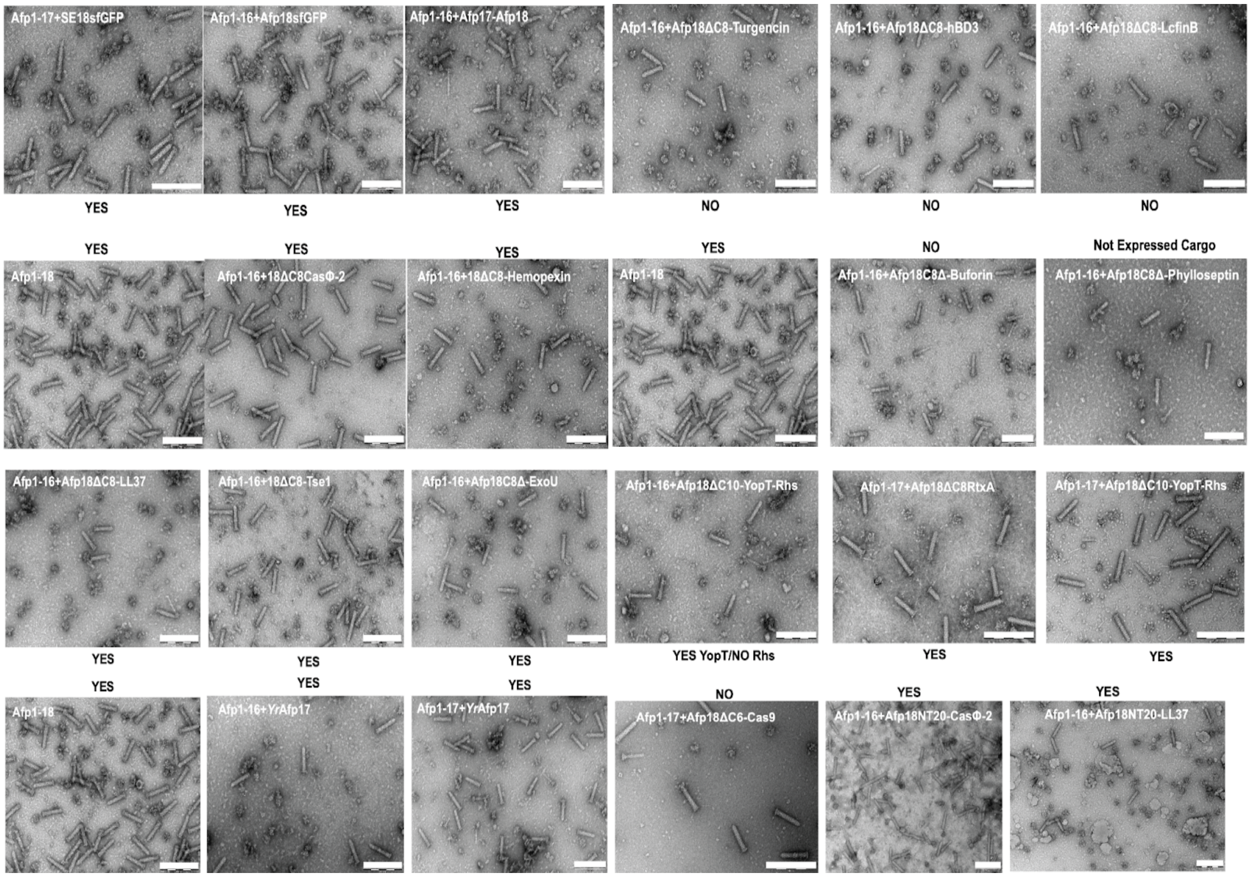


**Fig. S19.**

**Investigation of Afp particle morphology using negative staining EM of Afp particle and effector co-productions**

A 4µl sample of 'pure' particle preparations diluted to similar concentrations (in PBS) was used for negative staining EM and Afp particle morphology investigated. Labels YES/NO highlight whether we could detect effectors in particle preparations.

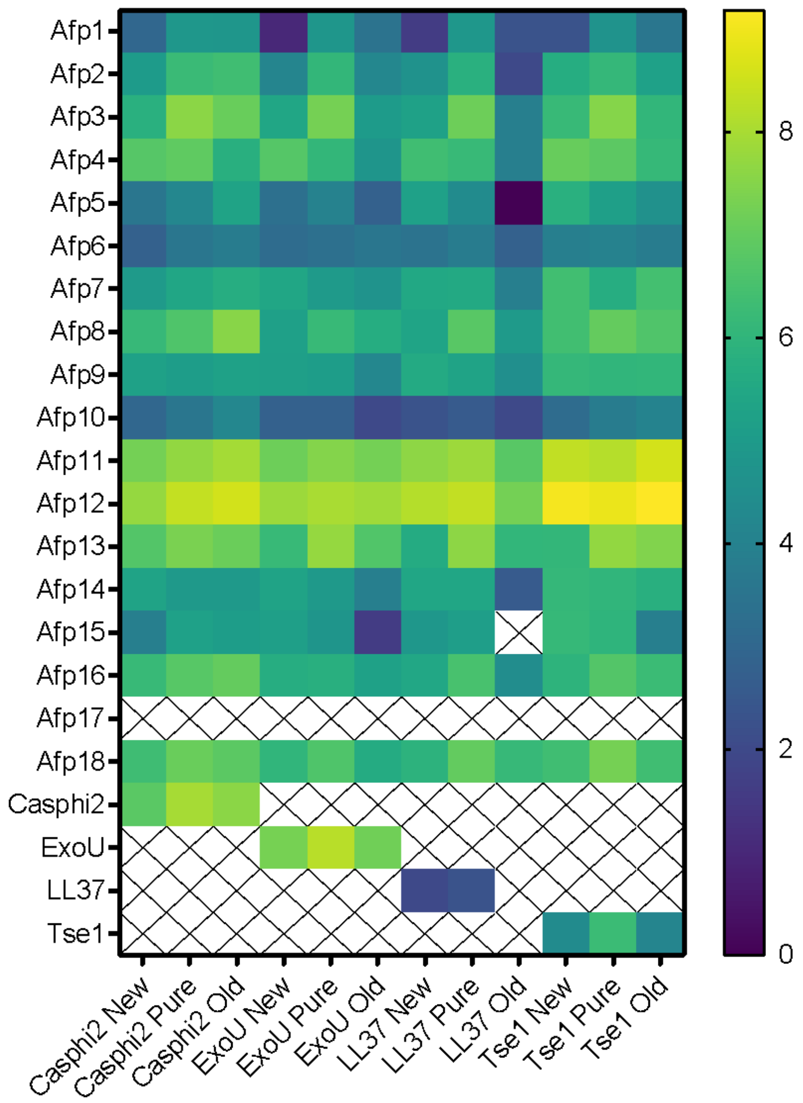


**Fig. S20.**

**Heatmap visualization of mass spectrometry analysis of Afp1-16+Afp18ΔC8-fusion samples**
'New' samples are fresh particle preparations, 'Old' particle preparations that have been stored in the fridge for 4 months prior to analysis (purification protocol '*Co-Expression and Purification of Toxins and Afp particle variants*') and 'Pure' particles which were further purified for cryo-EM analysis ('*Expression and Purification of Afp particles*') with final gradient purification steps. Afp17 was not detected in any of the three particle preparations; crossed out cells indicate no detection. The heatmap visualizes $log_2$-transformed spectral (MS/MS) counts of peptides mapping to each protein, $n$=2 summed technical replicates. In case peptides were homologous to multiple proteins (as entered into the FASTA search space), they were assigned to the protein expected to be present in the sample.

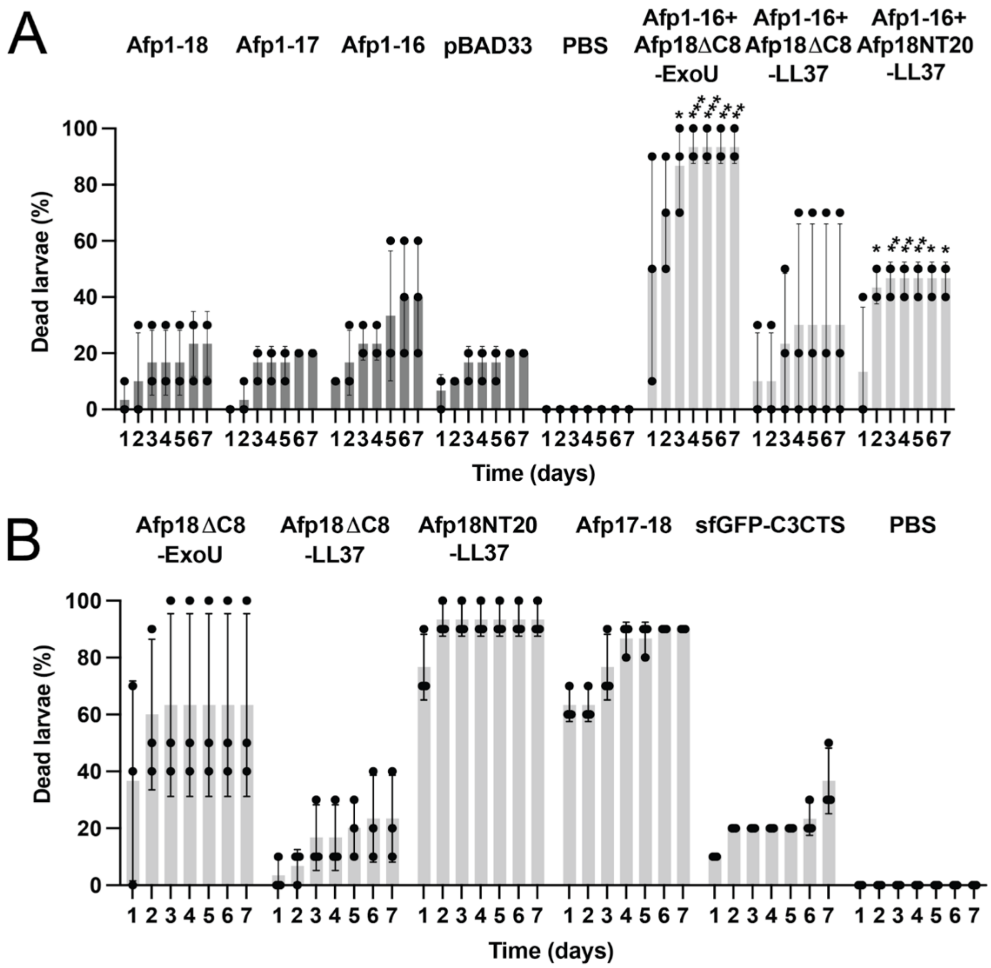


**Fig. S21.**

***In vivo* toxicity tests of Afp and toxin and toxin-effector chimeras on *G. melonella* larvae**

**(A)** Toxicity of *Serratia* Afp variants and Afp with toxin-chimeras. Anti-eukaryotic *P. aeruginosa* T3SS effector ExoU packed as Afp18ΔC8-ExoU and Afp18NT20-LL37 into Afp1-16 particles show significantly increased larvae killing after 1 and 2 days, respectively, as tested by two-way ANOVA with Dunnett multiple testing compared to pBAD33 control. Shown are the individual numbers from each experiment, mean and standard deviation of three independent experiments (n = 3 experiments, 10 larvae in each treatment). **(B)** Toxins and toxin-effector chimera were overexpressed in *E. coli* as toxicity control and cleared lysates injected into the posterior proleg. To ensure that the syringe and toxin components were produced and present in the protein lysate in about the same amounts, SDS-PAGE and immuno-detection against toxins was performed. 10 Galleria mellonella larvae were injected with 30 µl of filtered protein lysates and observed for 7 days. 30 µl of PBS buffer were injected as a control group to ensure that the solution used for the nanoparticle extraction was harmless to the larvae. The construct sfGFP C3CTS (superfolder GFP C-terminal Twin Strep Tag) in pET11a serves as a non-toxic protein expression control.

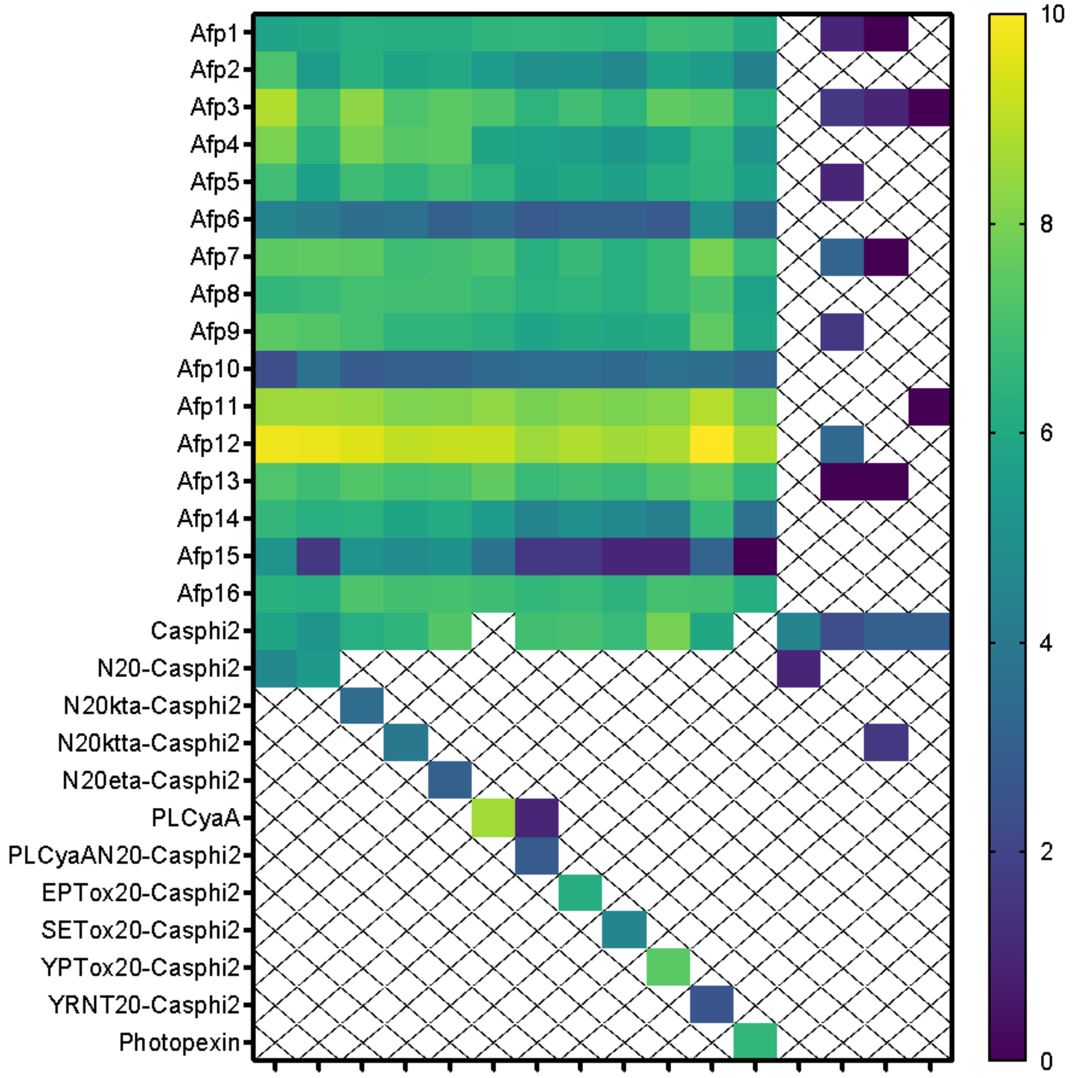


**Fig. S22.**

**Heatmap visualization of mass spectrometry analysis of Afp1-16 and NtSP-CasΦ-2 (Casphi2) fusions**

NtSP (NT20, N-terminal 20 amino acids) representing Afp18NT20 and mutant variants. Crossed out cells indicate no detection. The heatmap visualizes $log_2$-transformed spectral (MS/MS) counts of peptides mapping to each protein, $n$=2 summed technical replicates. In case peptides were homologous to multiple proteins (as entered into the FASTA search space), they were assigned to the protein expected to be present in the sample.

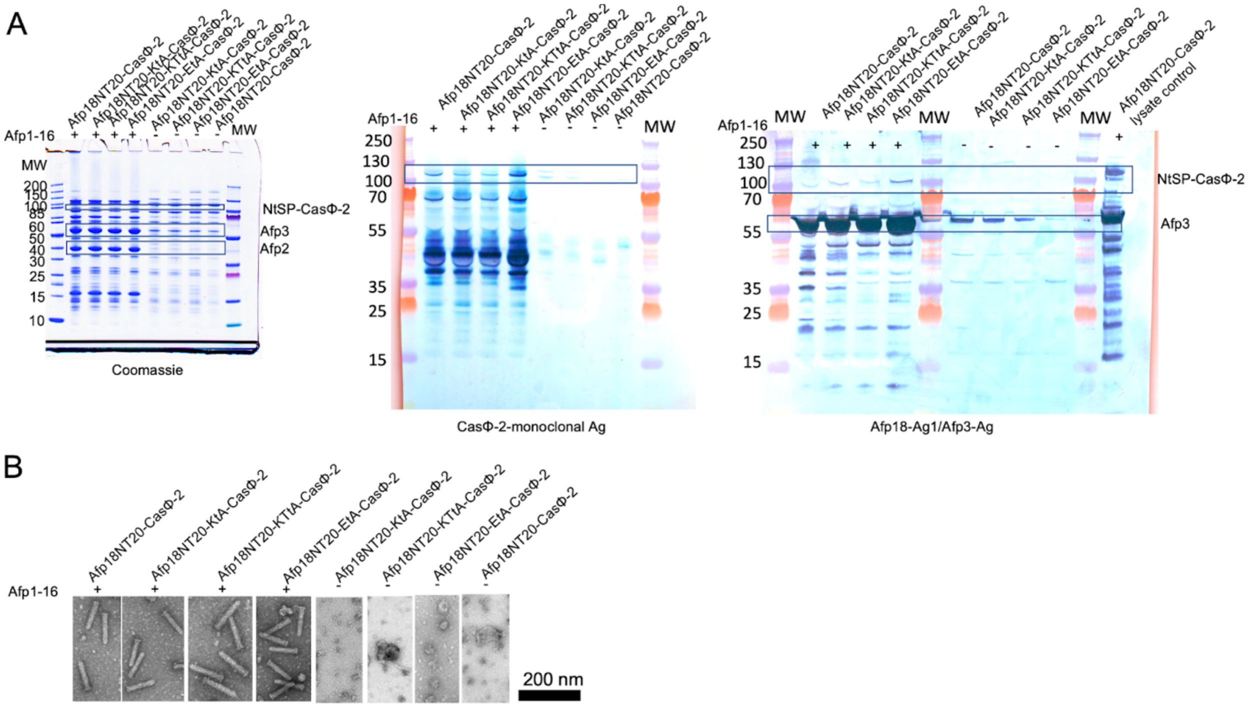


**Fig. S23.**

**Analysis of Afp18NT20 mutants fused to Casϕ-2 and Afp particle preparations**

Mutational analysis of Afp18NT20 N-terminal signal peptide fused to Casϕ-2 and validation of particle, effector components and particle morphology using Coomassie staining, immune detection blotting and negative staining EM. Hydrophilic residues were mutated, lysines to alanine (KtA), lysine and threonine to alanine (KTtA) and glutamic acids to alanine (EtA). (A) Coomassie staining and immune detection blotting showing particle and effector components. (B) Particle morphology was investigated using negative staining EM. All mutation variants still packed CasΦ-2 confirmed by negative staining and immuno-detection blotting (A) and particle morphology was confirmed by negative staining EM (B).

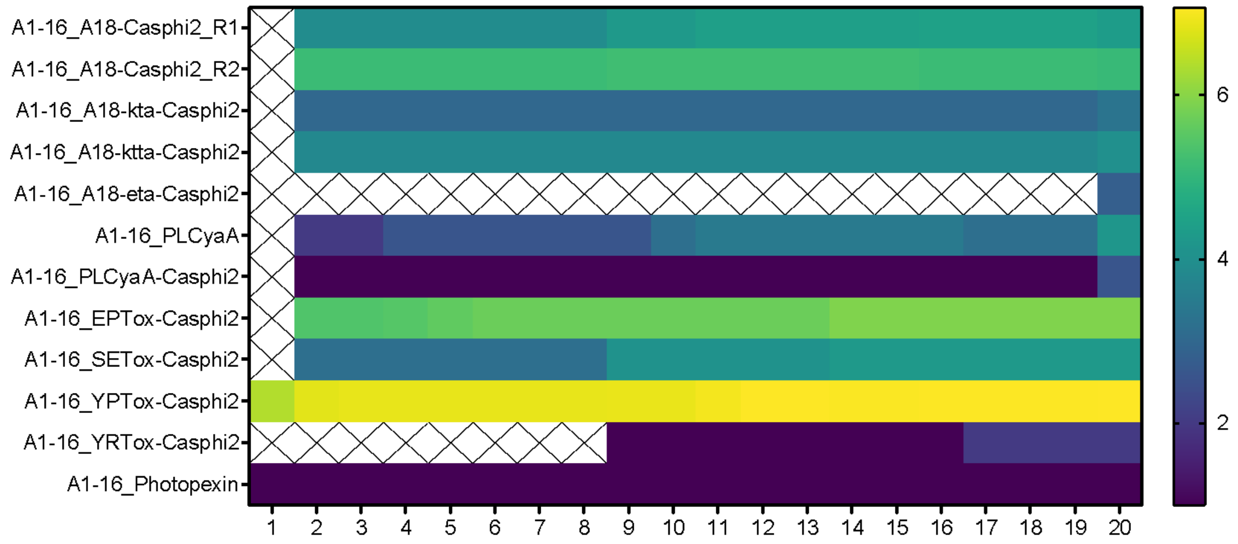


**Fig. S24.**

**Heatmap visualization of N-terminal peptide coverage in NtSP-CasΦ-2 (Casphi2) particle preparations using mass spectrometry**
NtSPs from *P. luminescens* CyaA (PLCyaA, CyaANT20), *Erwinia persicina* (EPTox, *Ep*Tox20), Salmonella enterica (SETox, *Se*Tox20), *Yersinia pekkanenii* (YPTox, *Yp*Tox20), *Yersinia ruckeri* (YRTox, *Yr*Afp17NT20) effectors were present for all particle preparations except for the EtA mutant of Afp18NT20 which was confirmed to be present by immune detection see also Fig. S19. We suspect this peptide has different MS-technical analytical properties, such as hydrophobicity or ionization efficiency, that render it unamenable to detection using standard MS approaches.

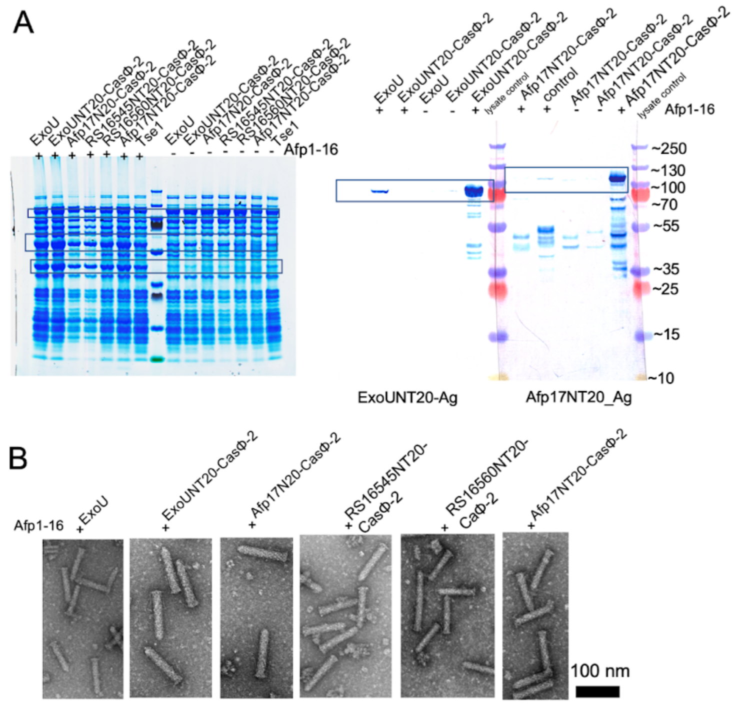


**Fig. S25.**

**Analysis of NtSPs with low hydrophilicity fused to Casϕ-2 and Afp particle preparations**

Two NtSP with low hydrophilic and hydrophobicity Afp17NT20 (ζ of 35%) and ExoUNT20 (ζ of 40%) were shown to not pack CasΦ-2. Two additional N-terminal peptides from *P. asymbiotica* PAU_RS16545 lysozyme N20 (RS16545NT20: MKLSEKGFELIKHFEGLRLH) and PAU_RS16560 LysR transcriptional regulator N20 (RS1656NT20:VFISKELSSFIAVAKNKSIN) (see Table 2) fused to CasΦ-2 were investigated as well as potential non-functional NtSPs. (A) Coomassie staining and immune detection using ExoUNT20 and Afp17NT20 specific antibodies. (B) Negative staining EM showing intact particle morphology for all Afp and effector co-productions.

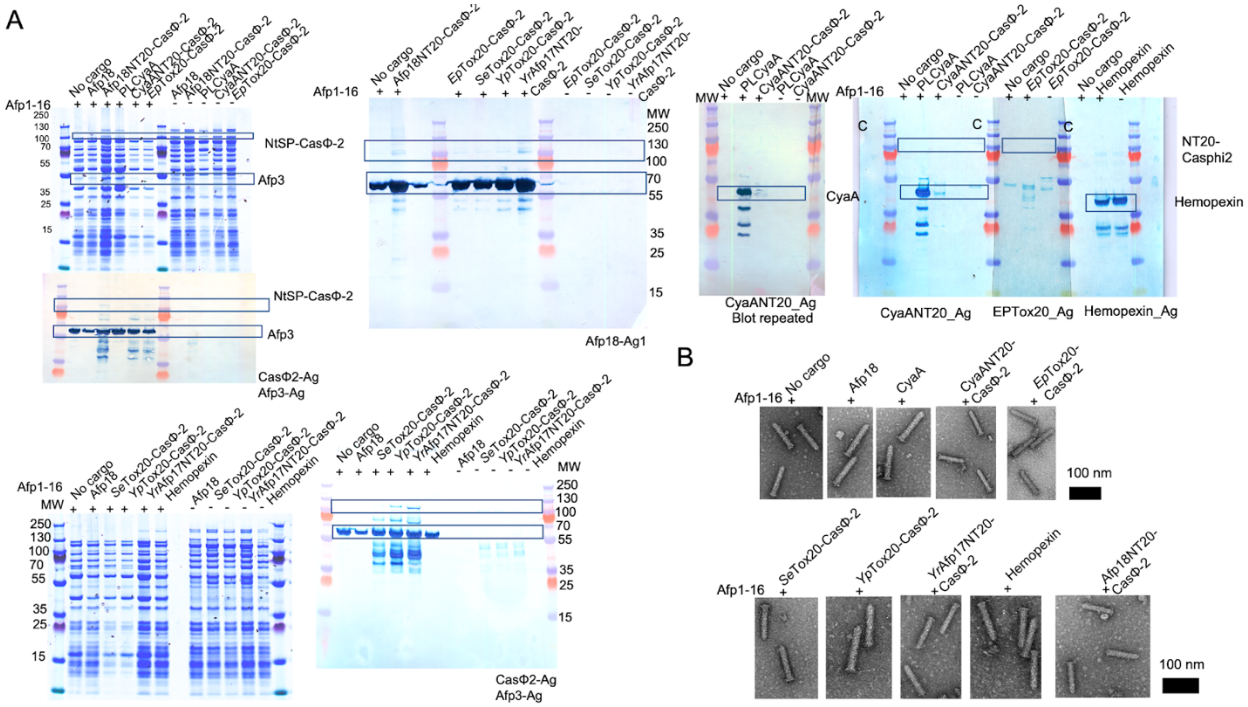

**Fig. S26.**

**NtSPs from Afp18NT20 homologs that pack CasΦ-2 into Afp**

NTSPs from eCIS effectors from other species pack CasΦ-2 into Afp. Effector NtSPs from *P. luminescens* CyaA (CyaA, CyaANT20), *Erwinia persicina* (*Ep*Tox20), *Salmonella enterica* (*Se*Tox20), *Yersinia pekanenii* (*Yp*Tox20), *Yersinia ruckeri* (*Yr*Afp17NT20). (A) Coomassie staining and immune detection with cargo and particle specific antibodies. Particle and effector presence can be validated over immune detection blotting and comparable amounts of particles were produced highlighted using Coomassie staining. (B) Negative staining EM showing intact particle morphology for all Afp preparations.

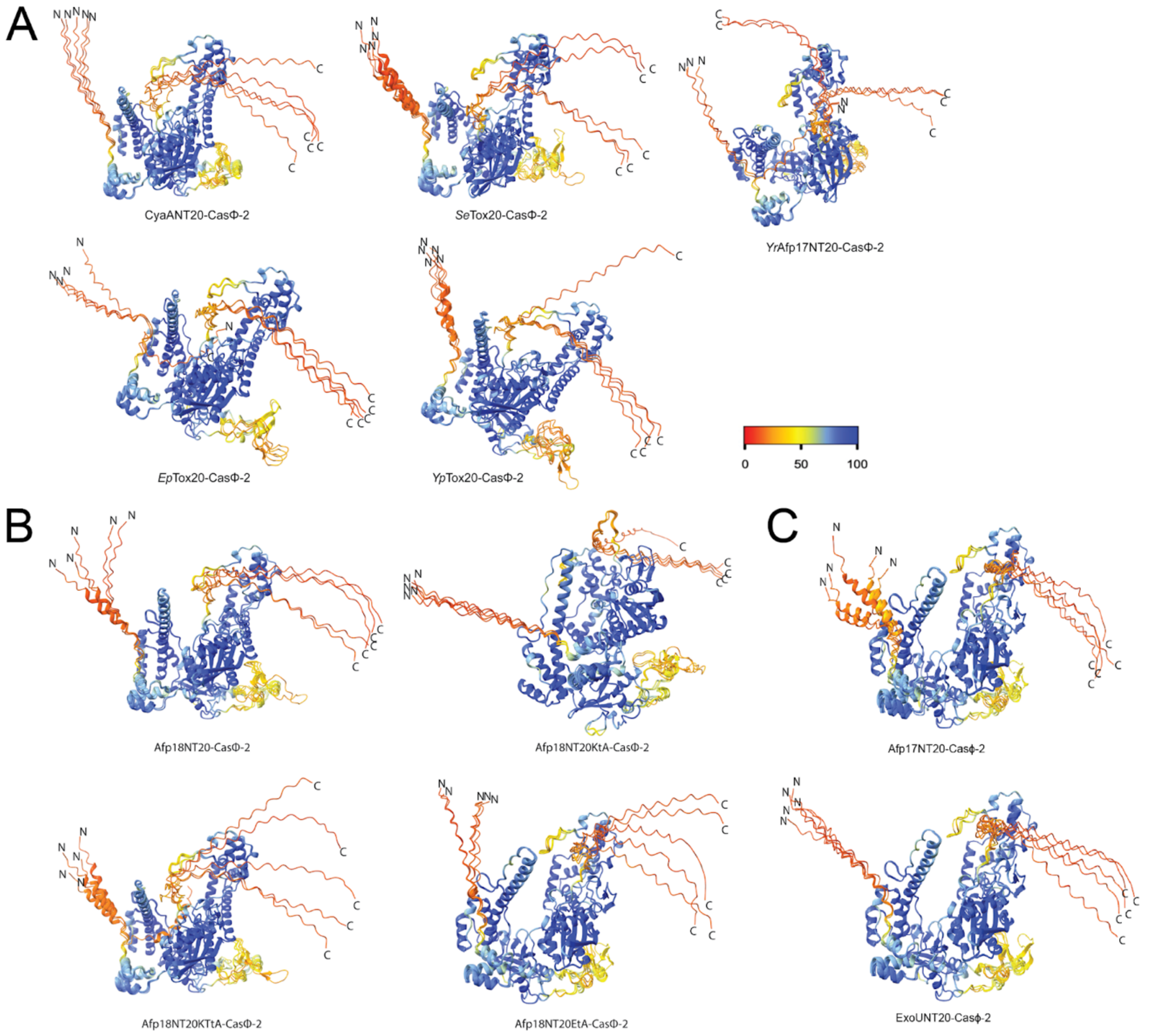


**Fig. S27.**

**Structural predictions of NtSP-CasΦ-2 chimeras**

AlphaFold structural predictions of chimeras of CasΦ-2 fused C-terminally to (A) NtSP from different species, (B) Afp18NT20 and mutant variants and (C) two NtSPs that did not pack fused to CasΦ-2 and their predicted AlphaFold structures. Representation of the 5 best models and colored highlighting model confidence by the pLDDT confidence measure (high confidence in blue and low confidence in red). CasΦ-2 is predicted with high confidence, N-terminal and C-termini are flexible/unstructured, thereby potentially accessible for protein-protein interaction however, predicted with low confidence.

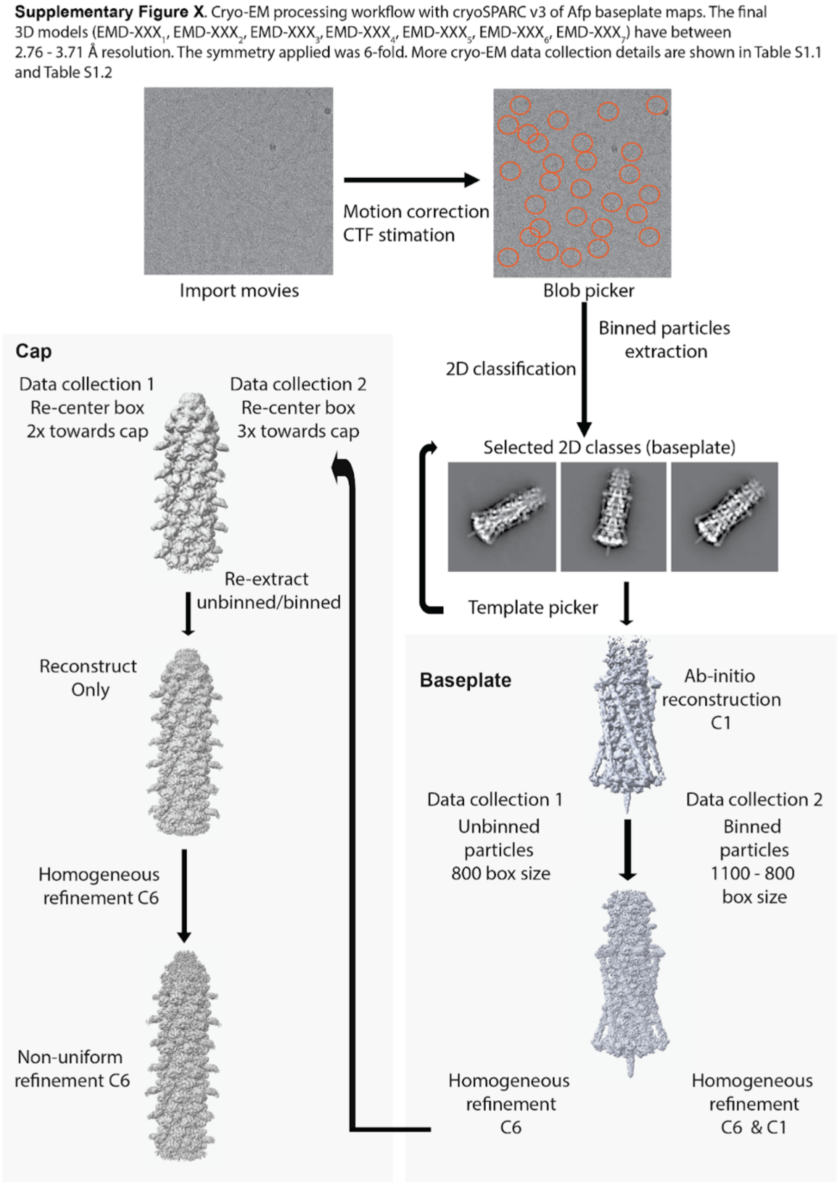


**Fig. S28.**

**Cryo-EM processing workflow with cryoSPARC v3 of Afp baseplate and cap maps**

The final 3D maps of baseplates from data collection 1: Afp1-18 (EMD-18524), Afp1-18ΔC4 (EMD-18526), Afp1-17 (EMD-18551), Afp1-16 (EMD-18552), Afp1-16+Afp18ΔC8-CasΦ-2 (EMD-18525), and from data collection 2: Afp1-16+Afp18ΔC8-CasΦ-2 (EMD-18527), Afp1-16+Afp18ΔC8-ExoU (EMD-18553) in C6 and Afp1-16+Afp18ΔC8-CasΦ-2 (EMD-18528), Afp1-16+Afp18ΔC8-ExoU (EMD-18580) in C1 symmetry, and from Afp-caps in C6 symmetry from data collection 1: Afp1-18 (EMD-18530), Afp1-18ΔC4 (EMD-18531), Afp1-17 (EMD-18575), Afp1-16 (EMD-18576), Afp1-16+Afp18ΔC8-CasΦ-2 (EMD-18532), and from data collection 2: Afp1-16+Afp18ΔC8-CasΦ-2 (EMD-18577), Afp1-16+Afp18ΔC8-ExoU (EMD-18579) in C6. More cryo-EM data collection details are shown in Table S2 and Table S3. Data collection 1 maps were collected on a Titan Krios G2 with a Falcon 3EC direct electron detector and data collection 2 maps on the same microscope updated with a Falcon 4i direct electron detector.

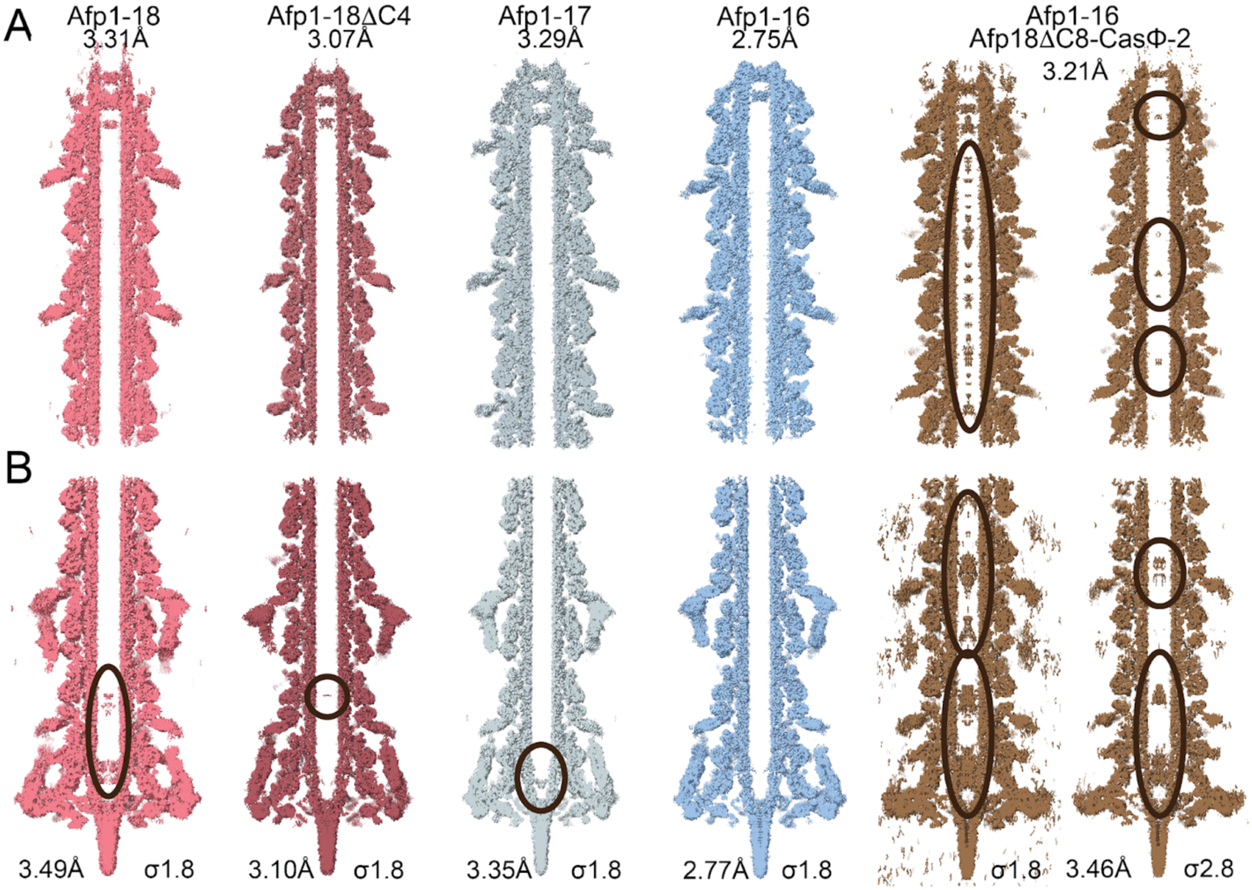


**Fig. S29.**

**Datacollection 1 baseplate and cap cryo-EM maps**

Cap **(A)** and baseplate **(B)** cryo-EM reconstructions of Afp particle preparations. Particle maps are C6 symmetrized. Density along the inside of the tail tube until the cap can be observed for particle preparations Afp1-16 with Afp18ΔC8-Casφ-2. The varying density close to the cap is caused by inhomogeneous length of Afp particles and averaged cap ends at different positions. Cap reconstruction processing details are summarized in Table S2.

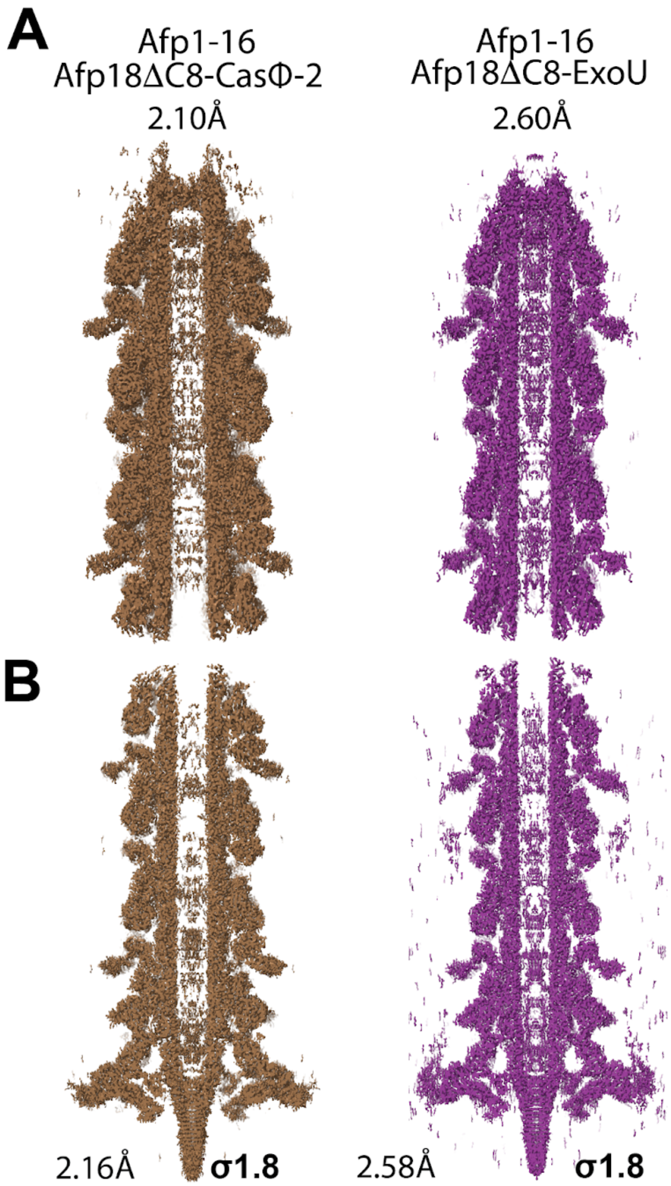


**Fig. S30.**

**Datacollection 2 baseplate and cap cryo-EM maps**

High-resolution cryo-EM maps (C6-symmetrized) of Afp cap and baseplate with two modified cargos Afp18ΔC8-CasΦ-2 (EMD-18577) and Afp18ΔC8-ExoU (EMD-18579) in C6. High-resolution cap (A) and baseplate (B) reconstructions of Afp particle preparations on an updated microscope set-up with a Falcon 4i direct electron detector. Particle maps are C6 symmetrized. Density along the inside of the tail tube until the cap can be observed for particle preparations Afp1-16 with Afp18ΔC8-Casϕ-2 and Afp18ΔC8-ExoU. Cap reconstruction processing details are summarized in Table S3.

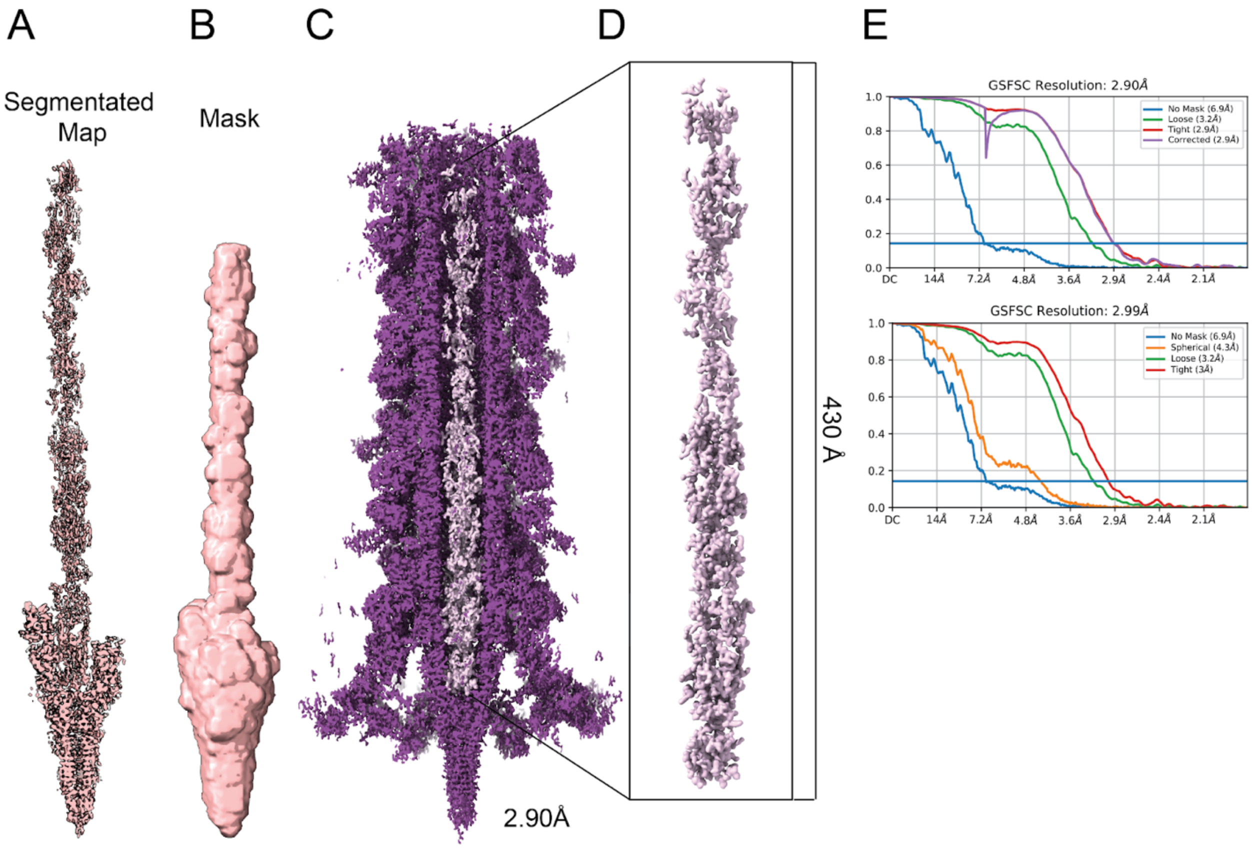


**Fig. S31.**
**Improvement of density inside the Afp tail tube using local refinement**
Local Refinement of Afp1-16 and Afp18ΔC8-ExoU in C1 using a segmentation mask created by segmentation (segmented map) in ChimeraX (A). (B) The input segmented map was processed in cryoSPARC to a mask (threshold 0.2, dilation radius and soft padding 5). (C) The local refinement using a mask of the inner tube density and the central spike from (B) refines to high nominal resolution in C1 (map at sigma 3, purple). The density inside the tail tube is highlighted as a segmentated map (map segmented at sigma 2.6, light pink). (D) Zoom in on the locally refined map inside the tail tube. Sharp features are observed but are not interpretable in terms of an atomic model.. (E) FSC curves generated in cryoSPARC. FSC curve 'corrected' after FSC-mask auto-tightening (top) and 'tight masked' FSC (bottom) of the local refinement in C1 (D).

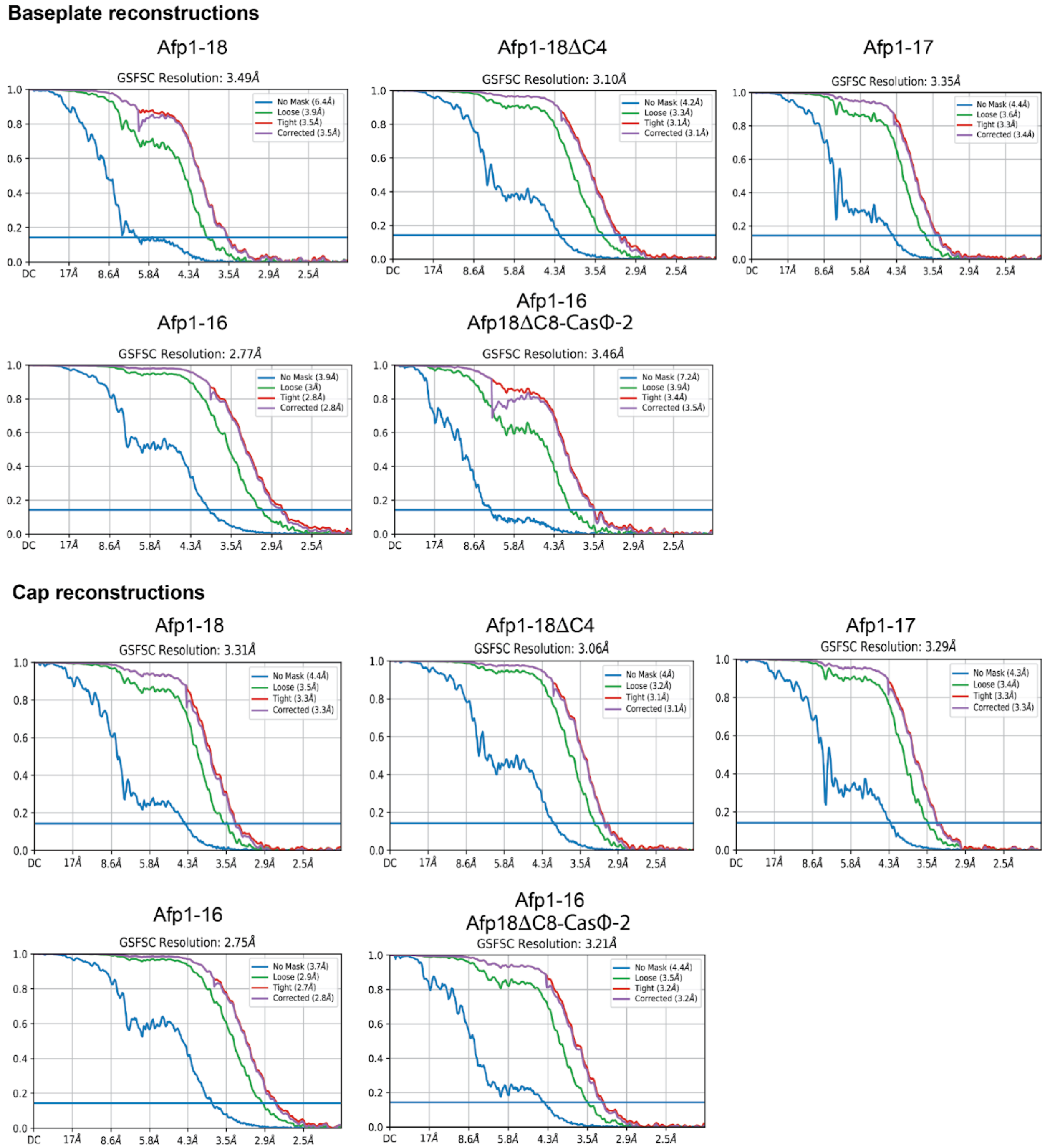


**Fig. S32.**
**Corrected FSC curves from cryoSPARC for cryoEM-maps from updated microscope - Data collection 1**

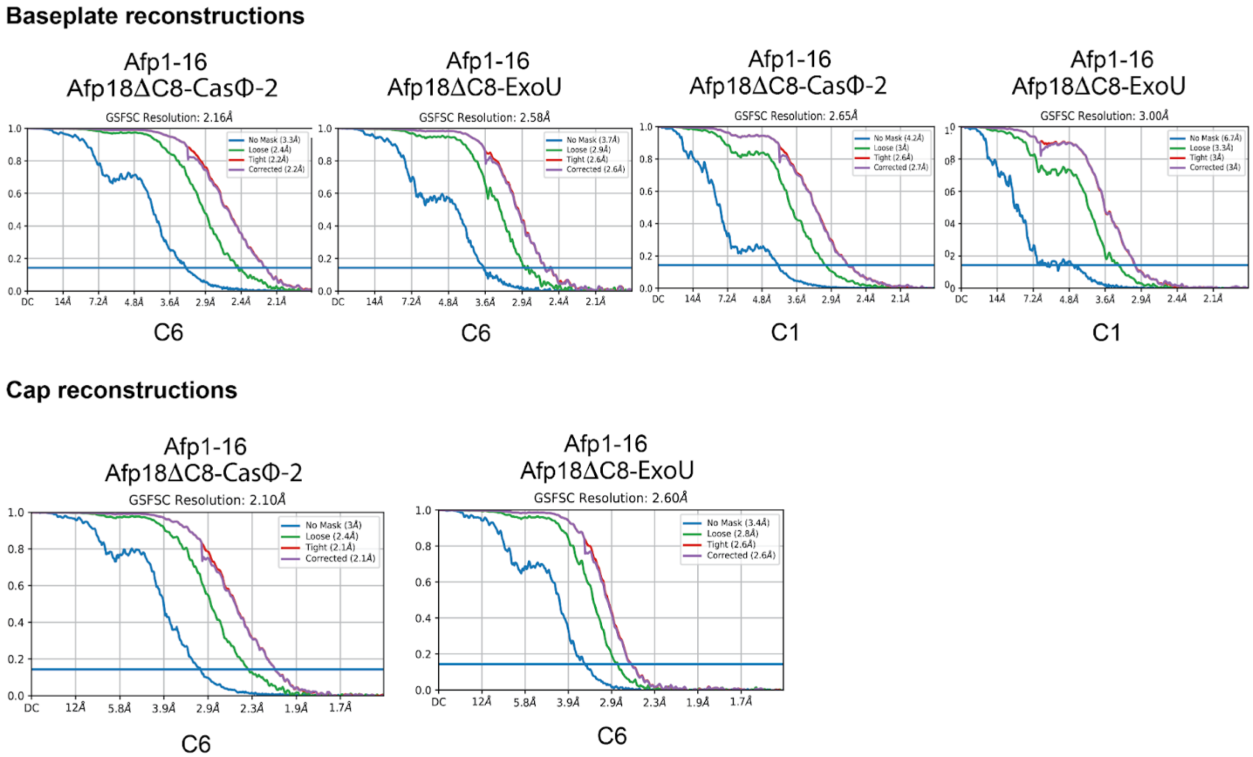


**Fig. S33.**
**Corrected FSC curves from cryoSPARC for cryoEM-maps from updated microscope - Data collection 2**

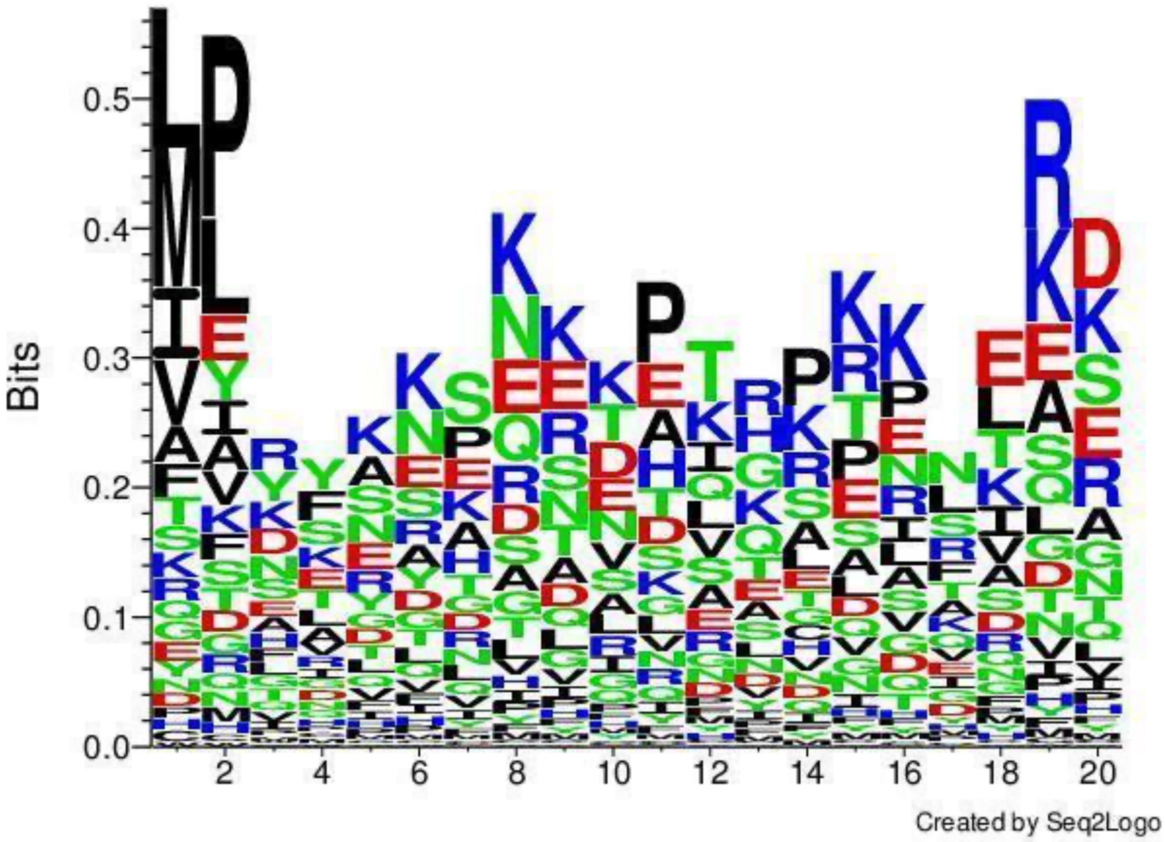


**Fig. S34.**
**Clustal Omega sequence alignment of NtSPs**
Multiple sequence alignment of NtSPs from Table S6 (Clustal Omega) and representation of evolutionary related packing motifs and amino acid preferences/motifs using Seq2Logo, a web-based sequence logo generation method for construction and visualization of amino acid motifs (https://services.healthtech.dtu.dk/service.php?Seq2Logo-2.0). Polar amino acids (lysine (K), glutamic acid (E), asparagine (N), arginine (R) are of high abundance.

| Organism | peptide name | Accession Code | NtSP - peptide | Syringe Gene Locations |
|---|---|---|---|---|
| *Serratia entomophila* | Afp18NT20 | AAT48355.1/ KHA73_24130 | MPYSSESKEKETHSKETERD | afp1 ( AAT48338) – afp18 (AAT48355) |
| *Yersinia ruckeri* | *Yr*Afp17NT20 | DJ39_RS03245 | MPYFNKSKKNEIRPEKSKEE | DJ39_RS03165 – DJ39_RS03245 |
| *Serratia fonticola* | *Sf*Tox20 | WP_021808094.1 | MPYSRESKEKEIHAKETERD | L581_RS22600 – L581_RS22520 |
| *Erwinia persicina* | *Ep*Tox20 | WP_137270131.1 | MPYFNELNEKETRSKETESG | IFT93_RS22375 – IFT93_RS22455 |
| *Yersinia pekkanenii* | *Yp*Tox20 | WP_049612744.1 | MLYSSESKEKKTHSKETERD | AEP37_RS09525 – AEP37_RS09605 |
| *Serratia ureilytica* | *Su*Tox20 | WP_198774613.1 | MPYFRESKEKDTHAKESKQD | JFQ86_16445 – JFQ86_RS16365 |
| *Serratia marcescens* | *Sm*Tox20 | AUO01772.1 | MPYSRESKEKDTHAKGSKQD | C0558_08345 – C0558_08275 |
| *Salmonella enterica* | *Se*Tox20 | HAU3143021.1 | MPYSSESKLKDTHLKEAESD | JDD69_004072 – JDD69_004088 |

**Table S1.**
**Identification of novel NtSPs with Afp18NT20 as search model**
Identification of novel packing motifs in proximity to CIS particles using Afp18NT20: MPYSSESKEKETHSKETERD as input sequence. Detected are NtSPs of other species with high protein homology (above 60%) using BLASTP®.

| | Afp1-18 | Afp1-18ΔC4 | Afp1-17 | Afp1-16 | Afp1-16 +Afp18ΔC8-CasΦ2 |
|---|---|---|---|---|---|
| **Data Collection and Processing** | | | | | |
| Voltage (kV) | 300 | | | | |
| Magnification (nominal) | 75,000 x | | | | |
| Total exposure ($e^-/Å^2$) | 39.83 | 38.24 | 41.22 | 39.67 | 34.53 |
| Pixel size (Å) | 1.08 | | | | |
| Underfocus range (μm) | 0.5 – 2.0 (0.3 steps) | | | | |
| Micrographs used (no.) | 8,813 | 4,574 | 14,780 | 6,589 | 9,875 |
| Final particles (no.) | 49,198 | 74,889 | 58,444 | 130,127 | 24,437 |
| Box size (pixels) | 800 | | | | |
| Symmetry imposed | C6 | | | | |
| Map-sharpened B-factor ($Å^2$) | -129.8 | -136.4 | -148.0 | -121.4 | -117.4 |
| Map resolution (Å) (FSC 0.143) | 3.49 | 3.10 | 3.35 | 2.77 | 3.46 |
| **Cap reconstructions** | | | | | |
| Final particles (no.) | 39,511 | 60,469 | 44,750 | 105,668 | 19,611 |
| Map-sharpened B-factor ($Å^2$) | -122.1 | -128.9 | -130.5 | -114.6 | -109.2 |
| Map resolution (Å) (FSC 0.143) | 3.31 | 3.06 | 3.29 | 2.85 | 3.21 |

**Table S2.**
**Cryo-EM data collection and processing statistics - Data collection 1**

Datasets collected at similar settings on a Titan Krios G2 with a Falcon 3EC direct electron detector and processing details. FSC curves from respective cryoSPARC jobs are listed below.

| | Afp1-16 +Afp18ΔC8-CasΦ2 | Afp1-16 +Afp18ΔC8-ExoU |
|---|---|---|
| Voltage (kV) | 300 | |
| Magnification (nominal) | 165,000 x | |
| Total exposure ($e^-/Å^2$) | 38 | 43 |
| Pixel size (Å) | 0.9 (original 0.725) | |
| Underfocus range (µm) | -0.5 to -2.0 µm | |
| Micrographs used (no.) | 6,877 | 10,142 |
| Final particles (no.) | 61,496 | 26,514 |
| Box size (pixels) | 1100 extracted, binned to 800 | |
| Symmetry imposed | C6/C1 | |
| Map-sharpened B-factor ($Å^2$) | -36.5/-30.5 | -51.3/-28.3 |
| Map resolution (Å) (FSC 0.143) | 2.16/2.65 | 2.58/3.00 |
| **Cap reconstructions** | | |
| Symmetry imposed | C6 | |
| Final particles (no.) | 43,551 | 19,237 |
| Map-sharpened B-factor ($Å^2$) | -29.3 | -49.0 |
| Map resolution (Å) (FSC 0.143) | 2.10 | 2.60 |

**Table S3.**
**Cryo-EM data collection and processing statistics - Data collection 2**
Datasets collected at similar settings on a Titan Krios G2 with a Falcon 4i direct electron detector. FSC curves from respective cryoSPARC jobs are listed below. FSC curves from respective cryoSPARC jobs are listed below

| Primer name | Used for plasmid | Sequence (5′-3′) |
|---|---|---|
| **Primers for pBAD33 Afp syringe cluster versions for native and co-expression** | | |
| pBAD33_lin_BamHI_FW | pBAB33 linearization for plasmids:<br>pBAD33_*afp1-18*<br>pBAD33_*afp1-17* | GGATCCTCTAGAGTCGACCTGC |
| pBAD33_lin_NheI_RV | | GCTAGCCCAAAAAAACGGGTATG |
| afp1_pBAD33_with_RBS_FW | pBAD33_*afp1-18* | ACCCGTTTTTTTGGGCTAGCAGGAGGAATTCACCATGGCTATTACCGCAGACGAC |
| afp_12500_RV_merge | | GCGCGAATAGGTCACCCCAAAGTTGTTAAAGGTGGCG |
| afp_12500_FW_merge | | TTTAACAACTTTGGGGTGACCTATTCGCG |
| afp18_pBAD33_RVlong | | AGGTCGACTCTAGAGGATCCTTAACCAAACACATTCCTCCACAAGATAAAC |
| SE_Afp1-17_pBAD33_RV | pBAD33_*afp1-17* | AGGTCGACTCTAGAGGATCCTTAGTTACACTCTGACCTAAATACTTGACAGAGC |
| SEAfp16_pBAD33OH_F1_RV | pBAD33_*afp1-16* | GGTCGACTCTAGAGGATCCTTAAATACCAGTCAGGTCTGTTTTATCAACCG |
| SEAfp8-9_merge_F1_FW | | CCTATTACGGTGAGGTTGCTATGAGTAACG |
| pBAD33_SEAfp16OH_F2_FW | | GGTATTTAAGGATCCTCTAGAGTCGACCTGCAGG |
| SEAfp8-9_merge_F2_RV | | GCAACCTCACCGTAATAGGTTATTTTTTCATGTTAATTTTCGC |
| SEAfp11_merge_FW | pBAD33_afp1-18ΔC4 | GTAACACCGGGCAATCAGGTGG |
| SEAfp11_merge_RV | | ACCTGATTGCCCGGTGTTACCG |
| SEAfp18C4trunc_pBAD33OH_RV | | CTAGAGGATCCTTATATATGATGTTCGGTCGTCTGGTCAAAAGTC |
| pBAD33OH_SEAfp18C4_FW | | GAACATCATATATAAGGATCCTCTAGAGTCGACCTGC |
| **Primers for pET11a cargoO co-expression plasmids** | | |
| pET11a_C3CTS_lin_FW | pET11a_*afp18_C3CTS* | GGTACCCTTGAGGTGCTG |
| p11a_01_RV | | ATGTATATCTCCTTCTTAAAGTTAAACAAAATTATTTCTAGAGG |
| Afp18_pET11a_C-3CTS_FW | | TTTAAGAAGGAGATATACATATGCCTTACTCCAGTGAGTCGAAGG |
| Afp18_pET11a_C-3CTS_RV | | AACAGCACCTCAAGGGTACCACCAAACACATTCCTCCACAAGATAAACATTAGTAATG |
| pET11a_C3CTS-Stop_lin_FW | pET11a_*afp18* | TAAGGTACCCTTGAGGTGCTGTTTCAGG |
| SE_Afp18_p11a_Stop_RV | | TTAACCAAACACATTCCTCCACAAGATAAACATTAGTAATG |
| pET11a_C3CTS-Stop_lin_FW | pET11a linearization primers for plasmids:<br>pET11a_*yrafp17*<br>pET11a_*hemopexin-PluDJC_08520*<br>pET11a_*afp17-afp18* | TAAGGTACCCTTGAGGTGCTGTTTCAGG |
| NEWpET11aLinYRAfp17tox_RV | | ATGTATATCTCCTTCTTAAAGTTAAACAAAATTATTTCTAGAGGGGAATTGTTATCC |
| sfGFP_SEAfp18OH_FW | pET11a_*afp18_sfgfp* | TGTGGAGGAATGTGTTTGGTATGCGTAAAGGCGAAGAGCTGTTC |
| sfGFP_SEAfp18OH_RV | | AGCACCTCAAGGGTACCTTATTTGTACAGTTCATCCATACCATGCG |
| pET11a_C3CTS-Stop_lin_FW | | TAAGGTACCCTTGAGGTGCTGTTTCAGG |
| afp18_nostop_RV | | ACCAAACACATTCCTCCACAAGATAAACATTAGTAATG |
| SEAfp17_OHpET11a_FW | pET11a_*afp17-afp18* | CTTTAAGAAGGAGATATACATATGCCGACTAAAACACCACAGTTACAG |
| SEAfp18_stop_pET11a_RV | | AGCACCTCAAGGGTACCTTAACCAAACACATTCCTCCACAAGATAAACATTAGTAATG |
| Yr_Tox_p11aC3CTS_FWlong | pET11a_*yrafp17* | CTTTAAGAAGGAGATATACATATGCCTTACTTTAATAAATCGAAGAAAAATGAAATAAGG |
| YR_Afp17_p11a_stop_RVpubl | | AACAGCACCTCAAGGGTACCTTAGCTCAGGCCTGAGCTTTGAGTGCTGTTAC |
| Photopexin_OH-for-pET11a_FW | pET11a_*hemopexin-PluDJC_08520* | GTGGACAGCAAATGGGTCGCATGAATATATCTAGTTATTTCTTCC |

| | | |
|---|---|---|
| Photopexin_OH-for-pET11a_RV | | GCTTTGTTAGCAGCCGGATCCTTATGTCAGAGGCCAATTCAAGAAC |
| SEAfp18C4_RV | pET11a_*afp18ΔC4_yr*<br>pET11a_*afp18ΔC4_hemopexin_PluDJC_08520*<br>pET11a_afp18ΔC4_sfgfp | TATATGATGTTCGGTCGTCTGGTCAAAAGTC |
| pET11a_C3CTS-Stop_lin_FW | | TAAGGTACCCTTGAGGTGCTGTTTCAGG |
| YRAfp17_aa1437_OHSEAfp18C4_FW | pET11a_*afp18ΔC4_yr* | CAGACGACCGAACATCATATAATGCGTATCGCTAAGATGTATACCCCAAC |
| YR_Afp17_p11a_stop_RVpubl | | AACAGCACCTCAAGGGTACCTTAGCTCAGGCCTGAGCTTTGAGTGCTGTTAC |
| PLDJCVC1_08520_OHSEAfp18C4_FW | pET11a_*afp18ΔC4_hemopexin_PluDJC_08520* | CAGACGACCGAACATCATATAATGAATATATCTAGTTATTTCTTCCTAAATGAAGAAAAC |
| PLDJCVC1_08520_OHpET11a_RV | | AGCACCTCAAGGGTACCTTATGTCAGAGGCCAATTCAAGAACACC |
| sfGFP_OHAfp18C4_FW | pET11a_*afp18ΔC4_sfgfp* | CCAGACGACCGAACATCATATAATGCGTAAAGGCGAAGAGCTGTTC |
| sfGFP_OHAfp18C4_RV | | AGCACCTCAAGGGTACCTTATTGAAGCTGCCACAAGGCAGGAAC |
| pET11a_C3CTS-Stop_lin_FW | pET11a_*afp18ΔC6_Yr*<br>pET11a_ *afp18ΔC6_sfgfp*<br>pET11a_*afp18ΔC6_cas9* | TAAGGTACCCTTGAGGTGCTGTTTCAGG |
| SEAfp18C6_RV | | TTTCAGGACAGCAACCTGCTGC |
| YRAfp17_aa502_OHSEAfp18C6_FW | pET11a_*afp18ΔC6_yr* | AGCAGGTTGCTGTCCTGAAAATGTCAGCATTGAGAGATGAATTAAAACGTGC |
| YR_Afp17_p11a_stop_RVpubl | | AACAGCACCTCAAGGGTACCTTAGCTCAGGCCTGAGCTTTGAGTGCTGTTAC |
| sfGFP_OHAfp18C6_FW | pET11a_*afp18ΔC6_sfgfp* | AGCAGGTTGCTGTCCTGAAAATGCGTAAAGGCGAAGAGCTGTTC |
| sfGFP_OHAfp18C6_RV | | AGCACCTCAAGGGTACCTTATTGAAGCTGCCACAAGGCAGG |
| Cas9_SE18C6OH_FW | pET11a_*afp18ΔC6_cas9* | GCAGGTTGCTGTCCTGAAAATGGATAAGAAATACTCAATAGGCTTAGATATCGGC |
| Cas9_pET11aOH_RV | | AGCACCTCAAGGGTACCTTATCAGTCACCTCCTAGCTGACTCAAATC |
| pET11a_C3CTS-Stop_lin_FW | pET11a_*afp18ΔC8_rtxA-PluDJC_12685* | TAAGGTACCCTTGAGGTGCTGTTTCAGG |
| SEAfp18C8_RV | | GGTTGACTCCAACCGTCTGGC |
| PLDJCVC5_RtxA_OHSE18C8_FW | | CCAGACGGTTGGAGTCAACCATGGTATATGAATACGATAAAACCATCGAAAG |
| PLDJCVC5_RtxA_pET11aOH_RV | | AGCACCTCAAGGGTACCTTAAGATGTTAATTGAATACGGGGTAATTCAAC |
| pET11a_C3CTS-Stop_lin_FW | pET11a_*afp18ΔC10_yopT-rhs-PAU_RS10125-20* | TAAGGTACCCTTGAGGTGCTGTTTCAGG |
| SEAfp18C10_RV | | GCTGACCCGTTGAAATACTGGGTC |
| PAlopT_Yoptox_SE18C10OH_FW | | CAGTATTTCAACGGGTCAGCATGGAACGTGAATATAATAAGAAAGAAAAACAGAAAAAGTC |
| PAlopT-RHStox_OHpET11a_RV | | AGCACCTCAAGGGTACCTTACCGTCTTTCAGGGGGCCTG |
| PLVC1_08560-13_OHSE12_FW | pBAD33 *afp1-16Δ13_PluDJC_08560_*fibre | TGACGAATCTGGAGAGCAAAATGGTAAAAAAATATAACCTATATAGTTCAGACG |
| SEAfp12_end_RV | | TTTGCTCTCCAGATTCGTCATGATG |
| SEAfp14_FW | | ATGACTAGAAATTTTTTATATACTCCCAGTATTACTATACAAAAGTG |
| PLVC1_08560-13_OHSE14_RV | | GGGAGTATATAAAAAATTTCTAGTCATTAGAGTTTCATAATAAAAGCTAAAATATAATAAGGC |
| pBAD33_mergeF1-2_RV | | CAGTGACGGCAATGTCTGATGC |
| pBAD33_mergeF1-2_FW | | ATCAGACATTGCCGTCACTGC |
| SE_mCherryFrag1_w/OHtomCherryLin_FW | pBAD33 *afp1-16*_mCherry-*afp3* | ATTCACTGAAGGAAAAGAACTTTGTTTAACTTTAAGAAGGAGATATAGATATCC |
| YRFull_mCherry_Linker_RV | | GAATTCACCAGAACCCGCCGCAGAACCCGCAGAACCCTTGTACAGCTCGTCCATGC |
| SE_mCherryFrag1_w/OHtomCherryLin_FW | | CGGCGGGTTCTGGTGAATTCATGGCTACTGTCACATCTGTACCGGGTG |

| | | |
|---|---|---|
| SE_mCherry_w/OHtoAfp13_RV | | AAGTAACTGCTGTACACATAGAAAGTGGCATTATGAAGTCCACACATCACG |
| SE_mCherry_LinFrag_FW | | TATGTGTACAGCAGTTACTTTTGCGCATTC |
| SEAfp2_AAW_RV | | GTTCTTTTCCTTCAGTGAATTATTGCGCGATATCC |

**Table S4.**
**List of primers used to produce plasmids used in this study**

| Antibody name | Target protein | Antigenic determinant sequence |
|---|---|---|
| Afp18-Ag1 | *S. entomophila* Afp18 toxin | SSESKEKETHSKETC |
| Afp18-Ag2 | *S. entomophila* Afp18 toxin | RVRKRAVADMAPPPPHQWC |
| Afp18-Ag3 | *S. entomophila* Afp18 toxin | CSSKFKSSDEFKVDK |
| Afp18-Ag4 | *S. entomophila* Afp18 toxin | IIESKNSLDHDTSYC |
| Afp18-Ag5 | *S. entomophila* Afp18 toxin | CPPTRDSRYFYENER |
| Afp18-Ag6 | *S. entomophila* Afp18 toxin | CNYKVREKDNLEKKL |
| Afp17-Ag1 | *S. entomophila* Afp17 toxin | DPSFTITKTNQDAGC |
| Afp17-Ag2 | *S. entomophila* Afp17 toxin | SFSLRERDEHESGYC |
| Afp2-Ag1: | *S. entomophila* Afp2 sheath protein | TTYPGVYLSEDAVSC |
| Afp3-Ag1: | *S. entomophila* Afp2 sheath protein | DSNVPSSARVTVSSTAVEC |
| *Yr*_Afp17_Ag | *Y. ruckeri* Afp17 toxin protein | DHSWVTDMPGNSTQSC |
| *Ep*_Tox20_Ag | *Erwinia persicina* N-terminal signal domain | FNELNEKETRSKETC |
| CyaANT20_Ag | *P. luminescens* CyaA toxin N-terminal signal domain | YSNSQRTPTQSTKNC |
| ExoUNT20_Ag | *P. aeruginosa* ExoU Type III Secretion system effector N-terminal signal domain | ATASSLNQEPVETC |
| Afp18N20KtA_Ag | *S. entomophila* Afp18 toxin N-terminal signal domain lysine (K) to alanine (A) mutant | SSESAEAETHSAETC |
| Afp17NT20_Ag | *S. entomophila* Afp17 pseudotoxin N-terminal domain | KTPQLQLAIEEFNKC |
| RHSx_Ag | *P. asymbiotica Rhs repeat protein PAU_RS10120* | SADPAGTVDGLNLYRC |
| Hemopexin_Ag | *P. luminescens PluDJC_08520 Hemopexin homology protein* | DTDLLGSNRDNSGGC |

**Table S5.**
**List of polyclonal rabbit antibodies (Ag) used in this study**

| Organism | gene name/peptide name | peptide | aa distribution using protparam | Total polar aa content (%) | CC(1,3) (lag=2) |
|---|---|---|---|---|---|
| *Yersinia ruckeri* | *Yr* Afp17NT20 | MPYFNKSKKNEIRPEKSKEE | 5% (R),10% (N), 20% (E), 25% (K), 10% (S), 5% (Y) | 75 | -1,516 |
| *Serratia entomophila* | Afp18NT20 | MPYSSESKEKETHSKETERD | 5% (R), 5% (D), 25% (E), 5% (H), 15% (K), 20% (S), 10% (T), 5% (Y) | 90 | -1,39 |
| *Serratia entomophila* | Afp17NT20 | MPTKTPQLQLAIEEFNKAIL | 5% (N), 10% (Q), 10% (E), 10% (K), 10% (T) | 45 | 1,981 |
| *Serratia fonticola* | *Sf* Tox20 | MPYSRESKEKEIHAKETERD | 10% (R), 25% (E),, 15% (K), 10% S, 5% (H), 5% (Y) | 70 | -1,163 |
| *Erwinia persicina* | *Ep* Tox20 | MPYFNELNEKETRSKETESG | 5% (R), 10% (N), 25% (E), 10% (K), 10% (S), 10% (T) | 70 | -2,109 |
| *Yersinia pekkanenii* | *Yp* Tox20 | MLYSSESKEKKTHSKETERD | 5% (R), 5% (D), 20% (E), 20% (K), 20% (S), 10% (T) | 80 | -2,15 |
| *Serratia ureilytica* | *Su* Tox20 | MPYFRESKEKDTHAKESKQD | 5% (R),10% (D), 15% (E), 5% (H), 20% (K), 10% (S), 5% (T), 5% (Y) | 75 | -0,904 |
| *Serratia marcescens* | *Sm* Tox20 | MPYSRESKEKDTHAKGSKQD | 5% (R), 10% (D), 10% (E), 5% (H), 20% (K), 15% (S), 5% (T), 5% (Y) | 75 | -0,789 |
| *Salmonella enterica* | *Se* Tox20 | MPYSSESKLKDTHLKEAESD | 10% (D), 15% (E), 10% (L), 15% (K), 20% (S), 5% (T), 5% (Y) | 80 | -1,76 |
| *Photorhabdus luminescens* | VC1_08520N20 hemopexin | MNISSYFFLNEENIRFNNQC | 5% (R), 25% (N), 5%(Q), 10% (E), 10% (S), 5% (Y) | 60 | -0,1579 |
| *Photorhabdus luminescens* | VC2_08635N20 hypothetical | MLSTEKHNKDTKHPRNREKK | 10% (R), 10%(N), 5% (D), 10%(E), 10%(H), 25%(K), 5%(S), 10%(T) | 85 | -2,164 |
| *Photorhabdus luminescens* | VC3_08730N20 hypothetical | MPNSKYSEKVNHSANGAEKC | 15%(N), 10%(E), 5%(H), 15%(K), 15%(S), 5%(Y) | 65 | -0,307 |
| *Photorhabdus luminescens* | VC4_08830N20 Toxin CyaA | MPRYSNSQRTPTQSTKNTRR | 20%(R), 10%(N), 10%(Q), 5%(K), 15%(S), 20%(T), 5% (Y) | 85 | -1,273 |
| *Photorhabdus luminescens* | VC5_12690N20 Cysteine prot. | MEHEYSEKEKPQKCPIQLRD | 5%(R), 5%(D), 10%(Q), 20%(E), 5%(H), 15%(K), 5%(S), 5%(Y) | 70 | -0,3316 |
| *Photorhabdus luminescens* | VC6_13210N20 hypothetical | MYDSKKKNSEPTTKKKFERS | 5%(R), 5%(N), 5%(D), 10%(E), 30%(K), 15% (S), 10%(T), 5%(Y) | 85 | -1,2533 |
| *Photorhabdus asymbiotica* | PAVC4_1PAU_RS13645 hypothetical | MPNKKYSENTHQGKKPLIKS | 10%(N), 5%(Q), 5%(E), 5%(H), 25%(K), 10%(S), 5%(T), 5%(Y) | 70 | -0,247 |
| *Photorhabdus asymbiotica* | PAVC4_2PAU_RS22355 hypothetical | MEREYSEKEKHKKHPIQLRD | 10%(R), 5%(D), 5%(Q), 20%(E), 10%(H), 20%(K), 5%(S), 5%(Y) | 80 | -1,2646 |
| *Photorhabdus asymbiotica* | PAVC4_3PAU_RS13655 hypothetical | MVHEYSINDRQKRHSFSSAN | 10%(R), 10%(N), 5%(D), 5%(Q), 5%(E), 5%(K), 20%(S), 5%(Y) | 65 | -1,886 |
| *Photorhabdus asymbiotica* | PAVC5_4PAU_RS16555 cytotoxicNF 1 | MLKYANPQTVATQRTKNTAK | 5%(R), 10%(N), 10%(Q), 15%(K), 20%(T), 5%(Y) | 65 | -0,8359 |
| *Photorhabdus asymbiotica* | PAU_RS10120 RHS repeat | MISTFDPAICAGTPTVTVLD | 10%(D), 5%(S), 20%(T), | 35 | -0,0745 |
| *Photorhabdus asymbiotica* | PAU_RS10125 YopT | MEREYNKKEKQKKSAIKLDD | 5%(R), 5%(N), 10%(D), 5%(Q), 15%(E), 30%(K), 5%(S), 5%(Y) | 80 | -1,242 |

**Table S6: Packing motif parameters for N-terminal peptides of CIS cargo**

Examples of packing motifs in novel and related CIS particles and their negative CC1,3(lag=2) values. Afp17NT20 is an experimentally confirmed motif that does not pack and has a high positive CC1,3(lag=2) value. The CC values were calculated over the available VaxiJen server (http://www.ddg-pharmfac.net/vaxijen/VaxiJen/VaxiJen.html) which offers ACC and CC calculations of proteins. Total polar amino acid (aa) content was calculated using the ProtParam server (https://web.expasy.org/protparam/).

| # | homologue to | genome code | location | full seq. identity % | strain | CDS | peptide | protein name | aa length |
|---|---|---|---|---|---|---|---|---|---|
| 1 | *Yr* Afp17NT20 | MT039173.1 | 96200:103054 | 61,8 | *Serratia proteamaculans strain AGR_4 plasmid unnamed1,* | ULG13700.1 | MPYASELKKLDKKPTENE | hypothetical protein | 2285 |
| 2 | *Yr* Afp17NT20 | CP009539.1 | 1748617:1754976 | 96,5 | *Yersinia ruckeri strain YRB, complete genome* | AJI94641.1 | MPYSNKSKKNEIRSEKSN | tcdA/TcdB catalytic glycosyltransferase domain protein | 2121 |
| 3 | Afp18NT20 | MT039196.1 | 91004:95170 | 39,1 | *Serratia proteamaculans strain AGR_1137 plasmid unnamed1* | ULG16493.1 | MPYSRESKEKETHPKETK | hypothetical protein | 1949 |
| 4 | *Ep* Tox20 | CP025698.1 | 1685283:1688714 | 34 | *Serratia marcescens strain SOLR4 chromosome* | AUO01772.1 | MPYSRESKEKDTHAKGS | hypothetical protein | 2166 |
| 5 | CyaANT20 | CP020335.1 | 1949579:1950583 | 88,5 | *Photorhabdus akhurstii strain 0813-124 phase II chromosome* | QXF33179.1 | MPRYSNSQRIPTQNSKNS | Insecticial toxin | 337 |
| 6 | CyaANT20 | CP020335.1 | 2002383:2003390 | 90 | *Photorhabdus akhurstii strain 0813-124 phase II chromosome* | QXF33220.1 | MPRYSNSQKIPTQGTKNT | Insecticial toxin | 336 |
| 7 | CyaANT20 | FM211060.1 | 69666:70673 | 79,9 | *Photorhabdus asymbiotica subsp. asymbiotica ATCC 43949 BAC clo* | CAR67759.1 | MPRYANYQINPKQNIKNS | Putative insecticidal toxin | 336 |
| 8 | CyaANT20 | FM162591.1 | 2407857:2408756 | 68,8 | *Photorhabdus asymbiotica ATCC43949 complete genome* | CAQ84235.1 | MFENDVGVRQQVKREFM | conserved hypothetical protein | 303 |

**Table S7.**

**Bioinformatic investigation of eCIS like regions in public databases**

Homologs to our experimentally validated toxins in these loci using tblasn with the same parameters, after which we extracted the 20 N-terminal amino acids, and homology reduced these with a 90% identity threshold. The search yielded 9 new potential NtSPs. Genome accession code (NCBI), location, bacterial strain, gene loci, peptide sequence, protein and length are highlighted.
